# Surveying the target tractability of cytokines with small molecules

**DOI:** 10.64898/2026.04.20.719718

**Authors:** Raavi, Sean P. Quinnell, Andrea Casiraghi, Becky Leifer, Benjamin Leu, Morgan Stilgenbauer, Fiona Wang, Haoyi Hou, Angela N. Koehler, Arturo J. Vegas

**Affiliations:** Koch Institute for Integrative Cancer Research, Massachusetts Institute of Technology, Cambridge, MA, 02139; Department of Biological Engineering, Massachusetts Institute of Technology, Cambridge, MA, 02139; The Broad Institute of MIT and Harvard, Cambridge, MA, 02142; Department of Chemistry, Boston University, Boston, MA 02115

## Abstract

Cytokines are key mediators of inflammation and are prominently involved in immune-mediated disorders, playing key roles in the pathogenesis of diseases such as rheumatoid arthritis, asthma, cancer, and systemic lupus erythematosus. Currently, cytokines are a challenging class of protein targets for traditional small-molecule drug discovery efforts. Biologic-based inhibitors have achieved clinical success, but the current suite of biologics therapies is limited, lack oral bioavailability, and have numerous side effects and compliance challenges. The development of small-molecule therapeutics is an attractive alternative that could further expand our therapeutic modulation of these targets. Here, we profiled a panel of 32 disease-relevant human cytokines to identify small-molecule ligands and inhibitors to survey their tractability for small-molecule modulation. Using a binding-first, small-molecule microarray-based approach we probed the binding preferences of each cytokine against a collection of 65,000 drug and lead-like compounds. We have identified 864 key chemical chemotypes that define structural motifs that bias for binding to specific cytokines. We further validated these chemotypes in a thermal denaturation sensitivity assay, resulting in 296 validated cytokine binders. We then prioritized three cytokines and established that novel, first-in-class inhibitors can be identified from these binders with potency ranging from single-digit to double-digit micromolar in reporter cellular assays. Boltz-2 predictions further delineated the binding landscape, underscoring how these inhibitors engage cytokine surfaces with defined structural complementarity. For the first time, our studies show that cytokines are indeed broadly amenable to small-molecule binding and inhibition with key insights into the chemical structures that can enable the inhibition of specific cytokines.

## INTRODUCTION

Cytokines are soluble, pleiotropic signaling proteins that mediate and balance the intricate communication network of the immune system and its responses.^1–4^ These diverse signaling molecules are produced by various immune cells, including T cells, B cells, macrophages, and dendritic cells.^5,6^ These cells use cytokines to signal in an autocrine, paracrine or endocrine manner to regulate and maintain immune homeostasis by immune cell activation, differentiation, proliferation and maturation, playing a crucial role in both innate and adaptive immunity.^3,7–12^ Dysregulation of cytokines can lead to a wide range of immune-mediated disorders, including chronic inflammation, autoimmune disorders, and cancer.^3,3,8,13^ For instance, dysregulated type 2 cytokines IL-4, IL-13 and IL-5 are hallmarks of chronic airway inflammation in asthma^14,15^, similarly, dysregulated IL-23 plays a critical upstream regulatory role in the psoriasis inflammatory cascade by regulating T_H_17 differentiation, leading to the downstream production of IL-17A and other proinflammatory cytokines.^16–20^

Cytokines exert their effects by binding to their cognate receptors on the cell surface through various binding modes, triggering downstream intracellular signaling cascades that modulate gene transcription. Depending on their structural families and receptor interactions, cytokines can activate multiple downstream signaling pathways, including the JAK/STAT, MAPK, NF-κB and ERK pathways.^21,22^ For instance, IL-13 and IL-4 both bind a type II receptor complex (comprising IL-14Rα1 and IL-13Rα1 subunits) and mediate signaling via JAK/STAT phosphorylation.^23^ Additionally, IL-2, IL-4, IL-7, IL-9, IL-15 and IL-21 utilize a common γc subunit (type I receptor), highlighting the interconnected nature of cytokine signaling.^24^ Interestingly, IL-23 and IL-12 not only share a receptor IL-12Rβ1 subunit for downstream STAT signaling but they also share p40 subunit underscoring their structural similarity.^25,26^ The IL-17 family consists of six cytokines (IL-17(A-F)) and five cell surface receptors (IL-17RA-E)^27,28^, and mediates signaling by recruiting Act1 which ubiquitinates and activates TRAF6 (TNF receptor-associated factor 6). TRAF6 activation causes IκBα phosphorylation, leading to its degradation and allowing NF-κB (p65 and p50) to translocate into the nucleus.^29,30^ Other downstream signaling pathways are also initiated as well including MAPK, C/EBP, AKT-dependent JAK2/STAT3 phosphorylation.^30–32^

These archetypical protein-protein interactions (PPIs) between cytokines and their cognate receptors have been previously targeted and clinically validated for immune-mediated disorders with various biologics primarily with therapeutic monoclonal antibodies (mAbs), given these PPIs are challenging to drug due to their characteristic flat and featureless binding surface, rendering this class of targets difficult to drug with specificity and potency.^33–36^ For instance, the current clinical standard for TNF-α is FDA approved mAbs, includes, infliximab for moderate-to-severe active rheumatoid arthritis^37^, and adalimumab for juvenile idiopathic arthritis^38^ with 30 biologics approved by FDA targeting either cytokine or their cognate receptors.^8,39–68^ However, monoclonal antibodies (mAbs) are primarily administered intravenously due to their large size and hydrophilicity, which limit absorption through other routes.^69,70^ Additionally, their poor gastrointestinal stability and non-specific clearance by the reticuloendothelial system further restrict alternative routes.^69,70^ mAbs may also induce immunogenic responses, both target-dependent and independent (immunogenicity), potentially impacting therapeutic efficacy.^39,71–74^ Maintaining these therapies presents additional challenges, including long systemic half-lives, limited tissue penetration, short tissue residence, and delayed peak concentrations. Although there is a growing preference for subcutaneous delivery (∼30%) due to patient acceptability, intramuscular and intralesional routes remain rare and highly indication-specific.^39,71–81^ While less specific than mAbs, small molecules offer advantages such as good bioavailability, tissue penetration, stability, rapid absorption, and metabolism, along with a lack of unwanted immunogenic responses.^39,75–77,82,83^ These characteristics, particularly the option for oral administration, make small molecules attractive drug candidates, ensuring easier maintenance with consistent dosing, enhanced compliance, and improved outcomes.^39,75–77,82,83^

Recent clinical advancements in small-molecule inhibitors targeting soluble cytokines are important therapeutic milestones. SAR441566, an allosteric small-molecule inhibitor of TNF-α, has progressed to a Phase II for moderate-to-severe rheumatoid arthritis (Clinical Trial Identifier: NCT06073093).

Similarly, Dice Therapeutics (now a subsidiary of Eli Lilly) completed a Phase IIb dose-ranging trial for its IL-17 inhibitor, DC-806, in moderate-to-severe plaque psoriasis (Clinical trial identifier: NCT05896527). These clinical advancements have been built on the efforts spanning last two decades leveraging a combinatorial approach of high throughput screening technologies, virtual screening, and biophysical & biochemical assays. Preclinical small molecule inhibitors for various other soluble cytokines have been identified as well including IL-1β, IL-18, IL-36C, MIF-1/2, C5, IL-2, IL-4, IL-15, IFNC.^77,84–110^ Despite extensive efforts to target soluble cytokines, a key challenge in the development of small-molecule cytokine inhibitors is a deficiency of known chemical matter that can be optimized for inhibiting their high-affinity protein-protein interactions (PPIs).^111,112^ Thus, highlighting the need for identification of novel chemical moieties to develop potent and selective drug candidates for these cytokines.

In this study, we profiled 32 human disease-relevant cytokines using high-throughput small-molecule microarray (SMM) screening with a library of 65,000 drug-like and lead-like compounds, leading to the identification of 2737 unique SMM positives. These positives with enriched chemotypes (864) were further validated in thermal denaturation sensitivity assay using either differential scanning fluorimetry (DSF) or nano-DSF depending on cytokine suitability, resulting in 281 confirmed cytokine binders. We prioritized three cytokines—**IL-17**, **IL-13**, and **IL-23**—for further evaluation of their binders in reporter cellular assays. This led to the discovery of novel, selective inhibitors for all three cytokines, with cellular potencies ranging from single digit to double-digit micromolar levels. We complemented these findings with Boltz-2 structural predictions, which identified specific cytokine interaction pockets that rationalize inhibitor engagement for cytokine modulation. These inhibitors provide a foundation for developing treatments for cytokine-specific immunological disorders and cancers. Additionally, this study serves as a starting point for demonstrating the broader tractability of these cytokines as drug targets.

## RESULTS

### Small-molecule microarrays screening of 32 human cytokines

Cytokines were selected for this study based on their known role in contributing to immune-mediated diseases and their commercial availability as poly His-or streptavidin-tagged proteins **(Table S1).** Small-molecule microarrays (SMMs) were utilized for high-throughput screening of this broad cytokine panel due to their ability to assess thousands of protein-ligand interactions simultaneously and their covalent immobilization of small molecules, which has been optimized to capture of a diverse set of functional groups. ^108,113–119^ Putative positives to recombinant poly-His or streptavidin tagged cytokines were identified by screening arrays consisting of a total of 65,000 covalently immobilized printed drug-and lead-like small molecules as well as controls as described previously **(Figure 2A and S1)**. ^108,113–120^ The screened library was evaluated for drug-likeness based on physicochemical properties and compared to ∼11K FDA-approved drugs. **Figure S2** shows the distribution of QED (quantitative estimate of drug-likeness, ranging from 0 = lowest to 1 = highest) for both datasets. The majority (39.2%) of the screened library falls within a QED range of 0.50–0.75. SMM assays encompassed data for four replicates for each small molecule and DMSO controls, evaluating ∼2.1 million ligand-protein interactions. Z-scores from the SMM data were computed as described previously^113,121,122^, with assay positives identified as those with a z-score atleast three standard deviations of the mean per slide. A total positivity rate of 0.5-0.8% was achieved per cytokine (11,837 total positives, **Figure 2B)**. Notably, SMM screening revealed a total of 2,737 unique positives across all 32 cytokines **(Figure 2C)**. These patterns of selectivity demonstrate that even closely related cytokines possess distinct small-molecule binding preferences, revealing previously unappreciated chemical vulnerabilities that can be exploited for designing highly selective inhibitors. For instance, we identified selective SMM positives for IL-1 family members—IL-1α, IL-1β, IL-18, and IL-33^123–125^. While IL-1α, IL-1β, and IL-33 belong to IL-1 and IL-33 subfamilies and share the co-receptor IL-1RAcP, IL-18 signals through a distinct receptor complex.^126,127^ All four cytokines, however, are members of the IL-1 family and are evolutionarily related, characterized by a conserved Toll/IL-1R (TIR) domain in their cytosolic regions.^128–131^ This domain, also found in Toll-like receptors (TLRs), underlies their well-established role in mediating innate immune responses.^128–131^ Similarly, selective positives were identified for cytokines such as IL-2, IL-7, and IL-21, which utilize a common γc receptor subunit that facilitates downstream signaling primarily through the JAK/STAT pathway.^24,132–135^ We also observed selective positives for IL-12 and IL-23, which share both the receptor subunit IL-12Rβ1 and a structural p40 subunit, highlighting their potential for novel, selective inhibitors aimed at modulating T_H_1/T_H_17 cell responses.^136–140^ In addition, selective SMM positives were detected for type 2 cytokines - IL-5, and IL-13, critical for regulating chronic airway inflammation and allergic responses through the production of T_H_2 cells or type 2 innate lymphoid cells.^14,15^ Lastly, we identified selective positives for clinically validated cytokine targets or their cognate receptors in rheumatoid arthritis, including TNF-α, IL-17, and IL-6.^37,38,44,57,64,136,141–157^ **(Figure 2C)**.

**Figure 1.**
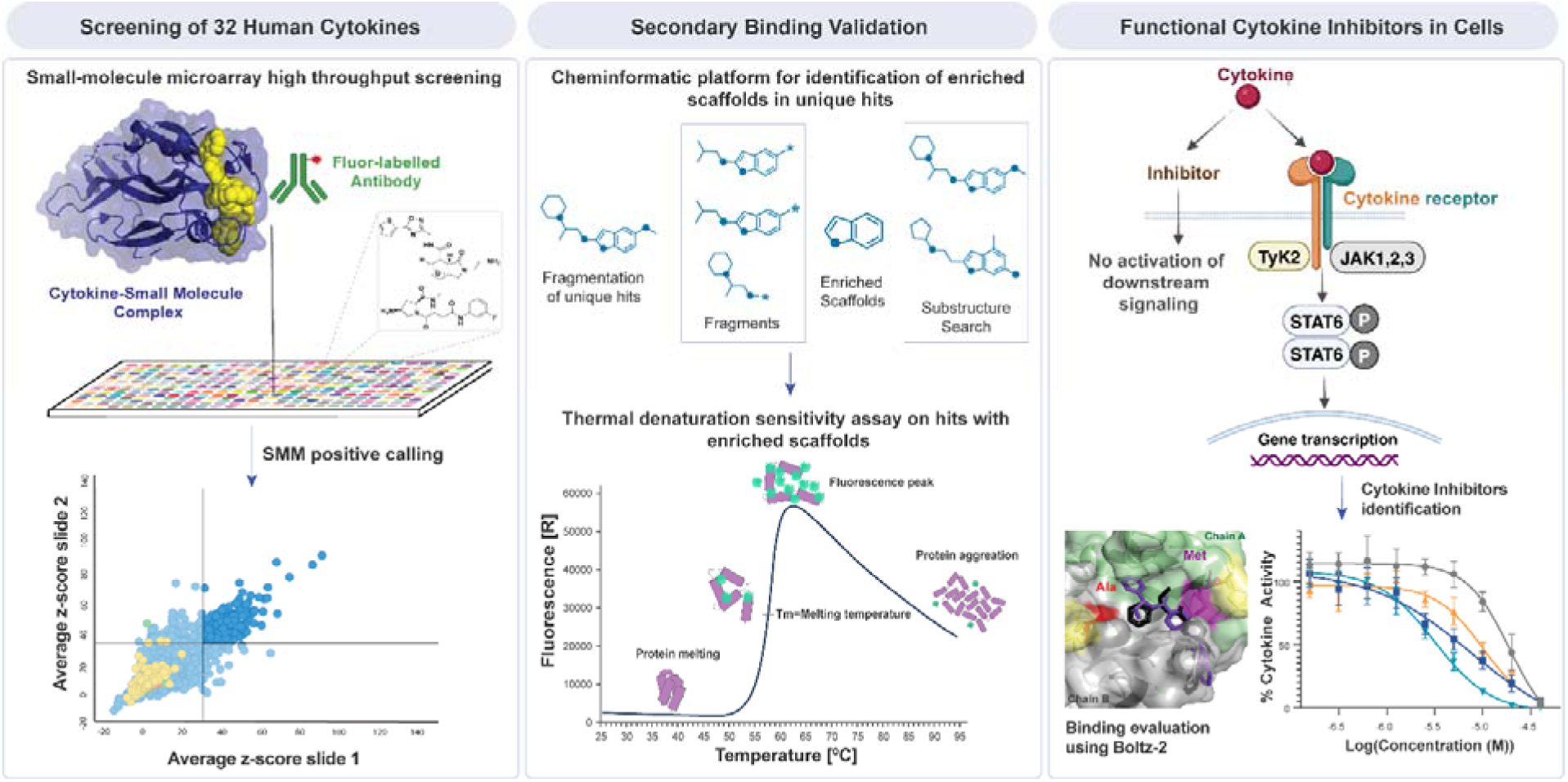
Workflow for surveying the tractability of soluble cytokines and identification of inhibitors for prioritized cytokines IL-17, IL-13 and IL-23. Thirty-two human cytokines were screened against a library of 65,000 drug-like and lead-like compounds using small-molecule microarrays (SMMs). SMM positive calling was performed using composite z-scores calculated by averaging per-feature z_i_-scores across slides, retaining only features with a z-score atleast three standard deviations of the mean per slide. Cytokine-specific positives were cross-referenced with an antibody counter-screen to remove non-specific binders. Unique SMM positives for each cytokine underwent fragmentation and scaffold-enrichment analysis to identify statistically enriched chemical motifs. Approximately 50 representative SMM positives per cytokine were then prioritized for secondary binding validation using a thermal denaturation sensitivity assay. For IL-17, IL-13, and IL-23, validated binders were assessed for functional activity in a HEK-Blue reporter assay at a single dose of 10 µM. Compounds showing ≥40% inhibition of cytokine signaling were advanced to dose-response studies to determine cellular potency. This multi-step workflow enabled the identification of novel IL-17 inhibitors and first-in-class inhibitors for IL-13 and IL-23, which were further evaluated for cytokine binding using Boltz-2, a deep-learning–based structure prediction model.

**Figure 2.**
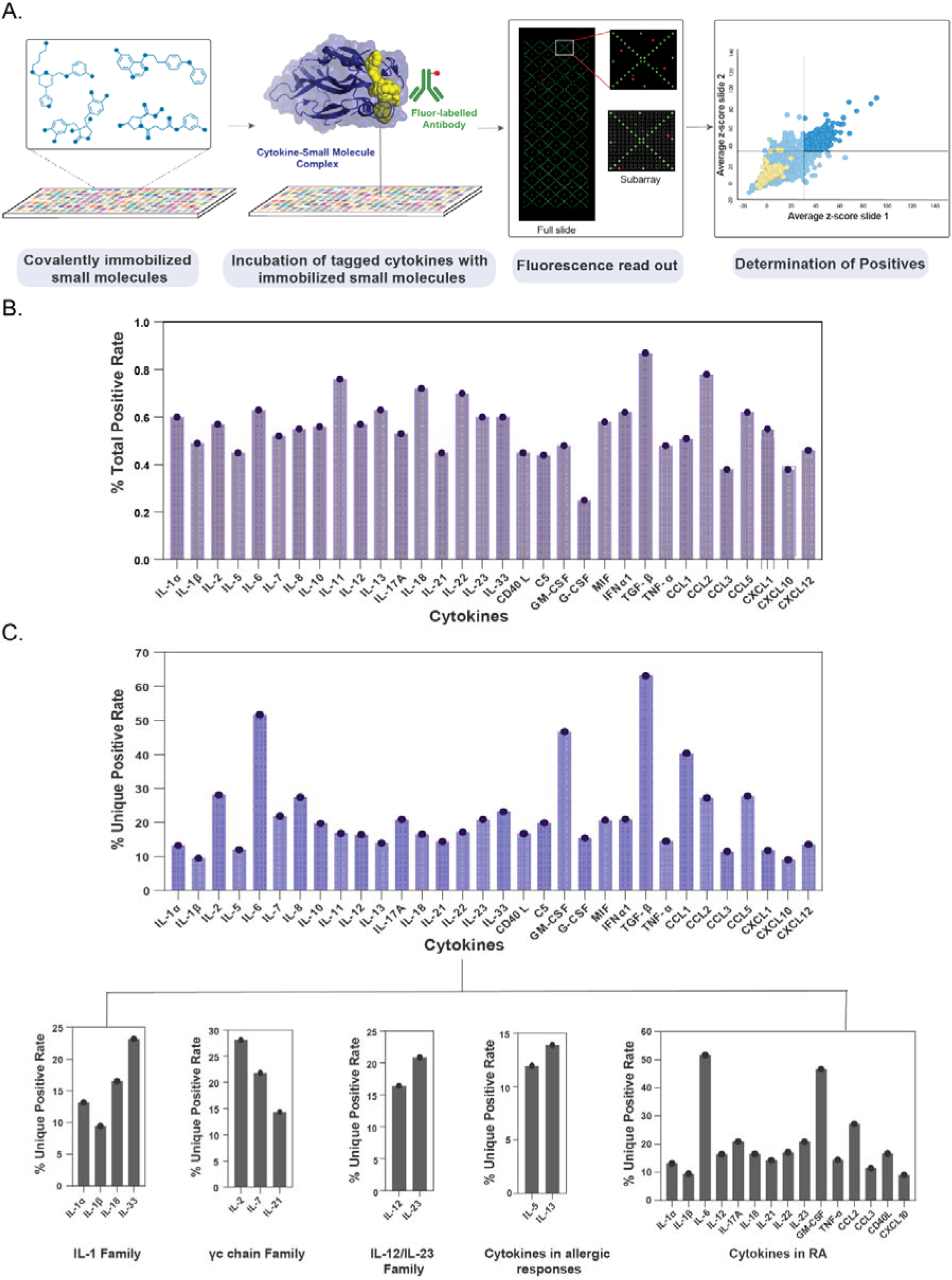
Small-molecule microarray screening of 33 cytokines. **(A)** Workflow for small-molecule microarray screening. **(B)** Total positive rate per cytokine, calculated as (total positives per cytokine / total printed features) * 100. **(C)** Unique positive rate per cytokine, calculated as (unique positives per cytokine / total positives per cytokine) *100, further shown in separate panels based on homology, and disease relevance.

### Selection of unique positives for secondary binding validation using thermal denaturation sensitivity assays

Unique positives for each cytokine were analyzed further for scaffold enrichment analysis. Prior to the enrichment analysis, all SMM positives were filtered for pan-assay interference compounds (PAINS).^158^ Molecular substructures were then calculated for each SMM positives list using BRICS^159^, RECAP^160^ and Bemis-Murcko^161^ algorithms which were then compared with the full library to determine if a substructure is significantly enriched in the positive set. **(Figure 3A)**. 606 uniquely enriched substructures were identified in 2,737 unique compounds across the 32 cytokines. Based on the enriched substructures and commercial availability of SMM positives consisting of these substructures, ∼50 unique compounds were chosen per cytokine for further secondary binding validation using thermal denaturation sensitivity assays (DSF/thermal shift). These experiments covered 26 cytokines (procurement costs prohibited pursuing the other 6 cytokines), screening 864 compounds at 2 µM in 0.1% DMSO (duplicates), confirming 296 cytokine binders with a ΔT_m_ cut-off >0.5°C and an average translation rate of 32.5%. Notably, C5 (100%), CD40L (97%), TNF-α (97%), IL-5 (86%), IL-6 (73%), IL-33 (55%), IL-12 (71%), and TGF-β (39%) exhibited significant translation rates ranging from 39% to 100% **(Figure 3B and Table S3)**. For IL-17, we observed substructure translation across multiple chemotypes. Sulfonyl-containing scaffolds showed 2 of 8 SMM positives validated as binders, and benzylamine-containing scaffolds similarly demonstrated translation, with 2 of 4 SMM positives validated. Carboxamide-derived macrocycles showed complete translation (2 of 2 validated), and additional amide-linked macrocyclic scaffolds also performed well (5 of 6 validated). Among heteroaromatic motifs, 1 of 3 pyrazole-containing SMM positives validated as a binder, while isoquinoline derivatives yielded 1 validated binder out of 2 SMM positives. For piperidinyl-substituted pyrazoles, 1 of 2 SMM positives validated. Pyridylalkyl scaffolds bearing cyclic tertiary amines showed full translation (2 of 2 validated), and finally, 1 of 2 SMM positives with a pyrrolidine–piperazine diamine scaffold validated as a binder.

**Figure 3.**
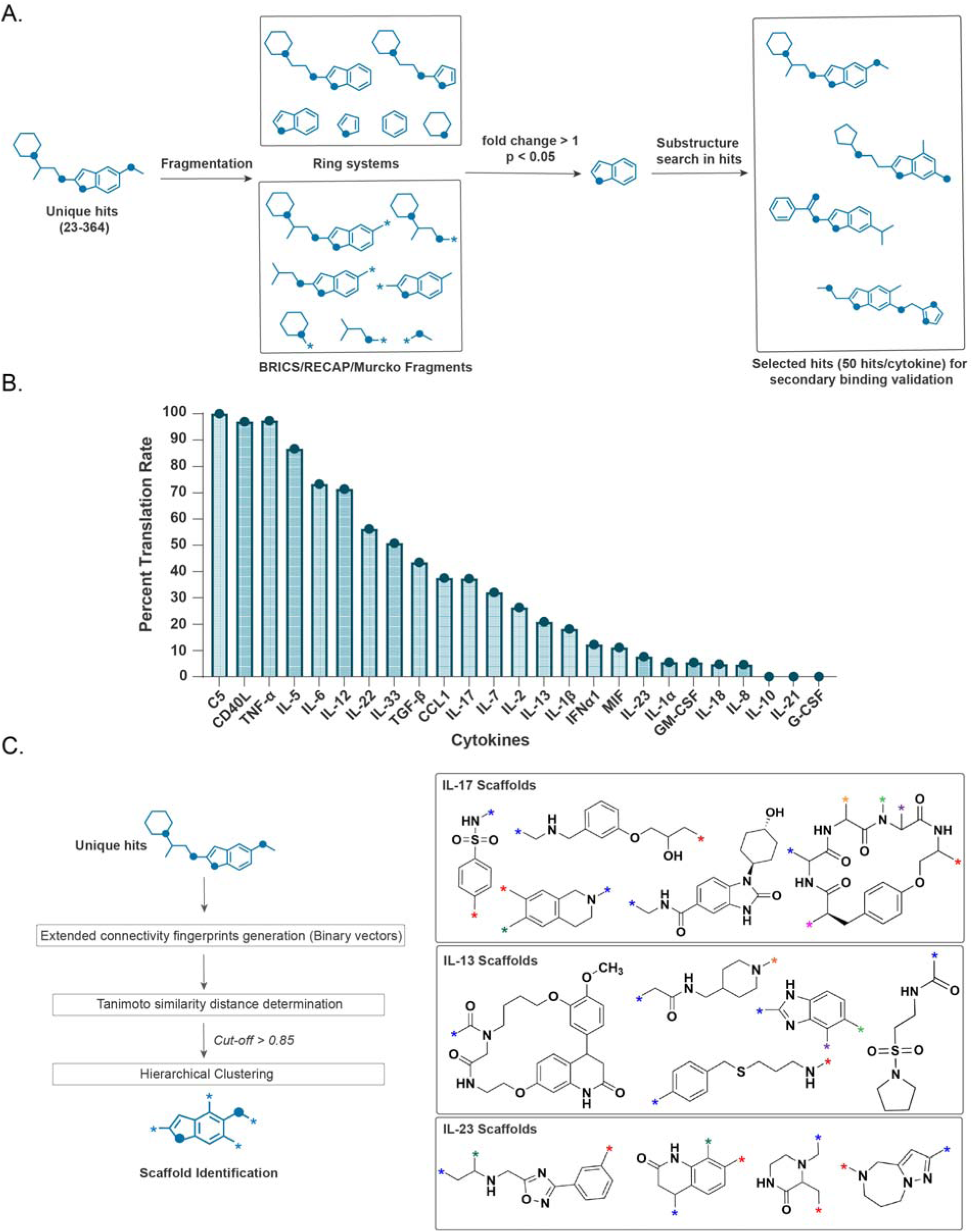
Unique positives with enriched scaffolds determination and further validation in thermal denaturation sensitivity assay. **(A)** Cheminformatic platform schematic for identification of enriched scaffolds from unique positives and substructure search in individual cytokine unique positive list. **(B)** Secondary binding validation of 25 cytokines with unique positives using a thermal denaturation sensitivity assay, focusing on enriched scaffolds. Percent translation rate was calculated as (validated binders per cytokine / total unique positives with enriched scaffolds per cytokine screened) *100. **(C)** Hierarchical clustering of topological fingerprints generated from unique SMM positives for prioritized cytokines IL-17, IL-13 and IL-23 for scaffold identification.

Following secondary binding validation, we next sought to identify which cytokines warranted deeper investigation through cell-based functional assays. To avoid selecting targets solely based on the number of validated binders, we instead grounded our choices by considering their relevance to three categories of biological relevance: 1) cytokines related by homology and that share receptor components or signaling logic, such as the IL-2 family including IL-4, IL-7, IL-9 and IL 21, which signal through the common γ-chain receptor,^23^ 2) disease-relevant cytokines with established roles in immune mediated disorders and validated clinically, including IL-17, IL-23, IL-6, IL-1α, IL-1β, and TNF-α,^162^ and 3) cytokines that play key immunoregulatory roles, such as IL-12 and IL-23, which both signal through IL-12Rβ1, share the p19 subunit, and shape T_H_1 versus T_H_17 differentiation.^163^ We also considered the IL-4 and IL-13 axis, which signals through the shared IL 4Rα chain, given its central role in type 2 inflammation and its importance in allergic and fibrotic disease.^23,164–166^

Integrating these criteria with practical considerations, including commercial trends and the availability of robust reporter cellular assays, led us to prioritize IL-17, IL-13, and IL-23 for downstream studies. IL-17 and IL-23 are key mediators of T_H_17 driven inflammation^16,167–169^ and are clinically validated therapeutic targets,^46,47,58,170^ providing benchmarks for assessing small molecule tractability. IL-13 represents a distinct and clinically important cytokine axis, functioning together with IL-4 through the shared IL-4Rα complex in type 2 inflammatory signaling.^23,164–166^ Of these three cytokines, only IL-17 has small-molecule inhibitors reported preclinically and in clinical trials, enabling us to compare our structures with those derived from distinct efforts. These cytokines allowed us to evaluate whether small molecule inhibitors can engage cytokines across mechanistically divergent but therapeutically validated pathways. Together, these considerations provide a coherent biological and translational rationale for prioritizing **IL-17**, **IL-13**, and **IL-23** for further studies.

Additionally, we aimed to identify scaffolds associated with these cytokines and correlate them with enriched substructures using the KNIME^171^ platform. To achieve this, topological fingerprints^172^ were generated from unique SMM positives for each of the three prioritized cytokines. Pairwise similarities were computed using Tanimoto distance^173^, and hierarchical clustering of these distance matrices revealed distinct scaffolds for each cytokine **(Figure 3C)**. For IL-17, we observed an enrichment of macrocyclic scaffolds, along with derivatives containing phenylsulfonamide, isoquinoline, methylaminomethyl-phenoxy-propanal, and hydroxycyclohexyl dihydrooxobenzoimidazole-carboxamide **(Figure 3C)**. For IL-13, identified scaffolds included derivatives such as a macrocycle, benzoimidazole, pyrrolidine-sulfonylethylaminomethanone, benzylthio-propylamine, and piperidinylmethyl-acetamide **(Figure 3C)**. In the case of IL-23, clustering revealed scaffolds encompassing derivatives of phenyl-oxadiazolylmethyl-ethanamine, dimethyl-piperazinone, dihydro-quinolinone, and dihydro-pyrazolo-diazepine **(Figure 3C)**. For all three cytokines, this hierarchical clustering highlighted scaffolds enriched with distinct substructures.

### Cell-based inhibitory activity determination of validated cytokine binders in HEK-Blue reporter cell line assays

The HEK-Blue IL-17 reporter cell line has been previously utilized for screening inhibitors of IL-17 activity.^30,174–178^ This genetically engineered cell line expresses the heterodimeric receptor IL-17 receptor (IL-17RA/IL-17RC) and IL-17 stimulation leads to secretion of embryonic alkaline phosphatase production (SEAP) in a NF-ĸB and AP-1-mediated manner **(Figure 4A)**. Thus, IL-17 stimulated SEAP production can be used to quantify inhibition. The 24 validated binders from thermal shift analysis **(Figure 3C)** were evaluated for preliminary inhibition in the HEK Blue IL-17 reporter cell line at a 10 µM concentration (4 replicates). Among these, eight IL-17 binders exhibited >40% inhibition at 10 µM **(Figure 4B and S3A, Table S4)**. These compounds were further evaluated for full dose-response in HEK-Blue IL-17 reporter cell line (repeated twice with n = 4), compounds **9** and **24** demonstrated a single digit micromolar potency of 6.43 µM and 3.16 µM respectively. Furthermore, compounds **22** (macrocycle) and **5** exhibited a potency of 20.9 µM and 29.9 µM **(Figure 4C and S3A)**. These inhibitors structurally span three distinct chemotypes: 1) **5** and **9** are compact heteroaromatic ligands containing amide or sulfonamide linkages that position polar groups around rigid aromatic cores, 2) **22** is a large, heteroatom-rich macrocycle with multiple amides and embedded ring systems that provide extensive three-dimensional surface coverage, and 3) **24** is an extended, non-macrocyclic multi-ring scaffold bearing several amide and heterocyclic features that afford a flexible yet directional interaction profile with IL-17. By comparison, current IL-17 small-molecule inhibitors in clinical trials (Clinical trial identifiers: NCT05896527, NCT06311656) from Dice Alpha Inc. (now a subsidiary of Eli Lilly) comprise carboxamide derivatives characterized by linear, non-macrocyclic polyamide frameworks with elongated backbones incorporating aromatic rings and bulky alicyclic substituents.^179,180^

**Figure 4.**
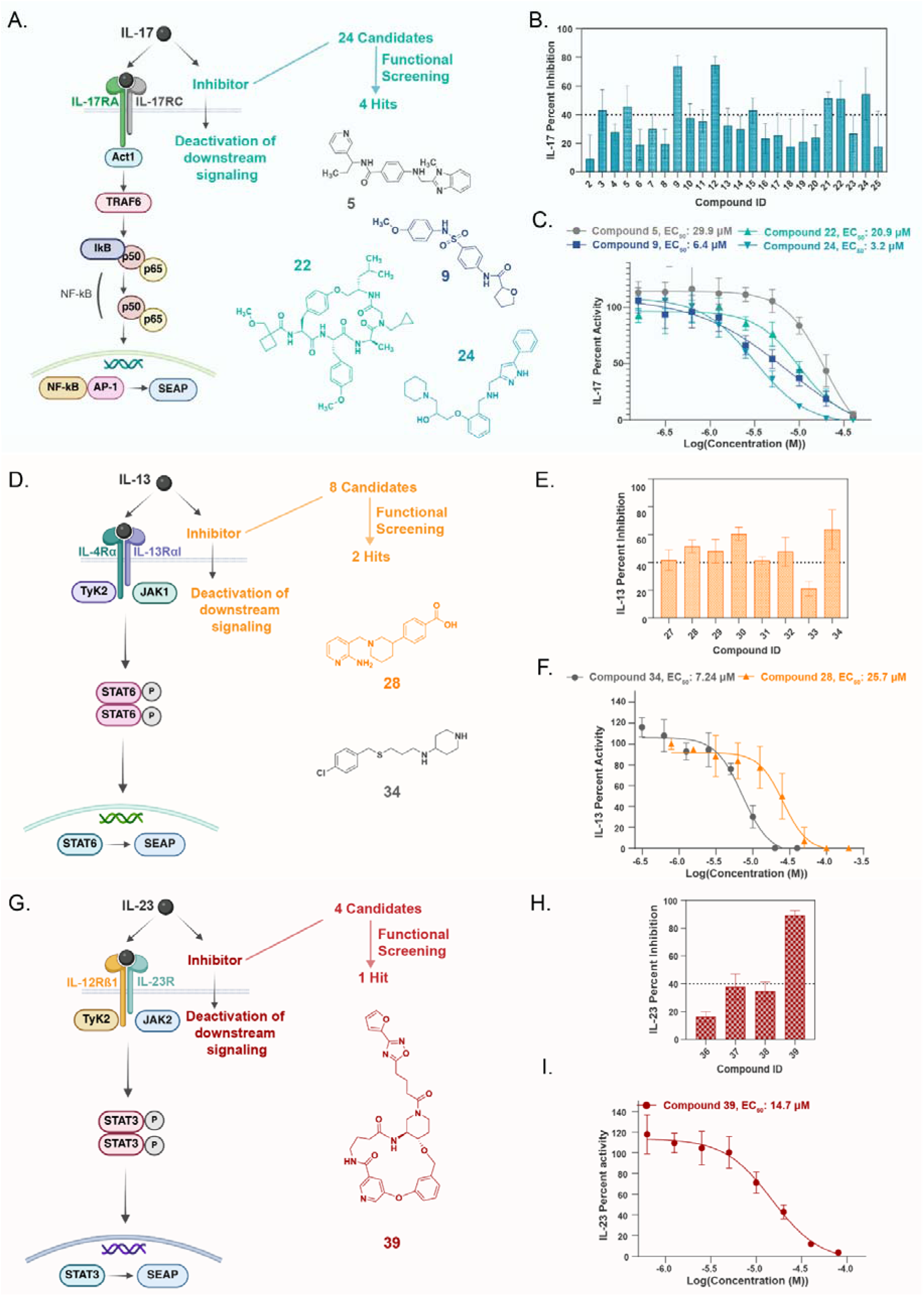
Evaluation of validated binders of prioritized cytokines IL-17, IL-13 and IL-23 in HEK-Blue reporter assays. **(A)** IL-17 signaling pathway in HEK-Blue IL-17 reporter assay. **(B)** Preliminary screening of 24 validated binders for IL-17 inhibition in HEK-Blue IL-17 reporter assay at 10 μM. **(C)** Dose-response evaluation of IL-17 inhibitors (**5**, **9**, **22**, **24**,) in HEK-Blue IL-17 reporter assay. **(D)** IL-13 signaling pathway in HEK-Blue IL-13 reporter assay. **(E)** Preliminary screening of 8 validated binders for IL-13 inhibition in HEK-Blue IL-13 reporter assay at 10 μM. **(F)** Dose-response evaluation of IL-13 inhibitors (**28**, **34**) in HEK-Blue IL-13 reporter assay. **(G)** IL-23 signaling pathway in HEK-Blue IL-23 reporter assay. **(H)** Preliminary screening of 4 validated binders for IL-23 inhibition in HEK Blue IL-23 reporter assay at 10 μM. **(I)** Dose-response evaluation of IL-23 inhibitor (**39**) in HEK-Blue IL-23 reporter

Similarly, HEK-Blue IL-4/IL-13 reporter cell line has been well established for screening antibodies^181,182^ and have also been to screen small molecule IL-4 inhibitors^108^. IL-13 binding to genetically engineered type II receptor in HEK Blue IL-4/IL-13 reporter cell line activates JAK1/TyK2 which facilitates STAT6 dimerization and phosphorylation, thus inducing SEAP production **(Figure 4D)**. From the 8 validated IL-13 binders from thermal shift analysis **(Figure 3C),** 7 compounds showed >40% inhibition at 10 µM in HEK Blue IL-4/IL-13 reporter cell line (4 replicates) **(Figure 4E and S3B, Table S5)**. Dose-responses were then determined for these 7 compounds, where compounds **28** and **34** exhibited the highest potencies of 25.7 µM and 7.24 µM potencies, respectively (repeated twice with n = 4) **(Figure 4F and S3B)**. Structurally, the IL-13 inhibitors fall into two distinct chemotypes: inhibitor **28** features a rigid fused heteroaromatic core linked to a para-hydroxybenzoic acid, creating an extended π-system with directional hydrogen-bonding potential, whereas inhibitor **34** is a smaller, flexible thioether-linked scaffold with a chlorophenyl ring and terminal secondary amine, representing a minimalist and adaptable framework for IL-13 engagement.

For IL-23, 4 validated binders from thermal shift were taken forward for preliminary inhibition **(Figure 3C)**. In the HEK-Blue IL-23 cell line, the IL-23:IL-23 receptor complex induces downstream SEAP production via JAK2/TyK2 mediated STAT3 dimerization and phosphorylation^183–186^**(Figure 4G)**. Out of the 4 validated binders, two compounds **37** and **39** demonstrated >40% inhibition at 10 µM (4 replicates). **(Figure 4H and S3C, Table S6)**. Upon dose-response evaluation, only **39** exhibited a potency of 14.7 µM in HEK-Blue IL-23 reporter cell line (repeated twice, n =4) **(Figure 4I and S4C)**. Structurally, inhibitor **39** is a heteroatom-rich macrocycle composed of ether linkages, tertiary amines, and pyridyl-containing aromatic units, with a single urea moiety extending from the ring. This arrangement creates a flexible, three-dimensional scaffold with multiple polar and aromatic features positioned along its periphery, enabling broad surface engagement with IL-23.

### Computational Binding Site Mapping of IL-17, IL-13, and IL-23 Inhibitors

To explore potential binding modes of IL-17, IL-13, and IL-23 inhibitors, we employed **Boltz-2**,^187^ a deep learning–based framework for biomolecular interaction prediction. We selected Boltz-2 because cytokine–small-molecule complexes are difficult to resolve by X-ray crystallography due to their conformational plasticity, and modeling therefore provides a practical route to generate binding-mode predictions that inform mechanistic interpretation and SAR. Boltz-2 is trained on all Protein Data Bank (PDB) structures released prior to June 1, 2023, with evaluation performed on structures deposited between June 1, 2023 and January 1, 2024^187^.

### Boltz-2 predicts IL-17 inhibitor engagement at the C-terminal binding site previously identified by AbbVie, with the macrocyclic inhibitor stabilizing the IL-17 dimer interface

As a qualitative check, we examined whether Boltz-2 recapitulates previously reported IL-17 binding poses. Boltz-2 generated poses for known IL-17 inhibitors (2016–2022), including macrocyclic antagonists from Pfizer and Eli Lilly that engage the IL-17 dimer interface, Pfizer’s HAP linear peptide, and AbbVie’s recently reported small molecule that binds the IL-17 C-terminal region.^188–192^ We anticipated the model to perform well in this task, given that these structures were part of the training set for Boltz-2, but would give us a sense of performance with an “easy” task. The predicted ligand orientations were broadly consistent with their corresponding PDB co-crystal structures **(Figures S4–S7)**, providing confidence that the model captures the general features of IL-17 ligand recognition.

Building on this validation, we next analyzed our discovered IL-17 inhibitors (compounds **5**, **9**, **22**, and **24**). Boltz-2 predicted that compounds **5**, **9**, and **24** occupy the C-terminal site of IL-17 with confidence scores of 0.839, 0.799, and 0.822, respectively **(Figures S9A, 5C, and S11A; Table S2)**. In contrast, compound **22**, a macrocyclic analog, was predicted to bind at the dimer interface with a confidence score of 0.825, the same region targeted by Pfizer and Eli Lilly macrocyclic antagonists, albeit from opposite directions **(Figure S10A, S4 and S5)**. To contextualize and add confidence in these predictions, we compared binding hypotheses for a SMM-negative compound (non-positive), and a validated binder (non-inhibitor). As expected, the SMM-negative compound **1** showed no engagement with known IL-17 binding pockets and minimal surface complementarity **(Figure 5A)**, while the binder **2** engaged the dimer interface in close proximity to Leu97(B) and Tyr62(A) **(Figure 5B)**. These results illustrate how Boltz-2 can generate structurally plausible poses that align with trends from our biochemical data, while recognizing that the method provides qualitative insights.

**Figure 5.**
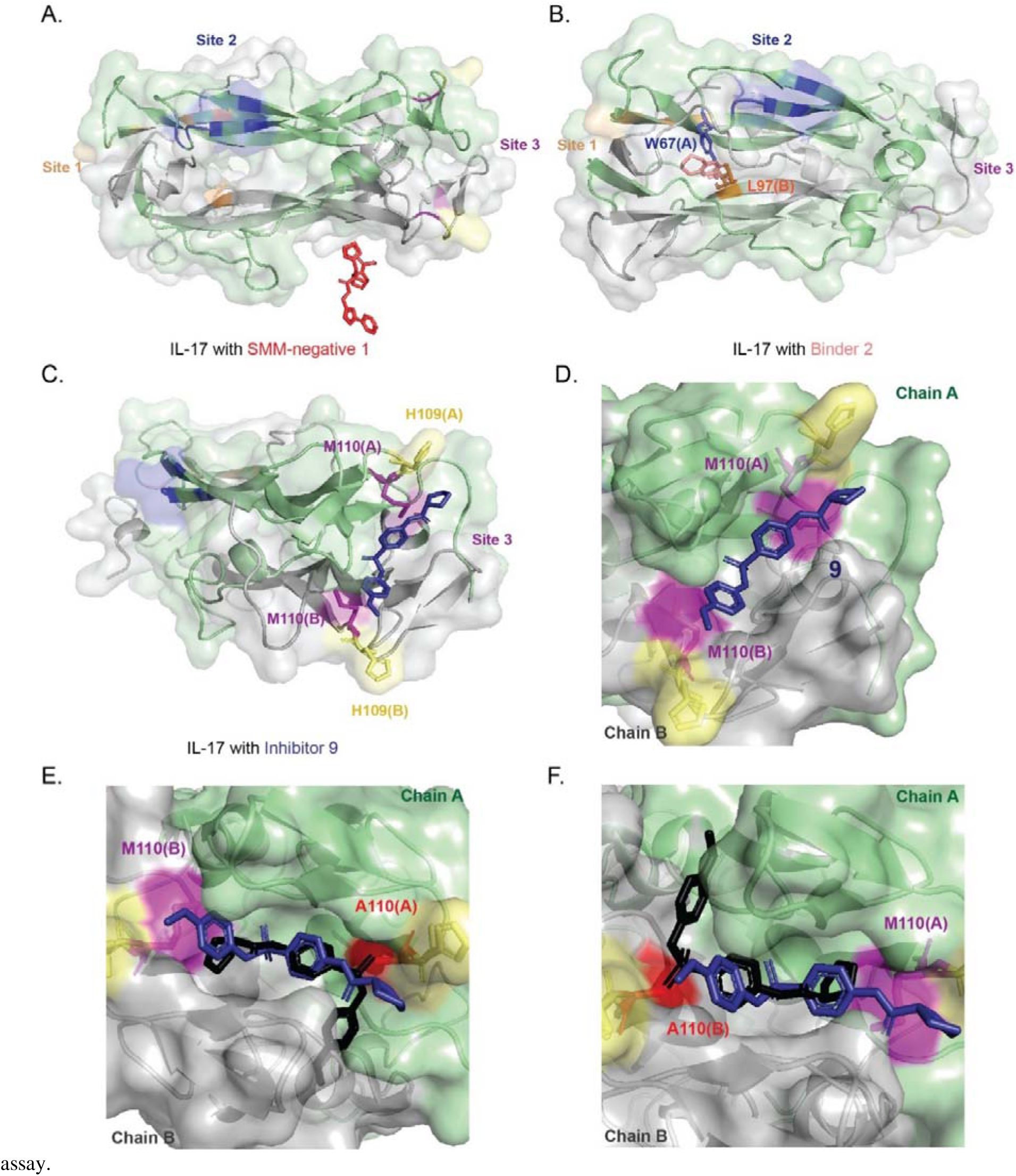
Boltz-2 prediction of IL-17 inhibitor 9 binding at a previously unrecognized C-terminal pocket. **(A)** SMM-negative compound 1 (red) shows minimal engagement with IL-17, remaining at a solvent-exposed surface region. **(B)** Binder 2 (light pink) engages residues within site 2. **(C)** Inhibitor **9** (dark blue) occupies a novel C-terminal binding pocket on IL-17, positioned between residues Met110 and His109 from both chains A and B. **(D)** Close-up of the C-terminal pocket showing Met110(A/B) (magenta) and His109(A/B) (yellow) forming the hydrophobic–aromatic cradle that accommodates inhibitor **9**. **(E–F)** *In silico* alanine-scanning mutagenesis using Boltz-2 demonstrates the functional relevance of this pocket: substitution of Met110(A) **(E)** or Met110(B) **(F)** with alanine (red) collapses the hydrophobic cleft and disrupts the predicted binding pose of inhibitor **9**. The black conformation represents the displaced pose adopted by inhibitor **9** upon mutation, highlighting the loss of key Met110-mediated contacts in stabilizing ligand engagement at the IL-17 C-terminus.

Previously, an NMR-based fragment screen of IL-17 identified a small hydrophobic pocket near the C-terminus, lined by Methionine residues from both chain A and chain B, capable of accommodating fragment-sized ligands.^193^ To assess whether these methionine residues contribute to the binding of our in-house inhibitors, we combined Boltz-2 predictions with *in silico* site-directed mutagenesis^194–196^, substituting each methionine with alanine and re-evaluating the binding of compounds **5**, **9**, and **24**. Boltz-2 suggests that the C-terminal pocket is shaped largely by Met110 on both IL-17 chains, and that these residues differentially influence how compact ligands orient within the cleft. For inhibitor **5**, mutation of Met110(A) caused a flipped, with its terminal aromatic ring repositioned toward the solvent-exposed region **(Figure S9B–C)**, while mutation of Met110(B) produced a lateral shift that maintained partial burial **(Figure S9D)**. These changes indicate that Met110(A) acts as the primary hydrophobic anchor, with Met110(B) fine-tuning pocket geometry. Inhibitor **9** showed a similar dependence on this methionine pair. Mutation of Met110(A) inverted its orientation, repositioning the sulfonamide-aryl unit toward solvent **(Figure 5E)**, whereas mutation of Met110(B) displaced the ligand upward within the cleft and reduced contact with both residues **(Figure 5F)**. Together, these results suggest that compact heteroaromatic scaffolds rely on coordinated stabilization from both Met110 residues to maintain a productive binding orientation.

For the inhibitor **22**, Boltz-2 predicts engagement of the IL-17 dimer interface rather than the C-terminal pocket **(Figure S10A–B)**. We selected Trp67(A) and Leu97(B) for mutation because both have been identified as key site-2 anchors by Pfizer and Eli Lilly IL-17 macrocycles, and Boltz-2 likewise predicted them to be the principal residues stabilizing inhibitor **22** at the dimer interface.^188–191^ When Trp67(A) was mutated, the macrocycle shifted toward chain B and lost its π-stacking interaction on chain A **(Figure S10C)**. In contrast, mutation of Leu97(B) caused the ligand to shift upward toward chain A (Figure S10D), reducing packing on chain B while partially restoring aromatic stacking beneath Trp67(A). These distinct shifts indicate that Trp67(A) and Leu97(B) contribute complementary anchoring roles, jointly maintaining the interfacial binding mode of inhibitor **22**.

For inhibitor **24**, Boltz-2 predicts engagement of the C-terminal pocket through an extended hydrophobic interface spanning Met110 on both chains **(Figure S11A–B)**. Mutation of Met110(A) caused the ligand to shift downward and inward toward chain B, reducing contact with the upper hydrophobic wall **(Figure S11C)**. In contrast, mutation of Met110(B) induced a flip of the distal aromatic–imidazole region toward chain B, altering the alignment of the phenyl–amide linkage **(Figure S11D)**. These shifts suggest that Met110(A) and Met110(B) provide complementary anchoring roles, and that the flexible multi-ring architecture of **24** allows it to accommodate changes in pocket shape while maintaining engagement within the same overall site.

### Boltz-2 predicts distinct IL-13 monomeric binding site engaged by first-in-class inhibitors through electrostatic interaction networks

A similar approach was applied to predict the binding interactions of IL-13 inhibitors, compounds **28** and **34**. IL-13 is composed of four α-helices and two β-strands.^197,198^ While there are no previously reported small-molecule IL-13 inhibitors, structural data of IL-13 bound to tralokinumab indicate interactions between helices A and D.^199^ Boltz-2 predicted that compound **28** interacts with helix D with a confidence score of 0.819, whereas compound **34** interacts between helices A and C with a confidence score of 0.729 **(Figure 6C, S12)**. We also analyzed SMM-negative **28** and binder **27** binding with IL-13, compound **28** doesn’t show any binding to IL-13 whereas compound 27 binds to solvent exposed region of IL-13 **(Figure 6A, 6B)**.

**Figure 6.**
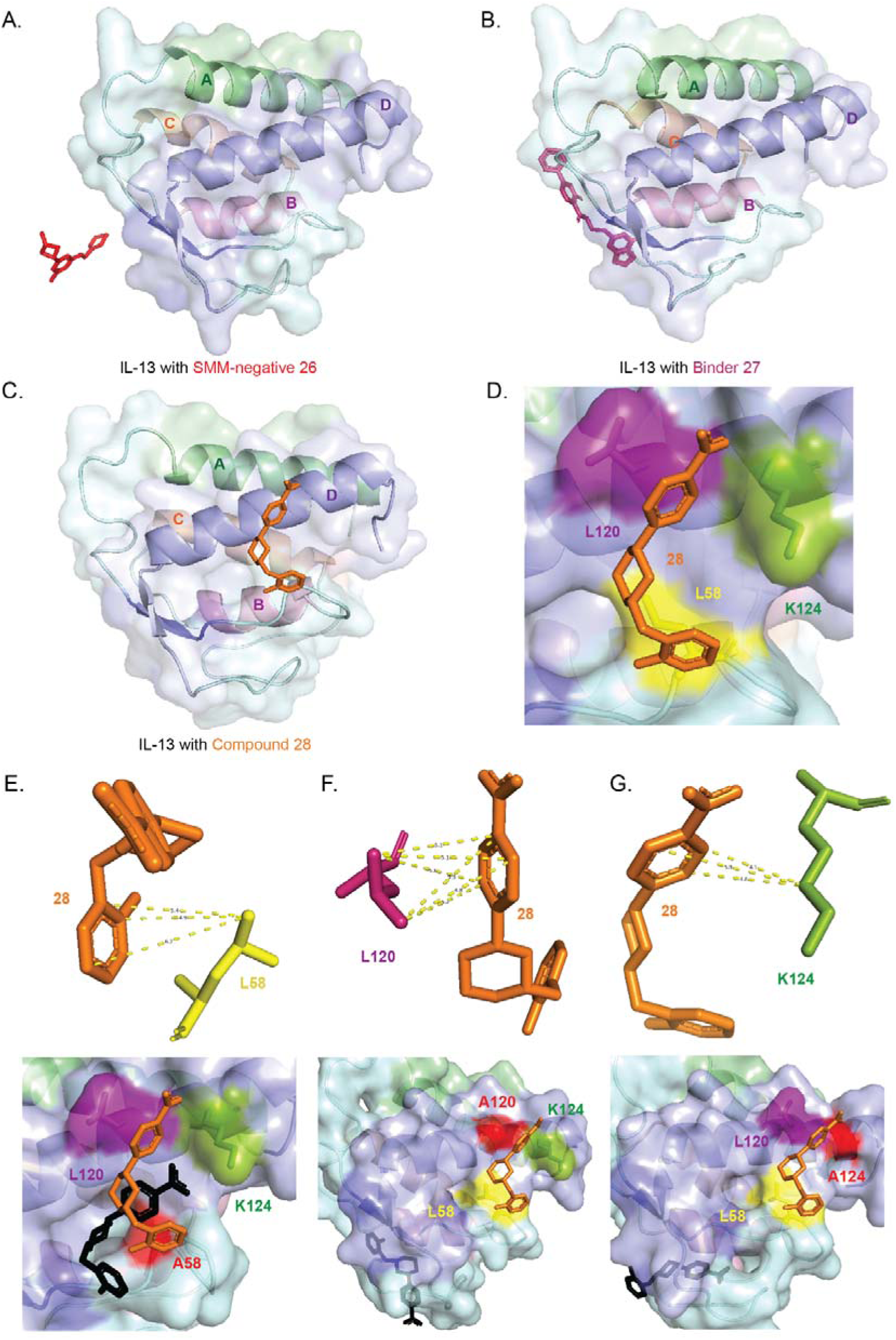
Boltz-2 prediction of IL-13 inhibitor 28 binding and *in silico* mutagenesis of the helix D pocket. (A) SMM-negative compound 26 (red) shows no detectable engagement with IL-13, remaining fully solvent-exposed. **(B)** Binder 27 (magenta) interacts only weakly along the IL-13 surface without accessing a defined pocket. **(C)** Inhibitor **28** (orange) binds along helix D, extending downward into a shallow hydrophobic region. **(D)** Close-up view of the binding environment highlighting L58 (yellow), L120 (magenta), and K124 (green) forming the core triad surrounding inhibitor 28. **(E–G)** *In silico* alanine-scanning mutagenesis using Boltz-2 reveals the importance of this triad for maintaining the IL-13 binding site architecture. Substitutions of L58, L120, or K124 with alanine (red) disrupts local packing and destabilizes the predicted pose of inhibitor 28. The black conformation represents the displaced pose adopted by inhibitor **28** upon mutation, emphasizing how each residue contributes uniquely to anchoring and orienting the ligand within the helix-D pocket.

To further validate these IL-13 binding interactions, we identified three key residues for inhibitor **28**, Leu58, Leu120, and Lys124 and performed *in silico* alanine-scanning mutagenesis followed by Boltz-2 prediction for each mutant individually. For inhibitor **28**, Leu58 forms a set of weak edge contacts with the ligand, with distances of 4.9–6.2 Å between the Leu58 side-chain and the lower aromatic ring **(Figure 6E)**. These interactions are modest but help position **28** within the binding region. Leu120 packs along the upper aromatic ring through a modest alkyl–π interaction, stabilizing the planar scaffold of **28** and contributing to the shape complementarity of the cleft **(Figure 6F)**. Lys124 lies along the opposite side of the ligand, where its positively charged side chain forms weak cation–π or electrostatic interactions with the upper aromatic ring of **28 (Figure 6G)**. Alanine substitution of Leu58 removes the hydrophobic base beneath **28**, causing the ligand to settle deeper into the cavity **(Figure 6E)**. Substituting Leu120 the key packing interaction that stabilizes the upper aromatic ring, causing **28** to disengage from its original site and relocate to an alternate surface on IL-13 **(Figure 6F)**. Substitution of Lys124 has a similar effect: removing its weak cation–π/electrostatic contribution leads to a fully displaced ligand pose, with **28** no longer occupying the original binding cleft **(Figure 6G)**. Collectively, these residues uphold the electrostatic landscape that allows compound **28** to adopt its stabilized orientation on IL-13.

Boltz-2 places inhibitor **34** within a shallow groove between helices A and C, where it engages a mixed polar–hydrophobic surface shaped by Glu31, Glu34, Lys81, Arg84, and Met85. Glu31 and Glu34 flank the upper groove and make weak polar contacts that help orient the heterocycle and aromatic core **(Figure S12C–D)**. Lys81 provides additional polar stabilization along the mid-groove **(Figure S12E)**, while Arg84 associates with the distal aromatic ring in the lower cavity **(Figure S12F)**. Met85 forms the hydrophobic floor beneath the phenyl ring **(Figure S12G)**, supporting the placement of the ligand across the groove.

*In silico* mutational analysis shows that several of these residues are essential for retaining the ligand in the groove at all. Substitution of Glu31 or Glu34 removes key polar constraints and causes inhibitor 34 to abandon its original binding site entirely, relocating to a distinct surface region **(Figure S12C–D)**. Loss of Lys81 similarly allows the ligand to collapse out of the groove, settling along the cavity floor near Met85 and Arg84 **(Figure S12E)**. Removal of Arg84 eliminates the anchoring element at the base of the pocket, producing a complete relocation of **34** away from the central **groove (Figure S12F)**. Only Met85 mutation preserves partial occupancy of the groove, although the ligand slides laterally across the lower cavity **(Figure S12G)**. Together, these results indicate that inhibitor **34** is not simply stabilized by individual contacts but is held in place by the combined polar and hydrophobic architecture created by these five residues.

### Boltz-2 predicts novel binding site for IL-23’s first-in-class inhibitor 39

Recently, an inhibitory peptide was crystallized with IL-23, revealing an induced-fit orthosteric pocket on the IL-23p19 subunit (2025).^200^ Building on this structural insight, we applied Boltz-2 to predict the binding interactions of our in-house IL-23 inhibitor, compound **39**. As an initial validation, we reproduced the peptide–IL-23 interaction (reported in 2025)^200^ using Boltz-2 and confirmed its overlap with the corresponding PDB structure **(Figure S8)**. Notably, Boltz-2 predicted that inhibitor **39** binds to the IL-12 subunit of the IL-23 heterodimer with a high confidence score of 0.882, suggesting a distinct binding mode that could provide novel avenues for IL-23 inhibition **(Figure 7C)**. To further validate this prediction, we examined the Boltz-2–predicted poses of a SMM-negative compound (non-positive) and a validated binder.

**Figure 7.**
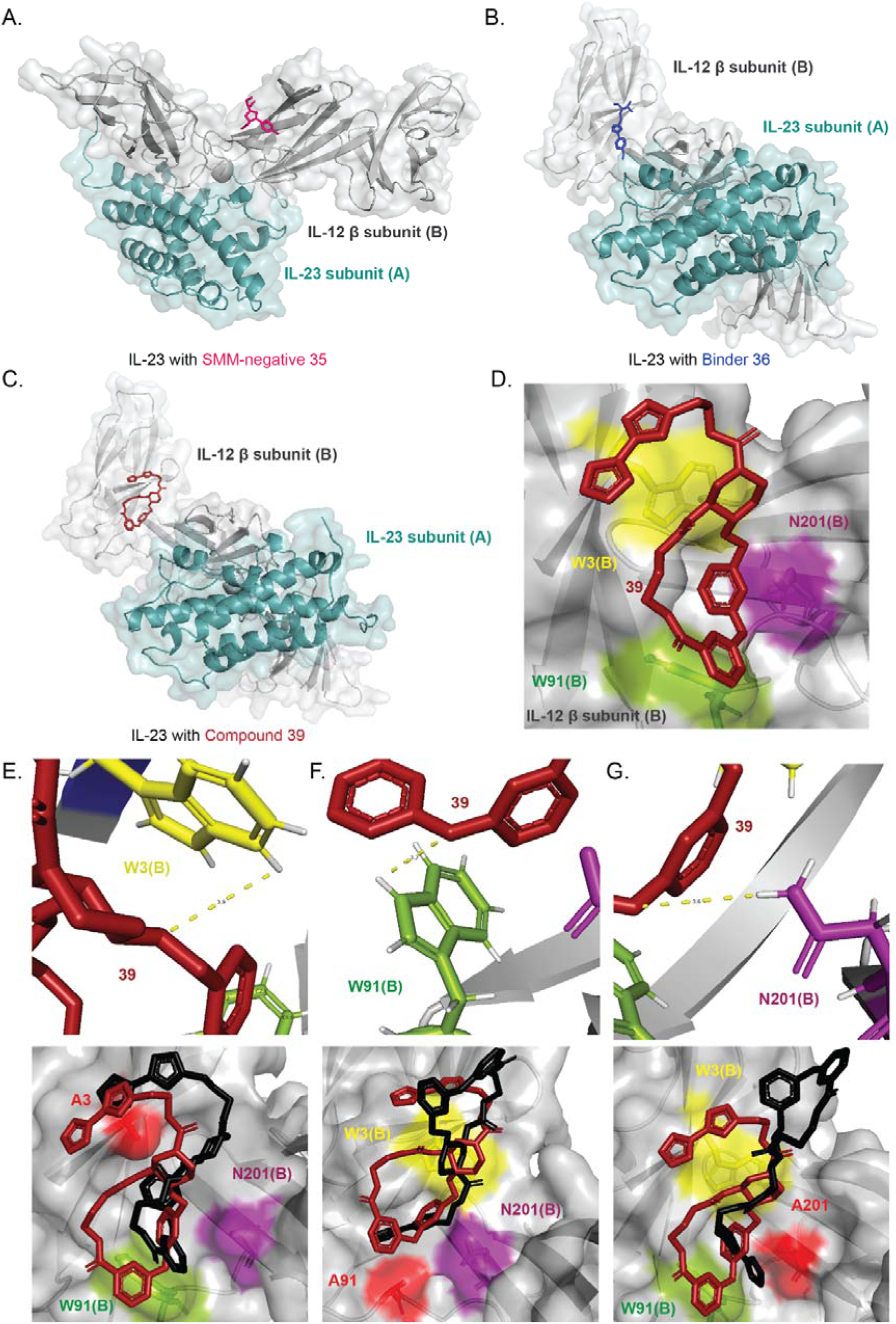
Boltz-2 prediction of IL-23 inhibitor 39 binding and *in silico* mutagenesis of the IL-12**β** interface pocket. **(A)** SMM-negative compound 35 (red) shows no detectable engagement with IL-23, remaining fully solvent-exposed on the IL-12β subunit.**(B)** Binder 36 (blue) engages the IL-12β subunit weakly, without forming a defined binding network. **(C)** Inhibitor **39** (dark red) binds at the IL-12β subunit interface, occupying a shallow cavity formed at the junction between IL-12β (gray) and IL-23p19 (teal). **(D)** Close-up view of the binding triad surrounding inhibitor **39**, highlighting W3(B) (yellow), W91(B) (green), and N201(B) (magenta) from the IL-12β subunit. These residues form a compact aromatic–polar groove that accommodates inhibitor **39**. **(E–G)** *In silico* alanine-scanning mutagenesis using Boltz-2 identifies the contribution of each residue to ligand stabilization. Substitution of W3(B), W91(B), or N201(B) with alanine (red) perturbs local molecular packing and disrupts the predicted pose of inhibitor **39**. The black conformation represents the displaced ligand orientation observed upon mutation, illustrating the destabilization that occurs when individual residues within the IL-12β interface pocket are perturbed.

As expected, the SMM-negative compound showed no detectable engagement with IL-23 **(Figure 7A)**. In contrast, compound **35**, a confirmed binder that produced ∼20% inhibition at 10 µM in the HEK-Blue IL-23 reporter assay, was predicted to bind at the IL-12 subunit of the IL-23 heterodimer **(Figure 7B)**.

We further identified key IL-12 subunit residues contributing to the inhibitor **39** binding, including Trp3(B), Trp91(B), and Asn201(B) **(Figure 7D–G**). Trp3(B) lies along the lower aromatic region of inhibitor **39**, where the benzene edge of its indole ring is oriented toward the diaryl ether oxygen that bridges two aromatic rings in the macrocycle, in a geometry consistent with a weak edge-on π–O interaction **(Figure 7E)**. Substitution of Trp3(B) with alanine disrupts this peripheral contact and causes the ligand to pivot toward the Trp91–Asn201 region, indicating that Trp3(B) helps position the inhibitor within the cleft **(Figure 7E)**.Trp91(B) provides the dominant anchoring interaction. Its indole N–H donates a hydrogen bond to the macrocyclic backbone ether oxygen adjacent to the secondary amide, while the indole ring simultaneously engages the nearby aromatic ring of the ligand through π-stacking **(Figure 7F)**. Mutation of Trp91(B) abolishes these contacts and yields a flipped, more solvent-exposed pose, establishing Trp91(B) as the primary determinant of ligand burial **(Figure 7F)**. Asn201(B) contributes a secondary stabilizing contact, with its side-chain amide forming a hydrogen bond to the same backbone ether oxygen engaged by Trp91(B) **(Figure 7G)**. Alanine substitution induces a similar flipped orientation, indicating that Asn201(B) works cooperatively with Trp91(B) to maintain ligand orientation within the pocket **(Figure 7G)**.

## Discussion and Conclusion

Profiling 32 human cytokines uncovered distinct SMM binding signatures for nearly every cytokine, demonstrating that soluble cytokines, traditionally viewed as challenging small-molecule targets, possess unique chemical interaction profiles that can be exploited for ligand discovery. Subsequent thermal shift validation of 864 enriched probes resulted in 296 confirmed cytokine binders, reflecting an average translation rate of 32.5% and highlighting that a substantial fraction of SMM-derived interactions persists across orthogonal biophysical formats. Evaluation of three prioritized cytokines (IL-17, IL-13, and IL-23) in HEK-Blue reporter assays further revealed variable translation rates and potencies, consistent with differences in structural topography and potential binding sites. IL-17 inhibitors displayed potencies from single to double-digit micromolar inhibitors, whereas first-in-class IL-13 and IL-23 inhibitors, respectively, showed a narrower potency range, potentially reflecting the comparatively flatter engagement surfaces observed for these cytokines.

The observed variation in potencies across the three cytokines likely reflects differences in their underlying structural landscapes, which is further supported by predictive insights from Boltz-2 modeling. For IL-17, Boltz-2 predicted that inhibitors **5**, **9**, and **24** occupy the C-terminal site with confidence scores of 0.839, 0.799, and 0.822, respectively. Among these, compounds **9** and **24** showed the strongest cellular potencies (6.43 µM and 3.16 µM), suggesting that engagement of this region may contribute to more effective inhibition. In contrast, compound **5**, despite being predicted to bind the same site, displayed more modest activity (29.9 µM). In contrast, the macrocyclic analog **22** was predicted to bind at the center of the dimer interface, engaging site 2 residues with a confidence score of 0.825, the same region targeted by Eli Lilly and Pfizer macrocyclic antagonists but localized in a different part of site 2. For IL-13, which presents a uniformly shallow engagement surface, Boltz-2 predicted that compound **28** binds helix D with a confidence score of 0.819, while compound **34** engages the A to C helical interface with a confidence score of 0.729. IL-23, although generally flatter than IL-17, contains localized shallow grooves on the IL-12C subunit. Boltz-2 reproduced the recently reported peptide-bound pose for IL-23 (2025) and predicted that compound **39** binds a distinct region on IL-12C with a high confidence score of 0.882. Collectively, these modeling results, while not experimentally validated structures, align with potency trends observed in biochemical assays and help explain why IL-17 supports higher potencies. In contrast, structural constraints associated with IL-13 and IL-23 likely underline the narrower potency range observed for their inhibitors, reflecting the long-standing challenge of identifying potent and selective small molecules for these cytokines.

These findings highlight several compelling future directions. First, the IL-13 inhibitors identified here provide a foundation for structure–activity relationship development, particularly around residues known to engage IL-13 monoclonal antibodies. Given the relatively shallow surface features of IL-13, fragment-based growth of these chemotypes or strategic extension into additional interaction patches may help yield higher affinity and more selective inhibitors. Second, the identification of a first-in-class small-molecule IL-23 inhibitor suggests an additional path for cytokine modulation. Future optimization of this heteroatom-rich macrocyclic chemotype could focus on tuning macrocycle rigidity, polar surface distribution, and peripheral aromatic contacts to enhance affinity and selectivity, while probing whether p40-directed small molecules represent a generalizable strategy for modulating IL-12/IL-23 signaling. Finally, integrating AI-driven modeling with the expansive SMM dataset generated for 32 cytokines offers a unique opportunity to develop predictive frameworks for cytokine ligandability, chemical preference, and selectivity. Such models could guide the prioritization of emerging chemotypes, identify transferable substructures across cytokine families, and accelerate the optimization of potency and pharmacological properties.

More broadly, this study represents the first systematic, combinatorial exploration of small-molecule tractability across a large panel of immunoregulatory cytokines. By combining high-throughput SMM screening, biophysical validation, cellular reporter assays, and predictive structural modeling, we expand the landscape of feasible cytokine targets and uncover first-in-class inhibitors for IL-13 and IL-23, as well as novel inhibitors for IL-17. These results challenge the prevailing assumption that soluble cytokines are uniformly undruggable and demonstrate that diverse cytokines possess chemically addressable regions suitable for ligand discovery. Collectively, this work establishes a framework for discovering small-molecule modulators of traditionally recalcitrant cytokines, underscores the value of integrating chemical diversity with orthogonal screening modalities, and sets the stage for the development of next-generation cytokine inhibitors with therapeutic relevance in immune-mediated diseases and cancer.

## Lead Contact

Further information and requests for resources and reagents should be directly addressed to the lead contact, Angela N. Koehler

## Materials availability

All compounds reported in this study are commercially available from Chembridge corporation. Please refer to their respective identifier under materials & methods section.

## Data and code availability

Any additional information required to reanalyze the data reported in this paper and code utilized for cheminformatics platform is available from the lead contact upon request.

## Acknowledgment

This publication was supported by the Boston University Ignition Award, a Koch Institute Frontier Award, and funding from the MIT Center for Precision Cancer Medicine (CPCM).

## Author Contributions

Conceptualization: A.J.V., A.N.K. and R.; SMM manufacturing: B.L., B.L., M.S., SMM screening: R., S.P.Q.; Cheminformatics platform: A.C., R., S.P.Q., Secondary binding validation: R., Hierarchical clustering: R., A.J.V., Cell-based assays: R., Boltz-2 studies: R., A.J.V., Data interpretation: R., A.J.V., and A.N.K. Original Draft writing and conceptualization: R., A.J.V., writing-review & editing, all authors: funding acquisition & supervision: A.J.V. & A.N.K.

## Materials & Methods

### Key Resources Table

#### Cytokines used for SMM screening

**Cytokines used for SMM screening:**
| No. | Cytokines | Tag | Identifier | Source | Expression |
| --- | --- | --- | --- | --- | --- |
| 1 | Human IL-1 alpha/IL-1F1 protein | His | ILA-H5216 | AcroBiosystems | HEK293 |
| 2 | Human IL-1 beta/IL-F2 protein | His | ILB-H51H3 | AcroBiosystems | HEK293 |
| 3 | Human IL-2 protein | His | IL2-H52H8 | AcroBiosystems | HEK293 |
| 4 | Human IL-5 protein | His | IL5-H52H3 | AcroBiosystems | HEK293 |
| 5 | Human IL-6 protein | His | IL6-H5243 | AcroBiosystems | HEK293 |
| 6 | Human IL-7 protein | His | Z03190-25 | Genscript | CHO |
| 7 | Human IL-8.CXCL-8 protein | His | IL8-H52H3 | AcroBiosystems | HEK293 |
| 8 | Human IL-10 protein | His | IL0-H4248 | AcroBiosystems | HEK293 |
| 9 | Human IL-11 Protein | His | IL1-H5243 | AcroBiosystems | HEK293 |
| 10 | Human IL-12 protein | His | PKSH030475 | Elabscience | HEK293 |
| 11 | Human IL-13 protein | His | IL3-H52H4 | AcroBiosystems | HEK293 |
| 12 | Human IL-17A/CTLA-8 protein | His | ILA-H5243 | AcroBiosystems | HEK293 |
| 13 | Human IL-8 protein | His | IL8-H52H3 | AcroBiosystems | HEK293 |
| 14 | Human IL-21 protein | His | ab235875 | Abcam | Baculovirus infected insect cells |
| 15 | Human IL-22 protein | His | IL2-H524a | Acrobiosystems | HEK293 |
| 16 | Human IL-23 alpha & IL-12 beta Heterodimer Protein, His Tag&Tag Free | His | ILB-H52W5 | Acrobiosystems | HEK293 |
| 17 | Human IL-33 protein | His | IL3-H52H7 | Acrobiosystems | HEK293 |
| 18 | Human CD40 Ligand / TNFSF5 Protein, His,Flag Tag (active trimer) | His | CDL-H52Db | Acrobiosystems | HEK293 |
| 19 | Human Complement C5 protein | His | CO5-H52Ha | Acrobiosystems | HEK293 |
| 20 | Biotinylated Human GM-CSF, epitope tag free, ultra sensitivity (primary amine labeling) | Biotinylated | GMF-H8214 | Acrobiosystems | HEK293 |
| 21 | Human G-CSF Protein | His | RP75570 | Thermo Fisher Scientific | E.coli |
| 22 | Biotinylated Human MIF Protein, His,Avitag™ | His | MIF-H82E7 | Acrobiosystems | HEK293 |
| 23 | Human IFN-alpha 1 Protein | His | IFA-H52H9 | Acrobiosystems | HEK293 |
| 24 | Biotinylated Human TGF-Beta 1 / TGFB1 Protein, Avitag™ | Biotinylated | TG1-H8217 | Acrobiosystems | HEK293 |
| 25 | Human TNF-alpha Protein (active trimer) | His | TNA-H5228 | Acrobiosystems | HEK293 |
| 26 | Biotinylated Human CCL1 / I-309 / C-C motif chemokine 1 Protein, His,Avitag™ | His | CC1-H82E9 | Acrobiosystems | HEK293 |
| 27 | MCP-1/CCL2 Protein, Human (Biotinylated, HEK293, His-Avi) | His | HY-P78173 | MedChemExpress | HEK293 |
| 28 | Human CCL3 protein | His | CCL3-475H | Creative Biomart | HEK293 |
| 29 | Human CCL5 protein | His | CCL3-475H | Creative Biomart | HEK293 |
| 30 | Human CXCL1 protein | His | PKSH032290 | Elabscience | HEK293 |
| 31 | Human CXCL10 Protein (Fc & His Tag) | His | PKSH032292 | Elabscience | HEK293 |
| 32 | Human CXCL12 protein | His | CXCL12-1219H | Creative Biomart | HEK293 |

#### Cytokines used for DSF, nanoDSF and cell-based studies

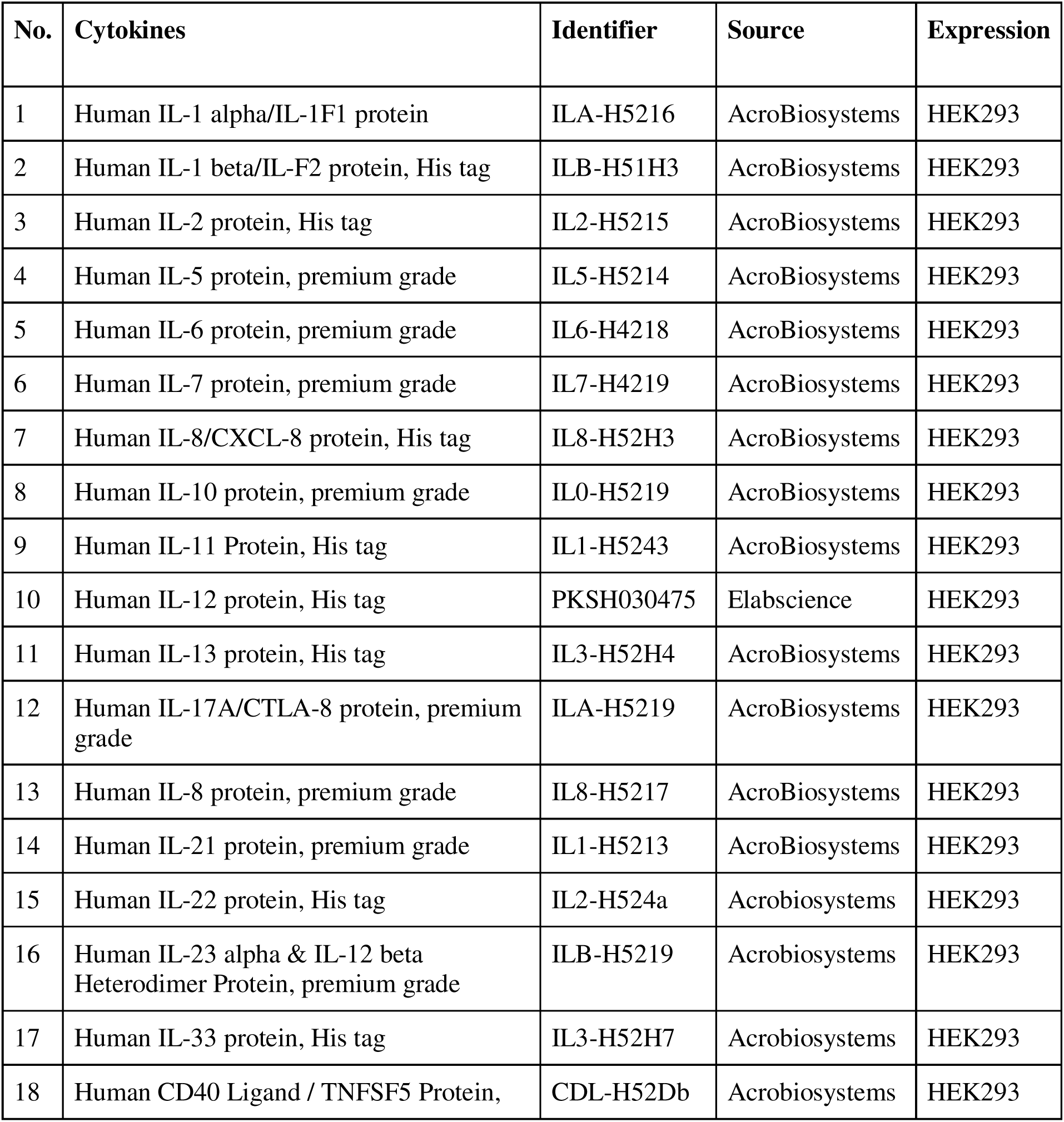
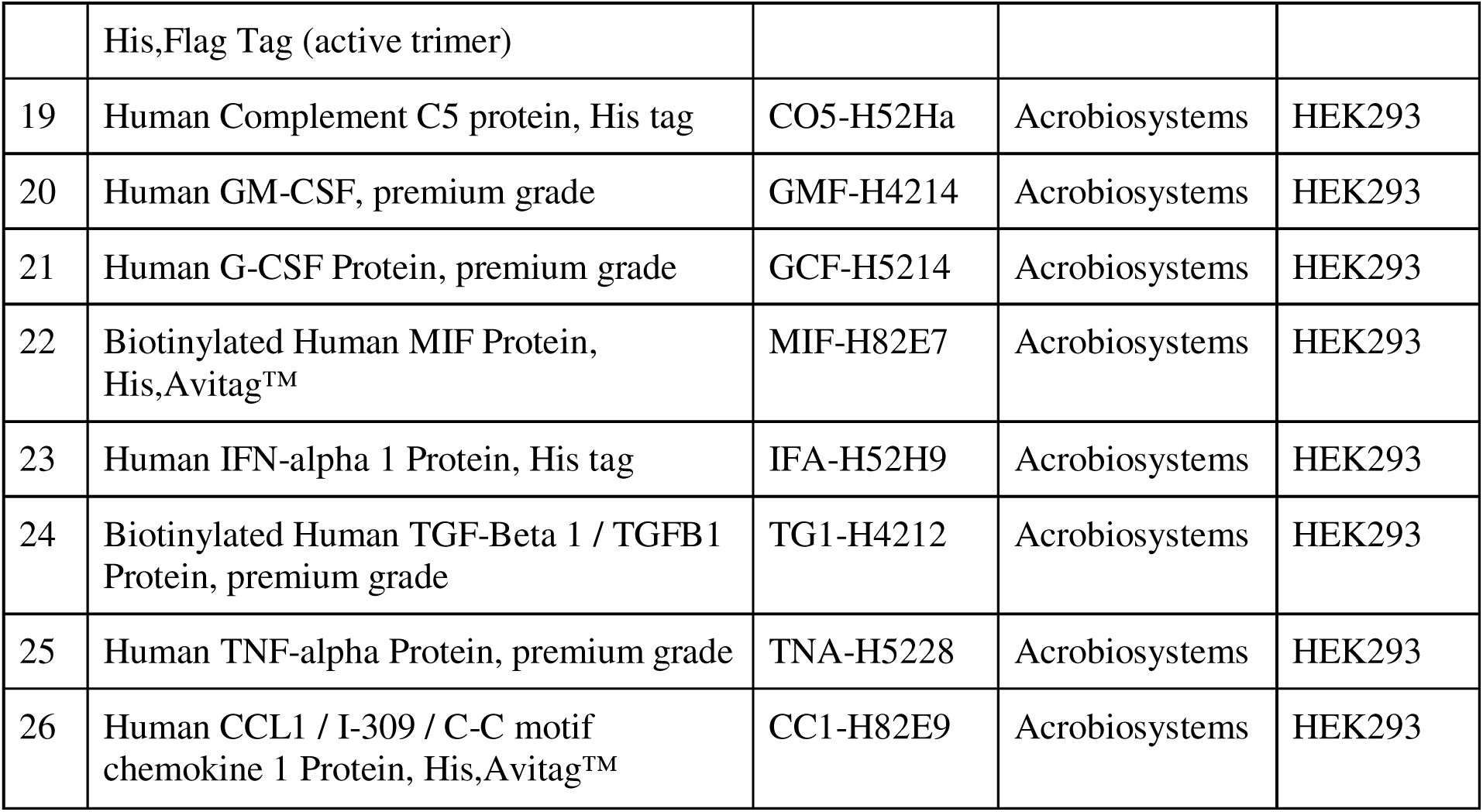

**Cytokines used for DSF, nanoDSF and cell-based studies:**
| No. | Cytokines | Identifier | Source | Expression |
| --- | --- | --- | --- | --- |
| 1 | Human IL-1 alpha/IL-1F1 protein | ILA-H5216 | AcroBiosystems | HEK293 |
| 2 | Human IL-1 beta/IL-F2 protein, His tag | ILB-H51H3 | AcroBiosystems | HEK293 |
| 3 | Human IL-2 protein, His tag | IL2-H5215 | AcroBiosystems | HEK293 |
| 4 | Human IL-5 protein, premium grade | IL5-H5214 | AcroBiosystems | HEK293 |
| 5 | Human IL-6 protein, premium grade | IL6-H4218 | AcroBiosystems | HEK293 |
| 6 | Human IL-7 protein, premium grade | IL7-H4219 | AcroBiosystems | HEK293 |
| 7 | Human IL-8/CXCL-8 protein, His tag | IL8-H52H3 | AcroBiosystems | HEK293 |
| 8 | Human IL-10 protein, premium grade | IL0-H5219 | AcroBiosystems | HEK293 |
| 9 | Human IL-11 Protein, His tag | IL1-H5243 | AcroBiosystems | HEK293 |
| 10 | Human IL-12 protein, His tag | PKSH030475 | Elabscience | HEK293 |
| 11 | Human IL-13 protein, His tag | IL3-H52H4 | AcroBiosystems | HEK293 |
| 12 | Human IL-17A/CTLA-8 protein, premium grade | ILA-H5219 | AcroBiosystems | HEK293 |
| 13 | Human IL-8 protein, premium grade | IL8-H5217 | AcroBiosystems | HEK293 |
| 14 | Human IL-21 protein, premium grade | IL1-H5213 | AcroBiosystems | HEK293 |
| 15 | Human IL-22 protein, His tag | IL2-H524a | Acrobiosystems | HEK293 |
| 16 | Human IL-23 alpha & IL-12 beta Heterodimer Protein, premium grade | ILB-H5219 | Acrobiosystems | HEK293 |
| 17 | Human IL-33 protein, His tag | IL3-H52H7 | Acrobiosystems | HEK293 |
| 18 | Human CD40 Ligand / TNFSF5 Protein, | CDL-H52Db | Acrobiosystems | HEK293 |
|  | His,Flag Tag (active trimer) |  |  |  |
| 19 | Human Complement C5 protein, His tag | CO5-H52Ha | Acrobiosystems | HEK293 |
| 20 | Human GM-CSF, premium grade | GMF-H4214 | Acrobiosystems | HEK293 |
| 21 | Human G-CSF Protein, premium grade | GCF-H5214 | Acrobiosystems | HEK293 |
| 22 | Biotinylated Human MIF Protein,<br>His,Avitag™ | MIF-H82E7 | Acrobiosystems | HEK293 |
| 23 | Human IFN-alpha 1 Protein, His tag | IFA-H52H9 | Acrobiosystems | HEK293 |
| 24 | Biotinylated Human TGF-Beta 1 / TGFB1<br>Protein, premium grade | TG1-H4212 | Acrobiosystems | HEK293 |
| 25 | Human TNF-alpha Protein, premium grade | TNA-H5228 | Acrobiosystems | HEK293 |
| 26 | Human CCL1 / I-309 / C-C motif<br>chemokine 1 Protein, His,Avitag™ | CC1-H82E9 | Acrobiosystems | HEK293 |

| **Reagent/Resource** | **Identifier** | **Source** |
| --- | --- | --- |
| Alexa Fluor® 647 mouse anti-His Tag Antibody | 652513 | Biolegend |
| Streptavidin, Alexa Fluor™ 647 Conjugate | S21374 | Thermo Fisher Scientific |
| Bovine Serum Albumin | A7030 | Sigma Aldrich |
| Gibco PBS, pH 7.4 | 10010072 | Thermo Fisher Scientific |
| DMSO (Dimethyl Sulfoxide , Anhydrous) Alfa<br>Aesar >99.8% | AA43998AE | Fisher Scientific |
| DMSO (Dimethyl Sulfoxide, Fisher<br>BioReagents™) | BP231-100 | Fisher Scientific |
| Tween-20, Molecular Biology Grade, Promega™ | PRH5152 | Fisher Scientific |
| Applied Biosystems Protein Thermal Shift Dye<br>Kit | 44-611-46 | Fisher Scientific |
| Hard-Shell PCR Plates, 384-well, thin-wall | HSP3805 | Bio-rad |
| Adhesive PCR Plate Seals | AB-0558 | Thermo Fisher Scientific |
| **Reporter cell lines** | **Identifier** | **Source** |
| HEK Blue IL-17A reporter cell line | Invivogen | hkb-il17 |
| HEK Blue IL-4/IL-13 reporter cell line | Invivogen | hkb-il413 |
| HEK Blue IL-23 reporter cell line | Invivogen | hkb-il23 |
| **Reagents/Resources for Cell-based studies** | **Identifier** | **Source** |
| Gibco™ DMEM, high glucose | 11-965-118 | Fisher Scientific |
| Gibco™ Fetal Bovine Serum, qualified, heat<br>inactivated | 10-438-026 | Fisher Scientific |
| 96-Well Clear Bottom Plates, Corning®, Plates<br>without Lids, Nonbinding | 07-200-770 | Fisher Scientific |
| MicroAmp Fast Optical 96 Well Reaction Plate | 43-469-0 | Thermo Fisher Scientific |
| Corning Costar Flat Bottom Cell Culture Plates, 96 well | 07-200-89 | Fisher Scientific |
| Nalgene; Sterile Disposable Filter Units with PES Membrane | 09-741-04 | Thermo Fisher Scientific |
| Blasticidin | ant-bl-05 | Invivogen |
| Zeocin | ant-zn-05 | Invivogen |
| Normocin | ant-nr-1 | Invivogen |
| Selection | hb-sel | Invivogen |
| Quanti-Blue | rep-qbs | Invivogen |
| **Compounds (structure shown in the manuscript and SI)** | **Identifier** | **Source** |
| **1** | 90009505 | ChemBridge Corporation |
| **2** | 9294722 | ChemBridge Corporation |
| **3** | 23186653 | ChemBridge Corporation |
| **4** | 9141738 | ChemBridge Corporation |
| **5** | 79438440 | ChemBridge Corporation |
| **7** | 7968494 | ChemBridge Corporation |
| **8** | 69283331 | ChemBridge Corporation |
| **9** | 9099148 | ChemBridge Corporation |
| **10** | 21876765 | ChemBridge Corporation |
| **11** | 99883887 | ChemBridge Corporation |
| **12** | 35030361 | ChemBridge Corporation |
| **13** | 31585875 | ChemBridge Corporation |
| **14** | 52590279 | ChemBridge Corporation |
| **15** | 70031656 | ChemBridge Corporation |
| **16** | 15024932 | ChemBridge Corporation |
| **17** | 14261167 | ChemBridge Corporation |
| **18** | 76447749 | ChemBridge Corporation |
| **19** | 15025039 | ChemBridge Corporation |
| **20** | 40079105 | ChemBridge Corporation |
| **21** | 10923751 | ChemBridge Corporation |
| **22** | 37049039 | ChemBridge Corporation |
| **23** | 19477454 | ChemBridge Corporation |
| **24** | 10577678 | ChemBridge Corporation |
| **25** | 22238812 | ChemBridge Corporation |
| **26** | 44388455 | ChemBridge Corporation |
| **27** | 62638488 | ChemBridge Corporation |
| **28** | 39820255 | ChemBridge Corporation |
| **29** | 56112019 | ChemBridge Corporation |
| **30** | 52055947 | ChemBridge Corporation |
| **31** | 73700720 | ChemBridge Corporation |
| **32** | 79923876 | ChemBridge Corporation |
| **33** | 71731196 | ChemBridge Corporation |
| **34** | 83438613 | ChemBridge Corporation |
| **35** | 12095577 | ChemBridge Corporation |
| **36** | 94201500 | ChemBridge Corporation |
| **37** | 69679209 | ChemBridge Corporation |
| **38** | 65814991 | ChemBridge Corporation |
| **39** | 14212139 | ChemBridge Corporation |

\*All commercial small molecules used in SMM screening were purchased from ChemBridge Corporation. Validated binders from thermal shift assays were purchased from ChemBridge Corporation. Compounds used for cell-based potency determination were dissolved in anhydrous DMSO and were aliquoted (10 mM stock concentration) and stored in – 80°C with desiccant pouches upon arrival.
\*\*Compound **9** loses activity within 4 weeks after solubilization, authors recommend using a relatively fresh batch of this compound for any studies.

#### Small molecule microarray screening of proinflammatory cytokines

Each SMM slide contained approximately 4000 printed features in duplicates. A total of 65,000 compounds were screened. The collection contained ‘lead-like’ compounds and commercially available compounds. Each slide was preincubated with 3 mL of 23 µM BSA in 1X PBS-T for 30 minutes at room temperature. The slides were then incubated for 30 minutes with a 3 mL solution of 2.3 nM Cytokine in 23 µM BSA, 1X PBS-T. Slides were then washed thrice for 2 minutes each with 1X PBS-T. Prior to antibody incubation, chambers were replaced to prevent any remaining protein that might lead to a background signal. Washed slides were then incubated with a 3 mL solution of anti-His mouse monoclonal antibody conjugated to AlexaFluor647 at a 1:2000 concentration for 30 minutes. For biotinylated cytokines, Streptavidin-AlexaFluor647 conjugate was used at 1:10,000 concentration. Slides were again washed thrice with 1X PBS-T and twice with 1X PBS. Finally, the slides were briefly rinsed twice with distilled water and spin-dried. The slides were scanned immediately using the GenePix 4000B fluorescence scanner (Molecular Devices). The images were analyzed using GenePix Pro software (Axon Instruments) and data was analyzed using Pipeline software as described below.

### SMM Statistical Analysis (SMM positive calling)

Raw data was analyzed based on the signal-to-noise ratio and reproducibility. For each feature, the signal-to-noise ratio (SNR) was defined as the median fluorescence intensity of the feature divided by the median fluorescence intensity of the surrounding slide area, defined as a radius 3 times the radius of the spot, excluding pixels within a certain overlap threshold of neighboring features. Then, for each feature, a robust z-score (z_i_), which is less influenced by outliers compared with the mean-based z statistic, was calculated for each feature (i) by the following equation:

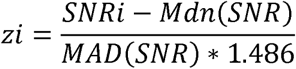

Where SNR_i_ is the SNR value for a given feature, *Mdn* (SNR) is the median of the SNR values for all features in the subarray, and the *MAD* (SNR) is the maximum absolute deviation of the SNR values for all features in the subarray. A composite z_i_ score of all the features was calculated for every slide and its duplicate to prevent batch-wise slide differences. Assay positives were defined as features with z_i_ scores ≥ 3 on each slide and reproducible across duplicate slides. Assay positives were then compared with antibody counter screens to filter out non-specific binders. Unique assay positives for each cytokine were pursued further.

### Fragment enrichment analysis

Screening molecules were fragmented using BRICS and RECAP rules (RDKit) and by enumerating all scaffolds contained within each molecule (ScaffoldGraph^201^). For each target, the number of molecules containing each fragment was counted in both the SMM positive set and the remaining non-positive portion of the screening library (not-positive set). Fragment frequency in both the SMM positive set and the not-positive set was calculated as the ratio of the number of molecules containing the fragment and the total number of molecules in the set. Fragment enrichment in the SMM positive set was defined as the ratio of fragment frequency in the SMM positive set over fragment frequency in the not-positive set. To evaluate the statistical significance of the calculated enrichments, Fisher’s exact test was performed, and p-values were adjusted to account for multiple test comparisons with the Benjamini-Hochberg procedure. Fragments with enrichment > 1 and adjusted p-value ≤ 0.05 were considered significantly enriched and the associated molecules were prioritized for further evaluation.

### Secondary binding validation of SMM positives using nanoDSF

10 mM stocks were prepared for each assay positive in 100% DMSO. Each assay positive was screened at a final concentration of 2 µM in 0.1% DMSO in duplicates against 1 µg of cytokine (1X PBS, pH = 7.4). Melting curves for each sample were obtained with the following conditions: 20-95°C at a ramp rate of 1°C/minute. Excitation power was adjusted to achieve fluorescence readings between 2,000-20,000 RFU for emission at 330 and 350 nm.

### Secondary binding validation of SMM positives using DSF

Unique positives for each cytokine was screened at a final concentration of 2 µM in 0.1% DMSO in duplicates. 1 µg of cytokine in PBS was added to 384 well plates, followed by 8X Spiro-orange, PBS, DMSO or SMM positives in duplicates. DSF was performed on a BioRad CFX384. The alteration in melting temperature was assessed by contrasting the lowest point of the protein’s melting curve in the presence of a vehicle with that in the presence of a small molecule.

### HEK Blue IL-4/IL-13 reporter cell line assay

HEK Blue IL-4/IL-13 reporter cell lines were maintained in DMEM high glucose with 10% Heat Inactivated Fetal Bovine Serum (Fisher Scientific), 10 µg/mL Blasticidin, 200 µg/mL Zeocin, 100 µg/mL Normocin, and 100 U/mL-100 µg/mL Pen-Strep. Inhibitory activity assays were conducted with cells between passages 11-17. Firstly, IL-13 induced SEAP production was tested in dose response manner with 0.2% DMSO. The EC_25_ for IL-13 induced SEAP production (8 ng/mL final well concentration) was chosen for screening small molecules.

For screening small molecules in a dose-dependent manner, 25 µL of analog in DMEM with 4% DMSO (final well DMSO concentration becomes 0.4%) was incubated with 25 µL of 80 ng/mL IL-13 in DMEM for 15-30 minutes at room temperature in a 96 well reaction plate (MicroAmp^TM^, Fisher Scientific). In the meantime, HEK Blue IL-4/IL-13 cells were washed, trypsinized, spun down, and resuspended in DMEM with 10% Heat Inactivated FBS and 100 U/mL-100 µg/mL Pen-Strep at a concentration of 3.125E5 cells/mL. 160 µL of resuspended cells was added to each well of a 96-well plate (Corning Costar, Flat bottom), avoiding the edges of the plate, and equilibrated in a 37°C, 5% CO_2_ incubator for 30 minutes. After 60-90 minutes, 40 µL of the small molecule/IL-13 solution mix was added to each well with cells. Each measurement was performed in quadruplicate. Cells were incubated for 22-24 hours at 37°C, 5% CO_2_. Quanti-Blue dye was used for IL-13 induced SEAP production. A stock solution of Quanti-Blue dye was prepared as per the manufacturer’s recommendation. 160 µL of the prepared dye was added to each well of a 96-well plate, 40 µL of cell supernatant was added to each well avoiding edges, and then incubated for 1-3 hours at 37°C. The optical density was then read at 650 nm with a Molecular Devices SpectraMax M5 Microplate Reader. Data was plotted and analyzed in GraphPad Prism 9.

### HEK Blue IL-23 reporter cell line assay

HEK-Blue IL-23 reporter cell lines were cultured in DMEM with 10% Heat Inactivated Fetal Bovine Serum (FBS), 100 U/mL-100 µg/mL Pen-Strep, 100 µg/mL Normocin and 1X Selectin. Cells were tested for SEAP production induced by Tag-free HEK293 expressed IL-23 alone prior to small molecule screening. EC_50_ for IL-23 was determined to be 1.8 ng/mL. The EC_25_ for IL-23 induced SEAP production (0.9 ng/mL final well concentration) was chosen for screening small molecules to determine their inhibitory activity.

22.5 µL of small molecule in DMEM 4% DMSO was added to 22.5 µL of 9 ng/mL IL-23 in DMEM in a non-binding 96-well reaction plate for 15-30 minutes at room temperature. HEK-Blue IL-23 reporter cell line was used between passages 11-17. Cells were washed with 1X PBS, trypsinized using 0.25% Trypsin-EDTA (1X) for 2 minutes and centrifuged for 5 minutes at 250xg. Cells were resuspended in DMEM 10% FBS and 100 U/mL-100 µg/mL Pen-Strep at a concentration of 3.125 x 10^5^ cells /mL.160 µL of cell suspension was added to each well of a 96-well plate, avoiding edges, and incubated for 30 minutes in a 37°C, 5% CO_2_ incubator. Lastly, 40 µL of IL-17 with small molecule/ vehicle solution was added to the cells preincubated in a 96-well plate. Measurements were done in quadruplicate. The plate was incubated for ∼22-24 hours at 37°C, 5% CO_2_. Quanti-Blue dye was prepared as per the manufacturer’s recommendation and 160 µl was dispensed in each well of a 96-well plate (avoid edges to avoid evaporation), 40 µl of cell supernatant was added to the same plate and incubated for 3 hours at 37°C, 5% CO_2_. Optical density was then measured at 650 nm using Molecular Devices SpectraMax M5 Microplate reader. Data was analyzed and plotted using GraphPad Prism 9.

### HEK Blue IL-17 reporter cell line assay

HEK-Blue IL-17 reporter cell lines were cultured in DMEM with 10% Heat Inactivated Fetal Bovine Serum (FBS), 100 U/mL-100 µg/mL Pen-Strep, 100 µg/mL Normocin and 1X Selectin. Cells were tested for SEAP production induced by Tag-free HEK293 expressed IL-17 alone prior to small molecule screening. EC_50_ for IL-17 was determined to be 2.66 ng/mL. The EC_25_ for IL-17 induced SEAP production (1.33 ng/mL final well concentration) was chosen for screening small molecules to determine their inhibitory activity.

22.5 µL of small molecule in DMEM 4% DMSO was added to 22.5 µL of 13.3 ng/ml IL-17 in DMEM in a non-binding 96-well reaction plate for 15-30 minutes at room temperature. HEK-Blue IL-17 reporter cell line was used between passages 11-17. Cells were washed with 1X PBS, trypsinized using 0.25% Trypsin-EDTA (1X) for 2 minutes and centrifuged for 5 minutes at 250xg. Cells were resuspended in DMEM 10% FBS and 100 U/mL-100 µg/mL Pen-Strep at a concentration of 3.125 x 10^5^ cells /mL.160 µL of cell suspension was added to each well of a 96-well plate, avoiding edges, and incubated for 30 minutes in a 37°C, 5% CO_2_ incubator. Lastly, 40 µL of IL-17 with small molecule/ vehicle solution was added to the cells preincubated in a 96-well plate. Measurements were done in quadruplicate. The plate was incubated for ∼22-24 hours at 37°C, 5% CO_2_. Quanti-Blue dye was prepared as per the manufacturer’s recommendation and 160 µL was dispensed in each well of a 96-well plate (avoid edges to avoid evaporation), 40 µL of cell supernatant was added to the same plate and incubated for 3 hours at 37°C, 5% CO_2_. Optical density was then measured at 650 nm using Molecular Devices SpectraMax M5 Microplate reader. Data was analyzed and plotted using GraphPad Prism 9.

### Boltz-2 studies

FASTA sequences for IL-17 (PDB ID: 4HR9), IL-13 (PDB ID: 3BL6), and IL-23 (PDB ID: 8UUI) were obtained from the RCSB Protein Data Bank and used as inputs for initial folding predictions using the Boltz-2 algorithm implemented through the Tamarind Bio platform^202^. Predicted structures were evaluated in PyMOL to confirm correct global folding and alignment with the corresponding experimental PDB references. After validation, SMILES strings for each compound were supplied together with the cytokine FASTA sequence to generate protein–ligand binding predictions.

For *in silico* site-directed mutagenesis, individual amino acid substitutions were introduced directly into the cytokine FASTA sequence, and each mutant was run independently with the corresponding compound using Boltz-2 via the Tamarind Bio tool. This allowed residue-specific assessment of predicted binding pose changes upon single-site perturbation.

## References

(1) O’Shea, J. J.; Murray, P. J. Cytokine Signaling Modules in Inflammatory Responses. Immunity 2008, 28 (4), 477–487. 10.1016/j.immuni.2008.03.002.

(2) Kulbe, H.; Chakravarty, P.; Leinster, D. A.; Charles, K. A.; Kwong, J.; Thompson, R. G.; Coward, J. I.; Schioppa, T.; Robinson, S. C.; Gallagher, W. M.; Galletta, L.; Australian Ovarian Cancer Study Group; Salako, M. A.; Smyth, J. F.; Hagemann, T.; Brennan, D. J.; Bowtell, D. D.; Balkwill, F. R. A Dynamic Inflammatory Cytokine Network in the Human Ovarian Cancer Microenvironment. Cancer Res 2012, 72 (1), 66–75. 10.1158/0008-5472.CAN-11-2178.

(3) Liu, C.; Chu, D.; Kalantar-Zadeh, K.; George, J.; Young, H. A.; Liu, G. Cytokines: From Clinical Significance to Quantification. Advanced Science 2021, 8 (15), 2004433. 10.1002/advs.202004433.

(4) Kany, S.; Vollrath, J. T.; Relja, B. Cytokines in Inflammatory Disease. Int J Mol Sci 2019, 20 (23), 6008. 10.3390/ijms20236008.

(5) Clark, R.; Kupper, T. Old Meets New: The Interaction Between Innate and Adaptive Immunity. Journal of Investigative Dermatology 2005, 125 (4), 629–637. 10.1111/j.0022-202X.2005.23856.x.

(6) Banyer, J. L.; Hamilton, N. H.; Ramshaw, I. A.; Ramsay, A. J. Cytokines in Innate and Adaptive Immunity. Rev Immunogenet 2000, 2 (3), 359–373.

(7) Dinarello, C. A. Historical Review of Cytokines. Eur J Immunol 2007, 37 (Suppl 1), S34–S45. 10.1002/eji.200737772.

(8) Yi, M.; Li, T.; Niu, M.; Zhang, H.; Wu, Y.; Wu, K.; Dai, Z. Targeting Cytokine and Chemokine Signaling Pathways for Cancer Therapy. Sig Transduct Target Ther 2024, 9 (1), 1–48. 10.1038/s41392-024-01868-3.

(9) Zhang, J.-M.; An, J. Cytokines, Inflammation and Pain. Int Anesthesiol Clin 2007, 45 (2), 27–37. 10.1097/AIA.0b013e318034194e.

(10) Van der Meide, P. H.; Schellekens, H. Cytokines and the Immune Response. Biotherapy 1996, 8 (3–4), 243–249. 10.1007/BF01877210.

(11) Steinke, J. W.; Borish, L. 3. Cytokines and Chemokines. Journal of Allergy and Clinical Immunology 2006, 117 (2), S441–S445. 10.1016/j.jaci.2005.07.001.

(12) Belardelli, F.; Ferrantini, M. Cytokines as a Link between Innate and Adaptive Antitumor Immunity. Trends in Immunology 2002, 23 (4), 201–208. 10.1016/S1471-4906(02)02195-6.

(13) Briukhovetska, D.; Dörr, J.; Endres, S.; Libby, P.; Dinarello, C. A.; Kobold, S. Interleukins in Cancer: From Biology to Therapy. Nat Rev Cancer 2021, 21 (8), 481–499. 10.1038/s41568-021-00363-z.

(14) Peters, M. C.; Kerr, S.; Dunican, E. M.; Woodruff, P. G.; Fajt, M. L.; Levy, B. D.; Israel, E.; Phillips, B. R.; Mauger, D. T.; Comhair, S. A.; Erzurum, S. C.; Johansson, M. W.; Jarjour, N. N.; Coverstone, A. M.; Castro, M.; Hastie, A. T.; Bleecker, E. R.; Wenzel, S. E.; Fahy, J. V. Refractory Airway Type 2 Inflammation in a Large Subgroup of Asthmatic Patients Treated with Inhaled Corticosteroids. Journal of Allergy and Clinical Immunology 2019, 143 (1), 104–113.e14. 10.1016/j.jaci.2017.12.1009.

(15) Hammad, H.; Lambrecht, B. N. The Basic Immunology of Asthma. Cell 2021, 184 (6), 1469–1485. 10.1016/j.cell.2021.02.016.

(16) Girolomoni, G.; Strohal, R.; Puig, L.; Bachelez, H.; Barker, J.; Boehncke, W. H.; Prinz, J. C. The Role of ILD23 and the ILD23/TH17 Immune Axis in the Pathogenesis and Treatment of Psoriasis. J Eur Acad Dermatol Venereol 2017, 31 (10), 1616–1626. 10.1111/jdv.14433.

(17) Krueger, J. G.; Eyerich, K.; Kuchroo, V. K.; Ritchlin, C. T.; Abreu, M. T.; Elloso, M. M.; Fourie, A.; Fakharzadeh, S.; Sherlock, J. P.; Yang, Y.-W.; Cua, D. J.; McInnes, I. B. IL-23 Past, Present, and Future: A Roadmap to Advancing IL-23 Science and Therapy. Front. Immunol. 2024, 15. 10.3389/fimmu.2024.1331217.

(18) McGeachy, M. J.; Chen, Y.; Tato, C. M.; Laurence, A.; Joyce-Shaikh, B.; Blumenschein, W. M.; McClanahan, T. K.; O’Shea, J. J.; Cua, D. J. The Interleukin 23 Receptor Is Essential for the Terminal Differentiation of Interleukin 17-Producing Effector T Helper Cells in Vivo. Nat Immunol 2009, 10 (3), 314–324. 10.1038/ni.1698.

(19) Aggarwal, S.; Ghilardi, N.; Xie, M.-H.; de Sauvage, F. J.; Gurney, A. L. Interleukin-23 Promotes a Distinct CD4 T Cell Activation State Characterized by the Production of Interleukin-17. J Biol Chem 2003, 278 (3), 1910–1914. 10.1074/jbc.M207577200.

(20) Di Cesare, A.; Di Meglio, P.; Nestle, F. O. The IL-23/Th17 Axis in the Immunopathogenesis of Psoriasis. J Invest Dermatol 2009, 129 (6), 1339–1350. 10.1038/jid.2009.59.

(21) Akdis, M.; Burgler, S.; Crameri, R.; Eiwegger, T.; Fujita, H.; Gomez, E.; Klunker, S.; Meyer, N.; O’Mahony, L.; Palomares, O.; Rhyner, C.; Quaked, N.; Schaffartzik, A.; Veen, W. V. D.; Zeller, S.; Zimmermann, M.; Akdis, C. A. Interleukins, from 1 to 37, and Interferon-γ: Receptors, Functions, and Roles in Diseases. Journal of Allergy and Clinical Immunology 2011, 127 (3), 701–721.e70. 10.1016/j.jaci.2010.11.050.

(22) Sundberg, T. B.; Xavier, R. J.; Schreiber, S. L.; Shamji, A. F. Small-Molecule Control of Cytokine Function: New Opportunities for Treating Immune Disorders. Current Opinion in Chemical Biology 2014, 23, 23–30. 10.1016/j.cbpa.2014.08.013.

(23) McCormick, S. M.; Heller, N. M. Commentary: IL-4 and IL-13 Receptors and Signaling. Cytokine 2015, 75 (1), 38–50. 10.1016/j.cyto.2015.05.023.

(24) Leonard, W. J.; Lin, J.-X.; O’Shea, J. J. The Γc Family of Cytokines: Basic Biology to Therapeutic Ramifications. Immunity 2019, 50 (4), 832–850. 10.1016/j.immuni.2019.03.028.

(25) Aggeletopoulou, I.; Assimakopoulos, S. F.; Konstantakis, C.; Triantos, C. Interleukin 12/Interleukin 23 Pathway: Biological Basis and Therapeutic Effect in Patients with Crohn’s Disease. World J Gastroenterol 2018, 24 (36), 4093–4103. 10.3748/wjg.v24.i36.4093.

(26) Chyuan, I.-T.; Lai, J.-H. New Insights into the IL-12 and IL-23: From a Molecular Basis to Clinical Application in Immune-Mediated Inflammation and Cancers. Biochemical Pharmacology 2020, 175, 113928. 10.1016/j.bcp.2020.113928.

(27) Huangfu, L.; Li, R.; Huang, Y.; Wang, S. The IL-17 Family in Diseases: From Bench to Bedside. Sig Transduct Target Ther 2023, 8 (1), 1–22. 10.1038/s41392-023-01620-3.

(28) Liu, S. Structural Insights into the Interleukin-17 Family Cytokines and Their Receptors. In Structural Immunology; Jin, T., Yin, Q., Eds.; Advances in Experimental Medicine and Biology; Springer Singapore: Singapore, 2019; Vol. 1172, pp 97–117. 10.1007/978-981-13-9367-9_5.

(29) Abdel-Moneim, A.; Bakery, H. H.; Allam, G. The Potential Pathogenic Role of IL-17/Th17 Cells in Both Type 1 and Type 2 Diabetes Mellitus. Biomedicine & Pharmacotherapy 2018, 101, 287–292. 10.1016/j.biopha.2018.02.103.

(30) Monin, L.; Gaffen, S. L. Interleukin 17 Family Cytokines: Signaling Mechanisms, Biological Activities, and Therapeutic Implications. Cold Spring Harb Perspect Biol 2018, 10 (4), a028522. 10.1101/cshperspect.a028522.

(31) Khalid, F.; Takagi, K.; Sato, A.; Yamaguchi, M.; Guestini, F.; Miki, Y.; Miyashita, M.; Hirakawa, H.; Ohi, Y.; Rai, Y.; Sagara, Y.; Sasano, H.; Suzuki, T. Interleukin (IL)-17A in Triple-Negative Breast Cancer: A Potent Prognostic Factor Associated with Intratumoral Neutrophil Infiltration. Breast Cancer 2023, 30 (5), 748–757. 10.1007/s12282-023-01467-0.

(32) Gu, F.-M.; Li, Q.-L.; Gao, Q.; Jiang, J.-H.; Zhu, K.; Huang, X.-Y.; Pan, J.-F.; Yan, J.; Hu, J.-H.; Wang, Z.; Dai, Z.; Fan, J.; Zhou, J. IL-17 Induces AKT-Dependent IL-6/JAK2/STAT3 Activation and Tumor Progression in Hepatocellular Carcinoma. Molecular Cancer 2011, 10 (1), 150. 10.1186/1476-4598-10-150.

(33) Oláh, J.; Szénási, T.; Lehotzky, A.; Norris, V.; Ovádi, J. Challenges in Discovering Drugs That Target the Protein–Protein Interactions of Disordered Proteins. Int J Mol Sci 2022, 23 (3), 1550. 10.3390/ijms23031550.

(34) Lu, H.; Zhou, Q.; He, J.; Jiang, Z.; Peng, C.; Tong, R.; Shi, J. Recent Advances in the Development of Protein–Protein Interactions Modulators: Mechanisms and Clinical Trials. Sig Transduct Target Ther 2020, 5 (1), 213. 10.1038/s41392-020-00315-3.

(35) Nada, H.; Choi, Y.; Kim, S.; Jeong, K. S.; Meanwell, N. A.; Lee, K. New Insights into Protein–Protein Interaction Modulators in Drug Discovery and Therapeutic Advance. Sig Transduct Target Ther 2024, 9 (1), 1–32. 10.1038/s41392-024-02036-3.

(36) Croft, M.; Salek-Ardakani, S.; Ware, C. F. Targeting the TNF and TNFR Superfamilies in Autoimmune Disease and Cancer. Nat Rev Drug Discov 2024, 23 (12), 939–961. 10.1038/s41573-024-01053-9.

(37) Melsheimer, R.; Geldhof, A.; Apaolaza, I.; Schaible, T. Remicade®(Infliximab): 20 Years of Contributions to Science and Medicine. Biologics: Targets and Therapy 2019, 139–178.

(38) Ellis, C. R.; Azmat, C. E. Adalimumab. In StatPearls; StatPearls Publishing: Treasure Island (FL), 2024.

(39) Nada, H.; Sivaraman, A.; Lu, Q.; Min, K.; Kim, S.; Goo, J.-I.; Choi, Y.; Lee, K. Perspective for Discovery of Small Molecule IL-6 Inhibitors through Study of Structure–Activity Relationships and Molecular Docking. J. Med. Chem. 2023, 66 (7), 4417–4433. 10.1021/acs.jmedchem.2c01957.

(40) Mullard, A. 2014 FDA Drug Approvals: The FDA Approved 41 New Therapeutics in 2014, but the Bumper Year Fell Short of the Commercial Power of the Drugs Approved in 2013. Nature Reviews Drug Discovery 2015, 14 (2), 77–82.

(41) Bartoli, F.; Bae, S.; Cometi, L.; Matucci Cerinic, M.; Furst, D. E. Sirukumab for the Treatment of Rheumatoid Arthritis: Update on Sirukumab, 2018. Expert Rev Clin Immunol 2018, 14 (7), 539–547. 10.1080/1744666X.2018.1487291.

(42) Mospan, G.; Mospan, C.; Vance, S.; Bradshaw, A.; Meosky, K.; Bowles, K. Drug Updates and Approvals: 2017 in Review. The Nurse Practitioner 2017, 42 (12), 8. 10.1097/01.NPR.0000526760.22854.f1.

(43) Heo, Y.-A. Satralizumab: First Approval. Drugs 2020, 80 (14), 1477–1482. 10.1007/s40265-020-01380-2.

(44) Scott, L. J. Tocilizumab: A Review in Rheumatoid Arthritis. Drugs 2017, 77 (17), 1865–1879. 10.1007/s40265-017-0829-7.

(45) Blegvad, C.; Skov, L.; Zachariae, C. Ixekizumab for the Treatment of Psoriasis: An Update on New Data since First Approval. Expert Rev Clin Immunol 2019, 15 (2), 111–121. 10.1080/1744666X.2019.1559730.

(46) Commissioner, O. of the. FDA approves new psoriasis drug Taltz. FDA. https://www.fda.gov/news-events/press-announcements/fda-approves-new-psoriasis-drug-taltz (accessed 2024-10-21).

(47) Roman, M.; Chiu, M. W. Spotlight on Brodalumab in the Treatment of Moderate-to-Severe Plaque Psoriasis: Design, Development, and Potential Place in Therapy. *Drug Design*, Development and Therapy 2017, 11, 2065. 10.2147/DDDT.S113683.

(48) Fala, L. Nucala (Mepolizumab): First IL-5 Antagonist Monoclonal Antibody FDA Approved for Maintenance Treatment of Patients with Severe Asthma. American Health & Drug Benefits 2016, 9 (Spec Feature), 106.

(49) Commissioner, O. of the. FDA approves first drug for Eosinophilic Granulomatosis with Polyangiitis, a rare disease formerly known as the Churg-Strauss Syndrome. FDA. https://www.fda.gov/news-events/press-announcements/fda-approves-first-drug-eosinophilic-granulomatosis-polyangiitis-rare-disease-formerly-known-churg (accessed 2024-10-21).

(50) Fasenra approved in the US for eosinophilic granulomatosis with polyangiitis. https://www.astrazeneca.com/media-centre/press-releases/2024/fasenra-approved-in-the-us-for-eosinophilic-granulomatosis-with-polyangiitis.html (accessed 2024-10-21).

(51) Castro, M.; Corren, J.; Pavord, I. D.; Maspero, J.; Wenzel, S.; Rabe, K. F.; Busse, W. W.; Ford, L.; Sher, L.; FitzGerald, J. M.; Katelaris, C.; Tohda, Y.; Zhang, B.; Staudinger, H.; Pirozzi, G.; Amin, N.; Ruddy, M.; Akinlade, B.; Khan, A.; Chao, J.; Martincova, R.; Graham, N. M. H.; Hamilton, J. D.; Swanson, B. N.; Stahl, N.; Yancopoulos, G. D.; Teper, A. Dupilumab Efficacy and Safety in Moderate-to-Severe Uncontrolled Asthma. N Engl J Med 2018, 378 (26), 2486–2496. 10.1056/NEJMoa1804092.

(52) Galletti, C.; Barbieri, M. A.; Ciodaro, F.; Freni, F.; Galletti, F.; Spina, E.; Galletti, B. Effectiveness and Safety Profile of Dupilumab in Chronic Rhinosinusitis with Nasal Polyps: Real-Life Data in Tertiary Care. Pharmaceuticals 2023, 16 (4), 630. 10.3390/ph16040630.

(53) Grey, A.; Katelaris, C. H. Dupilumab in the Treatment of Asthma. Immunotherapy 2019, 11 (10), 859–872. 10.2217/imt-2019-0008.

(54) FDA Approves Lilly’s EBGLYSS^TM^ (lebrikizumab-lbkz) for Adults and Children 12 Years and Older with Moderate-to-Severe Atopic Dermatitis | Eli Lilly and Company. https://investor.lilly.com/news-releases/news-release-details/fda-approves-lillys-ebglysstm-lebrikizumab-lbkz-adults-and (accessed 2024-10-21).

(55) FDA Approves Lebrikizumab Treatment of Eczema Among Patients Aged 12 and Older. Dermatology | GW School of Medicine and Health Sciences. https://dermatology.smhs.gwu.edu/news/fda-approves-lebrikizumab-treatment-eczema-among-patients-aged-12-and-older (accessed 2024-10-21).

(56) Association, N. E. LEO Pharma Inc. Announces U.S. FDA approval of Adbry® (tralokinumab-ldrm) for the Treatment of Moderate-to-severe Atopic Dermatitis in Pediatric Patients Aged 12-17 Years. National Eczema Association. https://nationaleczema.org/blog/leo-121523/ (accessed 2024-10-21).

(57) Autoinflammatory Disease Treatment | ILARIS® (canakinumab). https://www.ilaris.com/?site=415726-415729GK100009&utm_source=google&utm_mlr=415726-415729&utm_medium=cpc&utm_campaign=google_branded_415726-415729-ilaris-dtc-branded%3Bs%3Bph%3Bbr%3Bimm%3Bdtc%3Bbr_apr-2024&utm_content=ila_aosd_pfs_sjia_brandaware_n2_brand-general-exact&utm_term=canakinumab&gclid=CjwKCAjwg8qzBhAoEiwAWagLrDdeYFdjnxCYcIPkeTPa56jDExW7-hbn0V-XBAMAPuCOQkEPRXRbfxoCtkUQAvD_BwE&gclsrc=aw.ds (accessed 2024-06-19).

(58) Colquhoun, M.; Kemp, A. K. Ustekinumab. In StatPearls; StatPearls Publishing: Treasure Island (FL), 2024.

(59) Dmytrijuk, A.; Robie-Suh, K.; Cohen, M. H.; Rieves, D.; Weiss, K.; Pazdur, R. FDA Report: Eculizumab (Soliris) for the Treatment of Patients with Paroxysmal Nocturnal Hemoglobinuria. Oncologist 2008, 13 (9), 993–1000. 10.1634/theoncologist.2008-0086.

(60) Blair, H. A. Spesolimab: First Approval. Drugs 2022, 82 (17), 1681. 10.1007/s40265-022-01801-4.

(61) Research, C. for D. E. and. FDA Approves Emapalumab for Hemophagocytic Lymphohistiocytosis. FDA 2019.

(62) Burki, T. K. FDA Approval for Anifrolumab in Patients with Lupus. The Lancet Rheumatology 2021, 3 (10), e689. 10.1016/S2665-9913(21)00291-5.

(63) FASENRA approved for treatment of children aged 6 to 11 with severe asthma. https://www.astrazeneca-us.com/media/press-releases/2024/fasenra-approved-for-treatment-of-children-aged-6-to-11-with-severe-asthma.html (accessed 2024-10-21).

(64) Pan, A.; Gerriets, V. Etanercept. In StatPearls; StatPearls Publishing: Treasure Island (FL), 2025.

(65) KINERET® (anakinra) Official Website | Home. https://www.kineretrx.com/ (accessed 2025-11-28).

(66) TREMFYA® (guselkumab). https://www.tremfya.com/?utm_source=google&utm_medium=cpc&utm_campaign=EG-DTCB-BR-NA-Tremfya+Core-JJ-Core-Exact-NA&utm_content=Core-TXT-National-NA-1-EX&utm_term=tremfya&gad_source=1&gad_campaignid=22056523649&gbraid=0AAAAAq4HexIhzpyaKeo1WjGD9tfAv3cUQ&gclid=CjwKCAiAraXJBhBJEiwAjz7MZQKH20cuIlpGcf39ZSPgJM7BgP8s1Fh28OH0OT_zuLSwA8WVlBzyURoCHIEQAvD_BwE&gclsrc=aw.ds (accessed 2025-11-28).

(67) Learn more about SKYRIZI® (risankizumabJrzaa). https://skyrizi.com/ (accessed 2025-11-28).

(68) Plaque Psoriasis Treatment. ILUMYA® (tildrakizumab-asmn). https://www.ilumya.com (accessed 2025-11-28).

(69) Castelli, M. S.; McGonigle, P.; Hornby, P. J. The Pharmacology and Therapeutic Applications of Monoclonal Antibodies. Pharmacology Res & Perspec 2019, 7 (6), e00535. 10.1002/prp2.535.

(70) Silberstein, S.; Lenz, R.; Xu, C. Therapeutic Monoclonal Antibodies: What Headache Specialists Need to Know. Headache 2015, 55 (8), 1171–1182. 10.1111/head.12642.

(71) Mahalingaiah, P. K.; Ciurlionis, R.; Durbin, K. R.; Yeager, R. L.; Philip, B. K.; Bawa, B.; Mantena, S. R.; Enright, B. P.; Liguori, M. J.; Van Vleet, T. R. Potential Mechanisms of Target-Independent Uptake and Toxicity of Antibody-Drug Conjugates. Pharmacology & Therapeutics 2019, 200, 110–125. 10.1016/j.pharmthera.2019.04.008.

(72) Catapano, A. L.; Papadopoulos, N. The Safety of Therapeutic Monoclonal Antibodies: Implications for Cardiovascular Disease and Targeting the PCSK9 Pathway. Atherosclerosis 2013, 228 (1), 18–28. 10.1016/j.atherosclerosis.2013.01.044.

(73) Giblin, K. A.; Basili, D.; Afzal, A. M.; Rosenbrier-Ribeiro, L.; Greene, N.; Barrett, I.; Hughes, S. J.; Bender, A. New Associations between Drug-Induced Adverse Events in Animal Models and Humans Reveal Novel Candidate Safety Targets. Chem. Res. Toxicol. 2021, 34 (2), 438–451. 10.1021/acs.chemrestox.0c00311.

(74) Gómez-Mantilla, J. D.; Trocóniz, I. F.; Parra-Guillén, Z.; Garrido, M. J. Review on Modeling Anti-Antibody Responses to Monoclonal Antibodies. J Pharmacokinet Pharmacodyn 2014, 41 (5), 523–536. 10.1007/s10928-014-9367-z.

(75) Gilardi, D.; Gabbiadini, R.; Allocca, M.; Correale, C.; Fiorino, G.; Furfaro, F.; Zilli, A.; Peyrin-Biroulet, L.; Danese, S. PK, PD, and Interactions: The New Scenario with JAK Inhibitors and S1P Receptor Modulators, Two Classes of Small Molecule Drugs, in IBD. Expert Review of Gastroenterology & Hepatology 2020, 14 (9), 797–806. 10.1080/17474124.2020.1785868.

(76) Diao, L.; Meibohm, B. Pharmacokinetics and Pharmacokinetic–Pharmacodynamic Correlations of Therapeutic Peptides. Clin Pharmacokinet 2013, 52 (10), 855–868. 10.1007/s40262-013-0079-0.

(77) Zheng, J.; Chen, D.; Xu, J.; Ding, X.; Wu, Y.; Shen, H. C.; Tan, X. Small Molecule Approaches to Treat Autoimmune and Inflammatory Diseases (Part III): Targeting Cytokines and Cytokine Receptor Complexes. Bioorganic & Medicinal Chemistry Letters 2021, 48, 128229. 10.1016/j.bmcl.2021.128229.

(78) Deng, R.; Jin, F.; Prabhu, S.; Iyer, S. Monoclonal Antibodies: What Are the Pharmacokinetic and Pharmacodynamic Considerations for Drug Development? Expert Opinion on Drug Metabolism & Toxicology 2012, 8 (2), 141–160. 10.1517/17425255.2012.643868.

(79) Singh, A. P.; Shin, Y. G.; Shah, D. K. Application of Pharmacokinetic-Pharmacodynamic Modeling and Simulation for Antibody-Drug Conjugate Development. Pharm Res 2015, 32 (11), 3508–3525. 10.1007/s11095-015-1626-1.

(80) Dostalek, M.; Gardner, I.; Gurbaxani, B. M.; Rose, R. H.; Chetty, M. Pharmacokinetics, Pharmacodynamics and Physiologically-Based Pharmacokinetic Modelling of Monoclonal Antibodies. Clin Pharmacokinet 2013, 52 (2), 83–124. 10.1007/s40262-012-0027-4.

(81) Büttel, I. C.; Chamberlain, P.; Chowers, Y.; Ehmann, F.; Greinacher, A.; Jefferis, R.; Kramer, D.; Kropshofer, H.; Lloyd, P.; Lubiniecki, A.; Krause, R.; Mire-Sluis, A.; Platts-Mills, T.; Ragheb, J. A.; Reipert, B. M.; Schellekens, H.; Seitz, R.; Stas, P.; Subramanyam, M.; Thorpe, R.; Trouvin, J.-H.; Weise, M.; Windisch, J.; Schneider, C. K. Taking Immunogenicity Assessment of Therapeutic Proteins to the next Level. Biologicals 2011, 39 (2), 100–109. 10.1016/j.biologicals.2011.01.006.

(82) Di, L. Strategic Approaches to Optimizing Peptide ADME Properties. AAPS J 2015, 17 (1), 134–143. 10.1208/s12248-014-9687-3.

(83) Makurvet, F. D. Biologics vs. Small Molecules: Drug Costs and Patient Access. Medicine in Drug Discovery 2021, 9, 100075. 10.1016/j.medidd.2020.100075.

(84) Hommel, U.; Hurth, K.; Rondeau, J.-M.; Vulpetti, A.; Ostermeier, D.; Boettcher, A.; Brady, J. P.; Hediger, M.; Lehmann, S.; Koch, E.; Blechschmidt, A.; Yamamoto, R.; Tundo Dottorello, V.; Haenni-Holzinger, S.; Kaiser, C.; Lehr, P.; Lingel, A.; Mureddu, L.; Schleberger, C.; Blank, J.; Ramage, P.; Freuler, F.; Eder, J.; Bornancin, F. Discovery of a Selective and Biologically Active Low-Molecular Weight Antagonist of Human Interleukin-1β. Nat Commun 2023, 14, 5497. 10.1038/s41467-023-41190-0.

(85) Vulpetti, A.; Rondeau, J.-M.; Bellance, M.-H.; Blank, J.; Boesch, R.; Boettcher, A.; Bornancin, F.; Buhr, S.; Connor, L. E.; Dumelin, C. E.; Esser, O.; Hediger, M.; Hintermann, S.; Hommel, U.; Koch, E.; Lapointe, G.; Leder, L.; Lehmann, S.; Lehr, P.; Meier, P.; Muller, L.; Ostermeier, D.; Ramage, P.; Schiebel-Haddad, S.; Smith, A. B.; Stojanovic, A.; Velcicky, J.; Yamamoto, R.; Hurth, K. Ligandability Assessment of IL-1β by Integrated Hit Identification Approaches. J. Med. Chem. 2024. 10.1021/acs.jmedchem.4c00240.

(86) Krumm, B.; Meng, X.; Xiang, Y.; Deng, J. Identification of Small Molecule Inhibitors of Interleukin-18. Sci Rep 2017, 7 (1), 483. 10.1038/s41598-017-00532-x.

(87) Günther, S.; Sundberg, E. J. Molecular Determinants of Agonist and Antagonist Signaling through the IL-36 Receptor. J Immunol 2014, 193 (2), 921–930. 10.4049/jimmunol.1400538.

(88) Todorović, V.; Su, Z.; Putman, C. B.; Kakavas, S. J.; Salte, K. M.; McDonald, H. A.; Wetter, J. B.; Paulsboe, S. E.; Sun, Q.; Gerstein, C. E.; Medina, L.; Sielaff, B.; Sadhukhan, R.; Stockmann, H.; Richardson, P. L.; Qiu, W.; Argiriadi, M. A.; Henry, R. F.; Herold, J. M.; Shotwell, J. B.; McGaraughty, S. P.; Honore, P.; Gopalakrishnan, S. M.; Sun, C. C.; Scott, V. E. Small Molecule IL-36γ Antagonist as a Novel Therapeutic Approach for Plaque Psoriasis. Sci Rep 2019, 9 (1), 9089. 10.1038/s41598-019-45626-w.

(89) Senter, P. D.; Al-Abed, Y.; Metz, C. N.; Benigni, F.; Mitchell, R. A.; Chesney, J.; Han, J.; Gartner, C. G.; Nelson, S. D.; Todaro, G. J.; Bucala, R. Inhibition of Macrophage Migration Inhibitory Factor (MIF) Tautomerase and Biological Activities by Acetaminophen Metabolites. Proc Natl Acad Sci U S A 2002, 99 (1), 144–149. 10.1073/pnas.011569399.

(90) Lubetsky, J. B.; Dios, A.; Han, J.; Aljabari, B.; Ruzsicska, B.; Mitchell, R.; Lolis, E.; Al-Abed, Y. The Tautomerase Active Site of Macrophage Migration Inhibitory Factor Is a Potential Target for Discovery of Novel Anti-Inflammatory Agents *. Journal of Biological Chemistry 2002, 277 (28), 24976–24982. 10.1074/jbc.M203220200.

(91) Sun, H. W.; Bernhagen, J.; Bucala, R.; Lolis, E. Crystal Structure at 2.6-A Resolution of Human Macrophage Migration Inhibitory Factor. Proc Natl Acad Sci U S A 1996, 93 (11), 5191–5196. 10.1073/pnas.93.11.5191.

(92) Winner, M.; Meier, J.; Zierow, S.; Rendon, B. E.; Crichlow, G. V.; Riggs, R.; Bucala, R.; Leng, L.; Smith, N.; Lolis, E.; Trent, J. O.; Mitchell, R. A. A Novel, Macrophage Migration Inhibitory Factor Suicide Substrate Inhibits Motility and Growth of Lung Cancer Cells. Cancer Res 2008, 68 (18), 7253–7257. 10.1158/0008-5472.CAN-07-6227.

(93) Rajasekaran, D.; Zierow, S.; Syed, M.; Bucala, R.; Bhandari, V.; Lolis, E. J. Targeting Distinct Tautomerase Sites of D-DT and MIF with a Single Molecule for Inhibition of Neutrophil Lung Recruitment. FASEB J 2014, 28 (11), 4961–4971. 10.1096/fj.14-256636.

(94) Imaoka, M.; Tanese, K.; Masugi, Y.; Hayashi, M.; Sakamoto, M. Macrophage Migration Inhibitory Factor-CD74 Interaction Regulates the Expression of Programmed Cell Death Ligand 1 in Melanoma Cells. Cancer Sci 2019, 110 (7), 2273–2283. 10.1111/cas.14038.

(95) Ouertatani-Sakouhi, H.; El-Turk, F.; Fauvet, B.; Roger, T.; Le Roy, D.; Karpinar, D. P.; Leng, L.; Bucala, R.; Zweckstetter, M.; Calandra, T.; Lashuel, H. A. A New Class of Isothiocyanate-Based Irreversible Inhibitors of Macrophage Migration Inhibitory Factor (MIF). Biochemistry 2009, 48 (41), 9858–9870. 10.1021/bi900957e.

(96) Mawhinney, L.; Armstrong, M. E.; O’ Reilly, C.; Bucala, R.; Leng, L.; Fingerle-Rowson, G.; Fayne, D.; Keane, M. P.; Tynan, A.; Maher, L.; Cooke, G.; Lloyd, D.; Conroy, H.; Donnelly, S. C. Macrophage Migration Inhibitory Factor (MIF) Enzymatic Activity and Lung Cancer. Mol Med 2015, 20 (1), 729–735. 10.2119/molmed.2014.00136.

(97) Ouertatani-Sakouhi, H.; El-Turk, F.; Fauvet, B.; Cho, M.-K.; Pinar Karpinar, D.; Le Roy, D.; Dewor, M.; Roger, T.; Bernhagen, J.; Calandra, T.; Zweckstetter, M.; Lashuel, H. A. Identification and Characterization of Novel Classes of Macrophage Migration Inhibitory Factor (MIF) Inhibitors with Distinct Mechanisms of Action. J Biol Chem 2010, 285 (34), 26581–26598. 10.1074/jbc.M110.113951.

(98) Bai, F.; Asojo, O. A.; Cirillo, P.; Ciustea, M.; Ledizet, M.; Aristoff, P. A.; Leng, L.; Koski, R. A.; Powell, T. J.; Bucala, R.; Anthony, K. G. A Novel Allosteric Inhibitor of Macrophage Migration Inhibitory Factor (MIF). J Biol Chem 2012, 287 (36), 30653–30663. 10.1074/jbc.M112.385583.

(99) Cirillo, P. F.; Asojo, O. A.; Khire, U.; Lee, Y.; Mootien, S.; Hegan, P.; Sutherland, A. G.; Peterson-Roth, E.; Ledizet, M.; Koski, R. A.; Anthony, K. G. Inhibition of Macrophage Migration Inhibitory Factor by a Chimera of Two Allosteric Binders. ACS Med. Chem. Lett. 2020, 11 (10), 1843–1847. 10.1021/acsmedchemlett.9b00351.

(100) Bloom, J.; Pantouris, G.; He, M.; Aljabari, B.; Mishra, L.; Manjula, R.; Parkins, A.; Lolis, E. J.; Al-Abed, Y. Iguratimod, an Allosteric Inhibitor of Macrophage Migration Inhibitory Factor (MIF), Prevents Mortality and Oxidative Stress in a Murine Model of Acetaminophen Overdose. Molecular Medicine 2024, 30 (1), 43. 10.1186/s10020-024-00803-0.

(101) Sumaiya, K.; Selvambika, P.; Natarajaseenivasan, K. Anti-Macrophage Migration Inhibitory Factor (MIF) Activity of Ibudilast: A Repurposing Drug Attenuates the Pathophysiology of Leptospirosis. Microbial Pathogenesis 2022, 173, 105786. 10.1016/j.micpath.2022.105786.

(102) Tilstam, P. V.; Pantouris, G.; Corman, M.; Andreoli, M.; Mahboubi, K.; Davis, G.; Du, X.; Leng, L.; Lolis, E.; Bucala, R. A Selective Small-Molecule Inhibitor of Macrophage Migration Inhibitory Factor-2 (MIF-2), a MIF Cytokine Superfamily Member, Inhibits MIF-2 Biological Activity. J Biol Chem 2019, 294 (49), 18522–18531. 10.1074/jbc.RA119.009860.

(103) Zhang, M.; Yang, X.-Y.; Tang, W.; Groeneveld, T. W. L.; He, P.-L.; Zhu, F.-H.; Li, J.; Lu, W.; Blom, A. M.; Zuo, J.-P.; Nan, F.-J. Discovery and Structural Modification of 1-Phenyl-3-(1-Phenylethyl)Urea Derivatives as Inhibitors of Complement. ACS Med Chem Lett 2012, 3 (4), 317–321. 10.1021/ml300005w.

(104) Jendza, K.; Kato, M.; Salcius, M.; Srinivas, H.; De Erkenez, A.; Nguyen, A.; McLaughlin, D.; Be, C.; Wiesmann, C.; Murphy, J.; Bolduc, P.; Mogi, M.; Duca, J.; Namil, A.; Capparelli, M.; Darsigny, V.; Meredith, E.; Tichkule, R.; Ferrara, L.; Heyder, J.; Liu, F.; Horton, P. A.; Romanowski, M. J.; Schirle, M.; Mainolfi, N.; Anderson, K.; Michaud, G. A. A Small-Molecule Inhibitor of C5 Complement Protein. Nat Chem Biol 2019, 15 (7), 666–668. 10.1038/s41589-019-0303-9.

(105) Tilley, J. W.; Chen, L.; Fry, D. C.; Emerson, S. D.; Powers, G. D.; Biondi, D.; Varnell, T.; Trilles, R.; Guthrie, R.; Mennona, F.; Kaplan, G.; LeMahieu, R. A.; Carson, M.; Han, R.-J.; Liu, C.-M.; Palermo, R.; Ju, G. Identification of a Small Molecule Inhibitor of the IL-2/IL-2Rα Receptor Interaction Which Binds to IL-2. J. Am. Chem. Soc. 1997, 119 (32), 7589–7590. 10.1021/ja970702x.

(106) Braisted, A. C.; Oslob, J. D.; Delano, W. L.; Hyde, J.; McDowell, R. S.; Waal, N.; Yu, C.; Arkin, M. R.; Raimundo, B. C. Discovery of a Potent Small Molecule IL-2 Inhibitor through Fragment Assembly. J. Am. Chem. Soc. 2003, 125 (13), 3714–3715. 10.1021/ja034247i.

(107) Thanos, C. D.; Randal, M.; Wells, J. A. Potent Small-Molecule Binding to a Dynamic Hot Spot on IL-2. J. Am. Chem. Soc. 2003, 125 (50), 15280–15281. 10.1021/ja0382617.

(108) Quinnell, S. P.; Leifer, B. S.; Nestor, S. T.; Tan, K.; Sheehy, D. F.; Ceo, L.; Doyle, S. K.; Koehler, A. N.; Vegas, A. J. A Small-Molecule Inhibitor to the Cytokine Interleukin-4. ACS Chem. Biol. 2020, 15 (10), 2649–2654. 10.1021/acschembio.0c00615.

(109) Quéméner, A.; Maillasson, M.; Arzel, L.; Sicard, B.; Vomiandry, R.; Mortier, E.; Dubreuil, D.; Jacques, Y.; Lebreton, J.; Mathé-Allainmat, M. Discovery of a Small-Molecule Inhibitor of Interleukin 15: Pharmacophore-Based Virtual Screening and Hit Optimization. J. Med. Chem. 2017, 60 (14), 6249–6272. 10.1021/acs.jmedchem.7b00485.

(110) Thoidingjam, L. K.; Blouin, C. M.; Gaillet, C.; Brion, A.; Solier, S.; Niyomchon, S.; El Marjou, A.; Mouasni, S.; Sepulveda, F. E.; de Saint Basile, G.; Lamaze, C.; Rodriguez, R. Small Molecule Inhibitors of Interferon-Induced JAK-STAT Signalling. Angew Chem Int Ed Engl 2022, 61 (32), e202205231. 10.1002/anie.202205231.

(111) Imai, K.; Takaoka, A. Comparing Antibody and Small-Molecule Therapies for Cancer. Nat Rev Cancer 2006, 6 (9), 714–727. 10.1038/nrc1913.

(112) Hoelder, S.; Clarke, P. A.; Workman, P. Discovery of Small Molecule Cancer Drugs: Successes, Challenges and Opportunities. Mol Oncol 2012, 6 (2), 155–176. 10.1016/j.molonc.2012.02.004.

(113) Boskovic, Z. V.; Kemp, M. M.; Freedy, A. M.; Viswanathan, V. S.; Pop, M. S.; Fuller, J. H.; Martinez, N. M.; Figueroa Lazú, S. O.; Hong, J. A.; Lewis, T. A.; Calarese, D.; Love, J. D.; Vetere, A.; Almo, S. C.; Schreiber, S. L.; Koehler, A. N. Inhibition of Zinc-Dependent Histone Deacetylases with a Chemically Triggered Electrophile. ACS Chem. Biol. 2016, 11 (7), 1844–1851. 10.1021/acschembio.6b00012.

(114) Bradner, J. E.; McPherson, O. M.; Koehler, A. N. A Method for the Covalent Capture and Screening of Diverse Small Molecules in a Microarray Format. Nat Protoc 2006, 1 (5), 2344–2352. 10.1038/nprot.2006.282.

(115) Bradner, J. E.; McPherson, O. M.; Mazitschek, R.; Barnes-Seeman, D.; Shen, J. P.; Dhaliwal, J.; Stevenson, K. E.; Duffner, J. L.; Park, S. B.; Neuberg, D. S.; Nghiem, P.; Schreiber, S. L.; Koehler, A. N. A Robust Small-Molecule Microarray Platform for Screening Cell Lysates. Chem Biol 2006, 13 (5), 493–504. 10.1016/j.chembiol.2006.03.004.

(116) Kemp, M. M.; Wang, Q.; Fuller, J. H.; West, N.; Martinez, N. M.; Morse, E. M.; Weïwer, M.; Schreiber, S. L.; Bradner, J. E.; Koehler, A. N. A Novel HDAC Inhibitor with a Hydroxy-Pyrimidine Scaffold. Bioorganic & Medicinal Chemistry Letters 2011, 21 (14), 4164–4169. 10.1016/j.bmcl.2011.05.098.

(117) Stanton, B. Z.; Peng, L. F.; Maloof, N.; Nakai, K.; Wang, X.; Duffner, J. L.; Taveras, K. M.; Hyman, J. M.; Lee, S. W.; Koehler, A. N.; Chen, J. K.; Fox, J. L.; Mandinova, A.; Schreiber, S. L. A Small Molecule That Binds Hedgehog and Blocks Its Signaling in Human Cells. Nat Chem Biol 2009, 5 (3), 154–156. 10.1038/nchembio.142.

(118) Vegas, A. J.; Bradner, J. E.; Tang, W.; McPherson, O. M.; Greenberg, E. F.; Koehler, A. N.; Schreiber, S. L. Fluorous-Based Small-Molecule Microarrays for the Discovery of Histone Deacetylase Inhibitors. Angew Chem Int Ed Engl 2007, 46 (42), 7960–7964. 10.1002/anie.200703198.

(119) Chen, J.; Armstrong, A. H.; Koehler, A. N.; Hecht, M. H. Small Molecule Microarrays Enable the Discovery of Compounds That Bind the Alzheimer’s Aβ Peptide and Reduce Its Cytotoxicity. J Am Chem Soc 2010, 132 (47), 17015–17022. 10.1021/ja107552s.

(120) Struntz, N. B.; Chen, A.; Deutzmann, A.; Wilson, R. M.; Stefan, E.; Evans, H. L.; Ramirez, M. A.; Liang, T.; Caballero, F.; Wildschut, M. H. E.; Neel, D. V.; Freeman, D. B.; Pop, M. S.; McConkey, M.; Muller, S.; Curtin, B. H.; Tseng, H.; Frombach, K. R.; Butty, V. L.; Levine, S. S.; Feau, C.; Elmiligy, S.; Hong, J. A.; Lewis, T. A.; Vetere, A.; Clemons, P. A.; Malstrom, S. E.; Ebert, B. L.; Lin, C. Y.; Felsher, D. W.; Koehler, A. N. Stabilization of the Max Homodimer with a Small Molecule Attenuates Myc-Driven Transcription. Cell Chem Biol 2019, 26 (5), 711–723.e14. 10.1016/j.chembiol.2019.02.009.

(121) Seiler, K. P.; George, G. A.; Happ, M. P.; Bodycombe, N. E.; Carrinski, H. A.; Norton, S.; Brudz, S.; Sullivan, J. P.; Muhlich, J.; Serrano, M.; Ferraiolo, P.; Tolliday, N. J.; Schreiber, S. L.; Clemons, P. A. ChemBank: A Small-Molecule Screening and Cheminformatics Resource Database. Nucleic Acids Res 2008, 36 (Database issue), D351-359. 10.1093/nar/gkm843.

(122) Clemons, P. A.; Bodycombe, N. E.; Carrinski, H. A.; Wilson, J. A.; Shamji, A. F.; Wagner, B. K.; Koehler, A. N.; Schreiber, S. L. Small Molecules of Different Origins Have Distinct Distributions of Structural Complexity That Correlate with Protein-Binding Profiles. Proc Natl Acad Sci U S A 2010, 107 (44), 18787–18792. 10.1073/pnas.1012741107.

(123) Mantovani, A.; Dinarello, C. A.; Molgora, M.; Garlanda, C. IL-1 and Related Cytokines in Innate and Adaptive Immunity in Health and Disease. Immunity 2019, 50 (4), 778–795. 10.1016/j.immuni.2019.03.012.

(124) Boraschi, D.; Tagliabue, A. The Interleukin-1 Receptor Family. Seminars in Immunology 2013, 25 (6), 394–407. 10.1016/j.smim.2013.10.023.

(125) Zhou, L.; Todorovic, V. Interleukin-36: Structure, Signaling and Function. Adv Exp Med Biol 2021, 21, 191–210. 10.1007/5584_2020_488.

(126) Fields, J. K.; Günther, S.; Sundberg, E. J. Structural Basis of IL-1 Family Cytokine Signaling. Front Immunol 2019, 10, 1412. 10.3389/fimmu.2019.01412.

(127) Günther, S.; Deredge, D.; Bowers, A. L.; Luchini, A.; Bonsor, D. A.; Beadenkopf, R.; Liotta, L.; Wintrode, P. L.; Sundberg, E. J. IL-1 Family Cytokines Use Distinct Molecular Mechanisms to Signal through Their Shared Co-Receptor. Immunity 2017, 47 (3), 510–523.e4. 10.1016/j.immuni.2017.08.004.

(128) Heguy, A.; Baldari, C. T.; Macchia, G.; Telford, J. L.; Melli, M. Amino Acids Conserved in Interleukin-1 Receptors (IL-1Rs) and the Drosophila Toll Protein Are Essential for IL-1R Signal Transduction. J Biol Chem 1992, 267 (4), 2605–2609.

(129) Gay, N. J.; Keith, F. J. Drosophila Toll and IL-1 Receptor. Nature 1991, 351 (6325), 355–356. 10.1038/351355b0.

(130) Dunne, A.; O’Neill, L. A. J. The Interleukin-1 Receptor/Toll-Like Receptor Superfamily: Signal Transduction During Inflammation and Host Defense. Science’s STKE 2003, 2003 (171), re3–re3. 10.1126/stke.2003.171.re3.

(131) Cohen, P. The TLR and IL-1 Signalling Network at a Glance. J Cell Sci 2014, 127 (11), 2383–2390. 10.1242/jcs.149831.

(132) Wang, X.; Lupardus, P.; LaPorte, S. L.; Garcia, K. C. Structural Biology of Shared Cytokine Receptors. Annual review of immunology 2009, 27, 29. 10.1146/annurev.immunol.24.021605.090616.

(133) Ferrao, R.; Wallweber, H. J. A.; Ho, H.; Tam, C.; Franke, Y.; Quinn, J.; Lupardus, P. J. The Structural Basis for Class II Cytokine Receptor Recognition by JAK1. Structure 2016, 24 (6), 897–905. 10.1016/j.str.2016.03.023.

(134) The JAK/STAT signaling pathway: from bench to clinic - PMC. https://www.ncbi.nlm.nih.gov/pmc/articles/PMC8617206/ (accessed 2024-04-16).

(135) Morris, R.; Kershaw, N. J.; Babon, J. J. The Molecular Details of Cytokine Signaling via the JAK/STAT Pathway. Protein Sci 2018, 27 (12), 1984–2009. 10.1002/pro.3519.

(136) Tang, C.; Chen, S.; Qian, H.; Huang, W. Interleukin-23: As a Drug Target for Autoimmune Inflammatory Diseases. Immunology 2012, 135 (2), 112–124. 10.1111/j.1365-2567.2011.03522.x.

(137) Wojno, E. D. T.; Hunter, C. A.; Stumhofer, J. S. The Immunobiology of the Interleukin-12 Family: Room for Discovery. Immunity 2019, 50 (4), 851–870. 10.1016/j.immuni.2019.03.011.

(138) Langrish, C. L.; McKenzie, B. S.; Wilson, N. J.; de Waal Malefyt, R.; Kastelein, R. A.; Cua, D. J. IL-12 and IL-23: Master Regulators of Innate and Adaptive Immunity. Immunol Rev 2004, 202, 96–105. 10.1111/j.0105-2896.2004.00214.x.

(139) Lyakh, L.; Trinchieri, G.; Provezza, L.; Carra, G.; Gerosa, F. Regulation of Interleukin-12/Interleukin-23 Production and the T-Helper 17 Response in Humans. Immunol Rev 2008, 226, 112–131. 10.1111/j.1600-065X.2008.00700.x.

(140) Glassman, C. R.; Mathiharan, Y. K.; Jude, K. M.; Su, L.; Panova, O.; Lupardus, P. J.; Spangler, J. B.; Ely, L. K.; Thomas, C.; Skiniotis, G.; Garcia, K. C. Structural Basis for IL-12 and IL-23 Receptor Sharing Reveals a Gateway for Shaping Actions on T versus NK Cells. Cell 2021, 184 (4), 983–999.e24. 10.1016/j.cell.2021.01.018.

(141) Fleishaker, D. L.; Garcia Meijide, J. A.; Petrov, A.; Kohen, M. D.; Wang, X.; Menon, S.; Stock, T. C.; Mebus, C. A.; Goodrich, J. M.; Mayer, H. B.; Zeiher, B. G. Maraviroc, a Chemokine Receptor-5 Antagonist, Fails to Demonstrate Efficacy in the Treatment of Patients with Rheumatoid Arthritis in a Randomized, Double-Blind Placebo-Controlled Trial. Arthritis Res Ther 2012, 14 (1), R11. 10.1186/ar3685.

(142) Haringman, J. J.; Gerlag, D. M.; Smeets, T. J. M.; Baeten, D.; Van Den Bosch, F.; Bresnihan, B.; Breedveld, F. C.; Dinant, H. J.; Legay, F.; Gram, H.; Loetscher, P.; Schmouder, R.; Woodworth, T.; Tak, P. P. A Randomized Controlled Trial with an antiDCCL2 (Anti–Monocyte Chemotactic Protein 1) Monoclonal Antibody in Patients with Rheumatoid Arthritis. Arthritis & Rheumatism 2006, 54 (8), 2387–2392. 10.1002/art.21975.

(143) Caorsi, R.; Bertoni, A.; Matucci-Cerinic, C.; Natoli, V.; Palmeri, S.; Rosina, S.; Penco, F.; Malattia, C.; Consolaro, A.; Viola, S.; Papa, R.; Corcione, A.; Volpi, S.; Ravelli, A.; Gattorno, M. Long-Term Efficacy of MAS825, a Bispecific Anti-IL1β and IL-18 Monoclonal Antibody, in Two Patients with Systemic JIA and Recurrent Episodes of Macrophage Activation Syndrome. Rheumatology (Oxford) 2025, 64 (3), 1528–1533. 10.1093/rheumatology/keae440.

(144) Guttman-Yassky, E.; Brunner, P. M.; Neumann, A. U.; Khattri, S.; Pavel, A. B.; Malik, K.; Singer, G. K.; Baum, D.; Gilleaudeau, P.; Sullivan-Whalen, M.; Rose, S.; On, S. J.; Li, X.; Fuentes-Duculan, J.; Estrada, Y.; Garcet, S.; Traidl-Hoffmann, C.; Krueger, J. G.; Lebwohl, M. G. Efficacy and Safety of Fezakinumab (an Anti-IL-22 Monoclonal Antibody) in Adults with Moderate-to-Severe Atopic Dermatitis Inadequately Controlled by Conventional Treatments - A Randomized, Double-Blind, Phase 2a Trial. J Am Acad Dermatol 2018, 78 (5), 872–881.e6. 10.1016/j.jaad.2018.01.016.

(145) Kay, J.; Calabrese, L. The Role of Interleukin-1 in the Pathogenesis of Rheumatoid Arthritis. Rheumatology (Oxford) 2004, 43 *Suppl 3*, iii2–iii9. 10.1093/rheumatology/keh201.

(146) McInnes, I. B.; Liew, F. Y.; Gracie, J. A. Interleukin-18: A Therapeutic Target in Rheumatoid Arthritis? Arthritis Res Ther 2004, 7 (1), 38. 10.1186/ar1497.

(147) Gracie, J. A.; Forsey, R. J.; Chan, W. L.; Gilmour, A.; Leung, B. P.; Greer, M. R.; Kennedy, K.; Carter, R.; Wei, X.-Q.; Xu, D.; Field, M.; Foulis, A.; Liew, F. Y.; McInnes, I. B. A Proinflammatory Role for IL-18 in Rheumatoid Arthritis. J Clin Invest 1999, 104 (10), 1393–1401. 10.1172/JCI7317.

(148) Ehrenstein, M. R.; Evans, J. G.; Singh, A.; Moore, S.; Warnes, G.; Isenberg, D. A.; Mauri, C. Compromised Function of Regulatory T Cells in Rheumatoid Arthritis and Reversal by Anti-TNFα Therapy. Journal of Experimental Medicine 2004, 200 (3), 277–285. 10.1084/jem.20040165.

(149) Maini, R. N.; Taylor, P. C. Anti-Cytokine Therapy for Rheumatoid Arthritis. Annual Review of Medicine 2000, 51 (Volume 51, 2000), 207–229. 10.1146/annurev.med.51.1.207.

(150) Geiler, J.; McDermott, M. F. Gevokizumab, an Anti-IL-1β mAb for the Potential Treatment of Type 1 and 2 Diabetes, Rheumatoid Arthritis and Cardiovascular Disease. Curr Opin Mol Ther 2010, 12 (6), 755–769.

(151) Senolt, L. Emerging Therapies in Rheumatoid Arthritis: Focus on Monoclonal Antibodies. F1000Res 2019, 8, F1000 Faculty Rev-1549. 10.12688/f1000research.18688.1.

(152) Nishimoto, N.; Kishimoto, T. Humanized Antihuman IL-6 Receptor Antibody, Tocilizumab. Handb Exp Pharmacol 2008, No. 181, 151–160. 10.1007/978-3-540-73259-4_7.

(153) Yellin, M.; Paliienko, I.; Balanescu, A.; Ter-Vartanian, S.; Tseluyko, V.; Xu, L.-A.; Tao, X.; Cardarelli, P. M.; Leblanc, H.; Nichol, G.; Ancuta, C.; Chirieac, R.; Luo, A. A Phase II, Randomized, Double-Blind, Placebo-Controlled Study Evaluating the Efficacy and Safety of MDX-1100, a Fully Human Anti-CXCL10 Monoclonal Antibody, in Combination with Methotrexate in Patients with Rheumatoid Arthritis. Arthritis Rheum 2012, 64 (6), 1730–1739. 10.1002/art.34330.

(154) Yuan, N.; Yu, G.; Liu, D.; Wang, X.; Zhao, L. An Emerging Role of Interleukin-23 in Rheumatoid Arthritis. Immunopharmacol Immunotoxicol 2019, 41 (2), 185–191. 10.1080/08923973.2019.1610429.

(155) Smolen, J. S.; Agarwal, S. K.; Ilivanova, E.; Xu, X. L.; Miao, Y.; Zhuang, Y.; Nnane, I.; Radziszewski, W.; Greenspan, A.; Beutler, A.; Baker, D. A Randomised Phase II Study Evaluating the Efficacy and Safety of Subcutaneously Administered Ustekinumab and Guselkumab in Patients with Active Rheumatoid Arthritis despite Treatment with Methotrexate. Annals of the Rheumatic Diseases 2017, 76 (5), 831–839. 10.1136/annrheumdis-2016-209831.

(156) Karnell, J. L.; Albulescu, M.; Drabic, S.; Wang, L.; Moate, R.; Baca, M.; Oganesyan, V.; Gunsior, M.; Thisted, T.; Yan, L.; Li, J.; Xiong, X.; Eck, S. C.; de los Reyes, M.; Yusuf, I.; Streicher, K.; Müller-Ladner, U.; Howe, D.; Ettinger, R.; Herbst, R.; Drappa, J. A CD40L-Targeting Protein Reduces Autoantibodies and Improves Disease Activity in Patients with Autoimmunity. Science Translational Medicine 2019, 11 (489), eaar6584. 10.1126/scitranslmed.aar6584.

(157) Shiomi, A.; Usui, T.; Mimori, T. GM-CSF as a Therapeutic Target in Autoimmune Diseases. Inflamm Regen 2016, 36, 8. 10.1186/s41232-016-0014-5.

(158) Baell, J. B.; Nissink, J. W. M. Seven Year Itch: Pan-Assay Interference Compounds (PAINS) in 2017—Utility and Limitations. ACS Chem. Biol. 2018, 13 (1), 36–44. 10.1021/acschembio.7b00903.

(159) Degen, J.; Wegscheid-Gerlach, C.; Zaliani, A.; Rarey, M. On the Art of Compiling and Using “Drug-Like” Chemical Fragment Spaces. ChemMedChem 2008, 3 (10), 1503–1507. 10.1002/cmdc.200800178.

(160) Lewell, X. Q.; Judd, D. B.; Watson, S. P.; Hann, M. M. RECAPRetrosynthetic Combinatorial Analysis Procedure:D A Powerful New Technique for Identifying Privileged Molecular Fragments with Useful Applications in Combinatorial Chemistry. J. Chem. Inf. Comput. Sci. 1998, 38 (3), 511–522. 10.1021/ci970429i.

(161) Bemis, G. W.; Murcko, M. A. The Properties of Known Drugs. 1. Molecular Frameworks. J. Med. Chem. 1996, 39 (15), 2887–2893. 10.1021/jm9602928.

(162) McInnes, I. B.; Schett, G. Cytokines in the Pathogenesis of Rheumatoid Arthritis. Nat Rev Immunol 2007, 7 (6), 429–442. 10.1038/nri2094.

(163) Segal, B. M.; Constantinescu, C. S.; Raychaudhuri, A.; Kim, L.; Fidelus-Gort, R.; Kasper, L. H.; Ustekinumab MS Investigators. Repeated Subcutaneous Injections of IL12/23 P40 Neutralising Antibody, Ustekinumab, in Patients with Relapsing-Remitting Multiple Sclerosis: A Phase II, Double-Blind, Placebo-Controlled, Randomised, Dose-Ranging Study. Lancet Neurol 2008, 7 (9), 796–804. 10.1016/S1474-4422(08)70173-X.

(164) Bagnasco, D.; Ferrando, M.; Varricchi, G.; Passalacqua, G.; Canonica, G. W. A Critical Evaluation of Anti-IL-13 and Anti-IL-4 Strategies in Severe Asthma. Int Arch Allergy Immunol 2016, 170 (2), 122–131. 10.1159/000447692.

(165) Oh, C. K.; Geba, G. P.; Molfino, N. Investigational Therapeutics Targeting the IL-4/IL-13/STAT-6 Pathway for the Treatment of Asthma. European Respiratory Review 2010, 19 (115), 46–54. 10.1183/09059180.00007609.

(166) Le Floc’h, A.; Allinne, J.; Nagashima, K.; Scott, G.; Birchard, D.; Asrat, S.; Bai, Y.; Lim, W. K.; Martin, J.; Huang, T.; Potocky, T. B.; Kim, J. H.; Rafique, A.; Papadopoulos, N. J.; Stahl, N.; Yancopoulos, G. D.; Murphy, A. J.; Sleeman, M. A.; Orengo, J. M. Dual Blockade of ILD4 and ILD13 with Dupilumab, an ILD4Rα Antibody, Is Required to Broadly Inhibit Type 2 Inflammation. Allergy 2020, 75 (5), 1188–1204. 10.1111/all.14151.

(167) Liu, T.; Li, S.; Ying, S.; Tang, S.; Ding, Y.; Li, Y.; Qiao, J.; Fang, H. The IL-23/IL-17 Pathway in Inflammatory Skin Diseases: From Bench to Bedside. Front. Immunol. 2020, 11. 10.3389/fimmu.2020.594735.

(168) Larsen, J. M.; Bonefeld, C. M.; Poulsen, S. S.; Geisler, C.; Skov, L. IL-23 and TH17-Mediated Inflammation in Human Allergic Contact Dermatitis. Journal of Allergy and Clinical Immunology 2009, 123 (2), 486–492.e1. 10.1016/j.jaci.2008.09.036.

(169) Bunte, K.; Beikler, T. Th17 Cells and the IL-23/IL-17 Axis in the Pathogenesis of Periodontitis and Immune-Mediated Inflammatory Diseases. Int J Mol Sci 2019, 20 (14), 3394. 10.3390/ijms20143394.

(170) Armstrong, A. W.; Papp, K.; Kircik, L. Secukinumab: Review of Clinical Evidence from the Pivotal Studies ERASURE, FIXTURE, and CLEAR. J Clin Aesthet Dermatol 2016, 9 (6 Suppl 1), S7–S12.

(171) Berthold, M. R.; Cebron, N.; Dill, F.; Gabriel, T. R.; Kötter, T.; Meinl, T.; Ohl, P.; Sieb, C.; Thiel, K.; Wiswedel, B. KNIME: The Konstanz Information Miner. In Data Analysis, Machine Learning and Applications; Preisach, C., Burkhardt, H., Schmidt-Thieme, L., Decker, R., Eds.; Springer: Berlin, Heidelberg, 2008; pp 319–326. 10.1007/978-3-540-78246-9_38.

(172) RDKit. https://www.rdkit.org/ (accessed 2025-11-28).

(173) Bajusz, D.; Rácz, A.; Héberger, K. Why Is Tanimoto Index an Appropriate Choice for Fingerprint-Based Similarity Calculations? J Cheminform 2015, 7 (1), 20. 10.1186/s13321-015-0069-3.

(174) Wilson, S. C.; Caveney, N. A.; Yen, M.; Pollmann, C.; Xiang, X.; Jude, K. M.; Hafer, M.; Tsutsumi, N.; Piehler, J.; Garcia, K. C. Organizing Structural Principles of the IL-17 Ligand–Receptor Axis. Nature 2022, 609 (7927), 622–629. 10.1038/s41586-022-05116-y.

(175) Ma, H.; Zhang, W.; Liu, K.; Xu, B.; Li, M.; Meng, Q.; An, Z.; Chen, B. Generation and Characterization of QLS22001, a Humanized Monoclonal Antibody That Neutralizes IL-17A and IL-17F with an Extended Half-Life. International Immunopharmacology 2023, 117, 109947. 10.1016/j.intimp.2023.109947.

(176) Liu, Z.; Song, L.; Yang, J.; Liu, H.; Zhang, Y.; Pi, X.; Yan, Y.; Chen, H.; Yu, D.; Yin, C.; Liu, T.; Li, X.; Zhang, C.; Li, D.; Wang, Z.; Xiao, W. Discovery and Preclinical Evaluation of KYS202004A, a Novel Bispecific Fusion Protein Targeting TNF-α and IL-17A, in Autoimmune Disease Models. International Immunopharmacology 2024, 136, 112383. 10.1016/j.intimp.2024.112383.

(177) Park, A.; Heo, T.-H. Celastrol Regulates Psoriatic Inflammation and Autophagy by Targeting IL-17A. Biomedicine & Pharmacotherapy 2024, 172, 116256. 10.1016/j.biopha.2024.116256.

(178) Berger, S.; Seeger, F.; Yu, T.-Y.; Aydin, M.; Yang, H.; Rosenblum, D.; Guenin-Macé, L.; Glassman, C.; Arguinchona, L.; Sniezek, C.; Blackstone, A.; Carter, L.; Ravichandran, R.; Ahlrichs, M.; Murphy, M.; Pultz, I. S.; Kang, A.; Bera, A. K.; Stewart, L.; Garcia, K. C.; Naik, S.; Spangler, J. B.; Beigel, F.; Siebeck, M.; Gropp, R.; Baker, D. Preclinical Proof of Principle for Orally Delivered Th17 Antagonist Miniproteins. Cell 2024, 187 (16), 4305–4317.e18. 10.1016/j.cell.2024.05.052.

(179) Fatheree, P. R.; Linsell, M. S.; Jacobsen, J. R.; Linden, W. V. der; Church, T. J.; Aquino, C.; Paulick, M. IL-17 Ligands And Uses Thereof. US20200247785A1, August 6, 2020. https://patents.google.com/patent/US20200247785A1/en?oq=Dice+Alpha+Inc.+IL-17+Ligands+And+Uses+Thereof.+US2020247785A1.+2020. (accessed 2024-05-14).

(180) Fatheree, P. R.; Linsell, M. S.; Jacobsen, J. R.; Linden, W. A. V. D.; Church, T. J.; Aquino, C.; Paulick, M. G. Il-17a Modulators and Uses Thereof. WO2021055376A1, March 25, 2021. https://patents.google.com/patent/WO2021055376A1/en?oq=Dice+Alpha+Inc.+IL-17A+modulators+and+uses+thereof.+WO2021055376A1.+2021. (accessed 2024-05-14).

(181) Kim, J.-E.; Jung, K.; Kim, J.-A.; Kim, S.-H.; Park, H.-S.; Kim, Y.-S. Engineering of Anti-Human Interleukin-4 Receptor Alpha Antibodies with Potent Antagonistic Activity. Sci Rep 2019, 9 (1), 7772. 10.1038/s41598-019-44253-9.

(182) Obmolova, G.; Teplyakov, A.; Malia, T. J.; Keough, E.; Luo, J.; Sweet, R.; Jacobs, S. A.; Yi, F.; Hippensteel, R.; O’Neil, K. T.; Gilliland, G. L. Induced Conformational Change in Human IL D4 upon Binding of a SignalDneutralizing DARP In. Proteins 2015, 83 (6), 1191–1197. 10.1002/prot.24815.

(183) Vignali, D. A. A.; Kuchroo, V. K. IL-12 Family Cytokines: Immunological Playmakers. Nat Immunol 2012, 13 (8), 722–728. 10.1038/ni.2366.

(184) Teng, M. W. L.; Bowman, E. P.; McElwee, J. J.; Smyth, M. J.; Casanova, J.-L.; Cooper, A. M.; Cua, D. J. IL-12 and IL-23 Cytokines: From Discovery to Targeted Therapies for Immune-Mediated Inflammatory Diseases. Nat Med 2015, 21 (7), 719–729. 10.1038/nm.3895.

(185) Murray, P. J. The JAK-STAT Signaling Pathway: Input and Output Integration1. The Journal of Immunology 2007, 178 (5), 2623–2629. 10.4049/jimmunol.178.5.2623.

(186) Ecoeur, F.; Weiss, J.; Schleeger, S.; Guntermann, C. Lack of Evidence for Expression and Function of IL-39 in Human Immune Cells. PLoS One 2020, 15 (12), e0242329. 10.1371/journal.pone.0242329.

(187) Passaro, S.; Corso, G.; Wohlwend, J.; Reveiz, M.; Thaler, S.; Somnath, V. R.; Getz, N.; Portnoi, T.; Roy, J.; Stark, H.; Kwabi-Addo, D.; Beaini, D.; Jaakkola, T.; Barzilay, R. Boltz-2: Towards Accurate and Efficient Binding Affinity Prediction. bioRxiv June 18, 2025, p 2025.06.14.659707. 10.1101/2025.06.14.659707.

(188) Koštrun, S.; Fajdetić, A.; Pešić, D.; Brajša, K.; Bencetić Mihaljević, V.; Jelić, D.; Petrinić Grba, A.; Elenkov, I.; Rupčić, R.; Kapić, S.; Ozimec Landek, I.; Butković, K.; Grgičević, A.; Žiher, D.; Čikoš, A.; Padovan, J.; Saxty, G.; Dack, K.; Bladh, H.; Skak-Nielsen, T.; Feldbaek Nielsen, S.; Lambert, M.; Stahlhut, M. Macrolide Inspired Macrocycles as Modulators of the IL-17A/IL-17RA Interaction. J. Med. Chem. 2021, 64 (12), 8354–8383. 10.1021/acs.jmedchem.1c00327.

(189) Liu, S.; Dakin, L. A.; Xing, L.; Withka, J. M.; Sahasrabudhe, P. V.; Li, W.; Banker, M. E.; Balbo, P.; Shanker, S.; Chrunyk, B. A.; Guo, Z.; Chen, J. M.; Young, J. A.; Bai, G.; Starr, J. T.; Wright, S. W.; Bussenius, J.; Tan, S.; Gopalsamy, A.; Lefker, B. A.; Vincent, F.; Jones, L. H.; Xu, H.; Hoth, L. R.; Geoghegan, K. F.; Qiu, X.; Bunnage, M. E.; Thorarensen, A. Binding Site Elucidation and Structure Guided Design of Macrocyclic IL-17A Antagonists. Sci Rep 2016, 6 (1), 30859. 10.1038/srep30859.

(190) Ting, J. P.; Tung, F.; Antonysamy, S.; Wasserman, S.; Jones, S. B.; Zhang, F. F.; Espada, A.; Broughton, H.; Chalmers, M. J.; Woodman, M. E.; Bina, H. A.; Dodge, J. A.; Benach, J.; Zhang, A.; Groshong, C.; Manglicmot, D.; Russell, M.; Afshar, S. Utilization of Peptide Phage Display to Investigate Hotspots on IL-17A and What It Means for Drug Discovery. PLoS One 2018, 13 (1), e0190850. 10.1371/journal.pone.0190850.

(191) Espada, A.; Broughton, H.; Jones, S.; Chalmers, M. J.; Dodge, J. A. A Binding Site on IL-17A for Inhibitory Macrocycles Revealed by Hydrogen/Deuterium Exchange Mass Spectrometry. J Med Chem 2016, 59 (5), 2255–2260. 10.1021/acs.jmedchem.5b01693.

(192) Goedken, E. R.; Argiriadi, M. A.; Dietrich, J. D.; Petros, A. M.; Krishnan, N.; Panchal, S. C.; Qiu, W.; Wu, H.; Zhu, H.; Adams, A. M.; Bodelle, P. M.; Goguen, L.; Richardson, P. L.; Slivka, P. F.; Srikumaran, M.; Upadhyay, A. K.; Wu, B.; Judge, R. A.; Vasudevan, A.; Gopalakrishnan, S. M.; Cox, P. B.; Stoll, V. S.; Sun, C. Identification and Structure-Based Drug Design of Cell-Active Inhibitors of Interleukin 17A at a Novel C-Terminal Site. Sci Rep 2022, 12 (1), 14561. 10.1038/s41598-022-18760-1.

(193) Ramos, A. L.; Goedken, E. R.; Frank, K. E.; Argiriadi, M. A.; Bazzaz, S.; Bian, Z.; Brown, J. T. C.; Centrella, P. A.; Chen, H.-J.; Disch, J. S.; Donner, P. L.; Duignan, D. B.; Gikunju, D.; Greszler, S. N.; Guié, M.-A.; Habeshian, S.; Hartl, H. E.; Hein, C. D.; Hutchins, C. W.; Jetson, R.; Keefe, A. D.; Khan, H.; Li, H.-Q.; Olszewski, A.; Ortiz Cardona, B. J.; Osuma, A.; Panchal, S. C.; Phelan, R.; Qiu, W.; Shotwell, J. B.; Shrestha, A.; Srikumaran, M.; Su, Z.; Sun, C.; Upadhyay, A. K.; Wood, M. D.; Wu, H.; Zhang, R.; Zhang, Y.; Zhao, G.; Zhu, H.; Webster, M. P. Discovery of Small Molecule Interleukin 17A Inhibitors with Novel Binding Mode and Stoichiometry: Optimization of DNA-Encoded Chemical Library Hits to In Vivo Active Compounds. J. Med. Chem. 2024, 67 (8), 6456–6494. 10.1021/acs.jmedchem.3c02397.

(194) Kortemme, T.; Baker, D. A Simple Physical Model for Binding Energy Hot Spots in Protein–Protein Complexes. Proceedings of the National Academy of Sciences 2002, 99 (22), 14116–14121. 10.1073/pnas.202485799.

(195) Kortemme, T.; Kim, D. E.; Baker, D. Computational Alanine Scanning of Protein-Protein Interfaces. Sci STKE 2004, 2004 (219), pl2. 10.1126/stke.2192004pl2.

(196) Lipsh-Sokolik, R.; Fleishman, S. J. Addressing Epistasis in the Design of Protein Function. Proceedings of the National Academy of Sciences 2024, 121 (34), e2314999121. 10.1073/pnas.2314999121.

(197) Moy, F. J.; Diblasio, E.; Wilhelm, J.; Powers, R. Solution Structure of Human IL-13 and Implication for Receptor Binding1. Journal of Molecular Biology 2001, 310 (1), 219–230. 10.1006/jmbi.2001.4764.

(198) Eisenmesser, E. Z.; Horita, D. A.; Altieri, A. S.; Byrd, R. A. Solution Structure of Interleukin-13 and Insights into Receptor Engagement1. Journal of Molecular Biology 2001, 310 (1), 231–241. 10.1006/jmbi.2001.4765.

(199) Popovic, B.; Breed, J.; Rees, D. G.; Gardener, M. J.; Vinall, L. M. K.; Kemp, B.; Spooner, J.; Keen, J.; Minter, R.; Uddin, F.; Colice, G.; Wilkinson, T.; Vaughan, T.; May, R. D. Structural Characterisation Reveals Mechanism of IL-13-Neutralising Monoclonal Antibody Tralokinumab as Inhibition of Binding to IL-13Rα1 and IL-13Rα2. Journal of Molecular Biology 2017, 429 (2), 208–219. 10.1016/j.jmb.2016.12.005.

(200) Lecomte, F. C.; Joseph, J. S.; Stalewski, J.; Shen, Q.; Arnoult, E.; Sridhar, V.; Liu, M.; Hu, Y.; Gasendo, J. G.; Ben Arie, H.; Keinan, N.; Keidar, L.; Aviv, I.; Ruvinov, E.; Grandjean, J.; Dores-Silva, P. R.; Mak, A.; Santoso, B.; Kim, S.; Shende, V.; Wever, W. J.; Mirzadegan, T.; Zhu, Z.; Fuchs, B.; Pinton, P.; Szabady, R. Identification of an Induced Orthosteric Pocket in IL-23: A New Avenue for Non-Biological Therapeutic Targeting. ACS Chem. Biol. 2025, 20 (7), 1609–1618. 10.1021/acschembio.5c00181.

(201) Scott, O. B.; Edith Chan, A. W. ScaffoldGraph: An Open-Source Library for the Generation and Analysis of Molecular Scaffold Networks and Scaffold Trees. Bioinformatics 2020, 36 (12), 3930–3931. 10.1093/bioinformatics/btaa219.

(202) Tamarind Bio. https://www.tamarind.bio/ (accessed 2025-11-30).

