## Supplemental File for "Surveying the target tractability of cytokines with small molecules"

**Supplemental Information**

**Supplemental Figure 1. Representative scatter plots of prioritized cytokines IL-17, IL-13 and IL-23. (A)** IL-17 scatter plot. **(B)** IL-13 scatter plot. **(C)** IL-23 scatter plot.

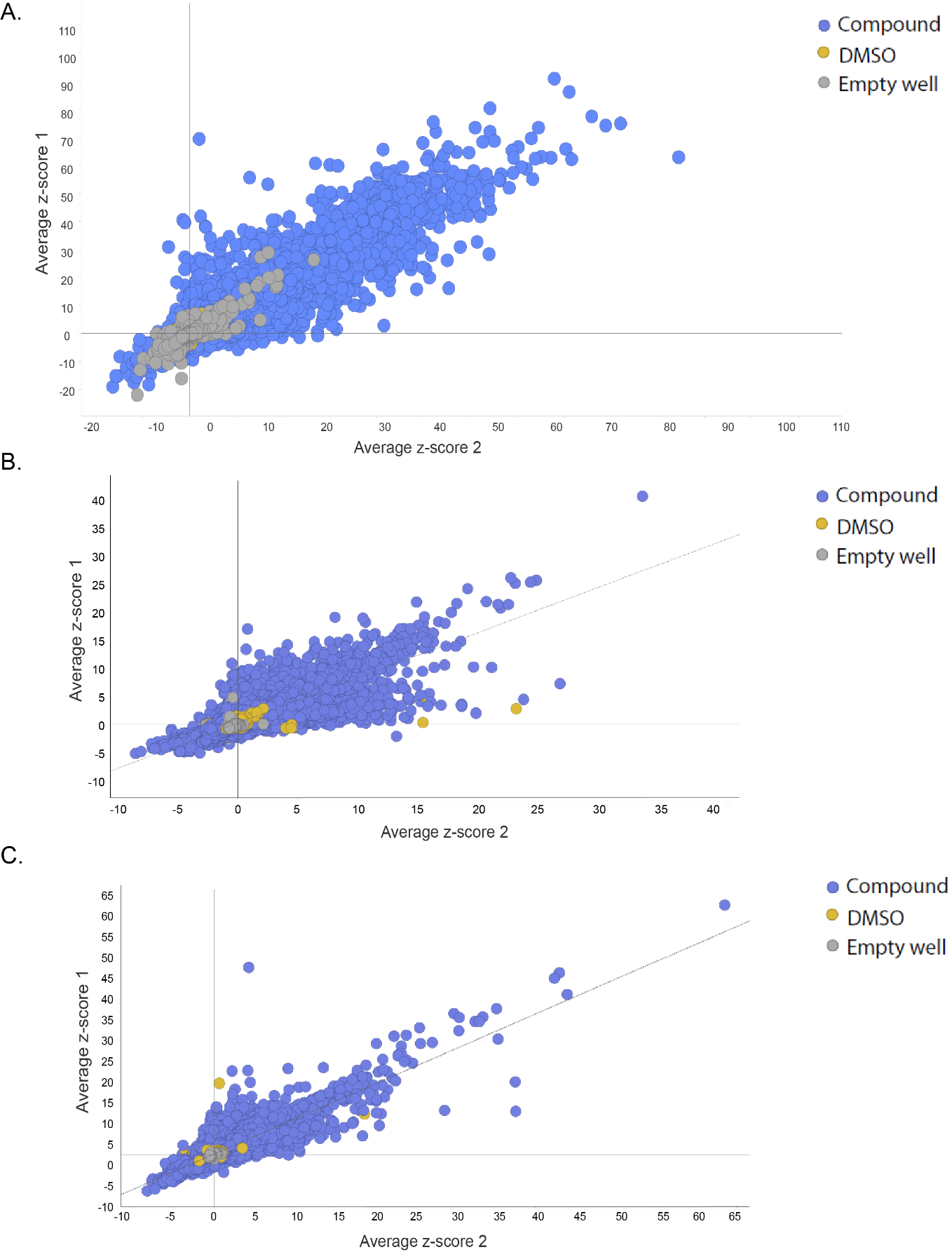

**Supplemental Figure 2. Comparison of quantitative estimate of drug-likeness (QED) between our ~65K compound collection and ~11K FDA-approved drugs from DrugBank.**

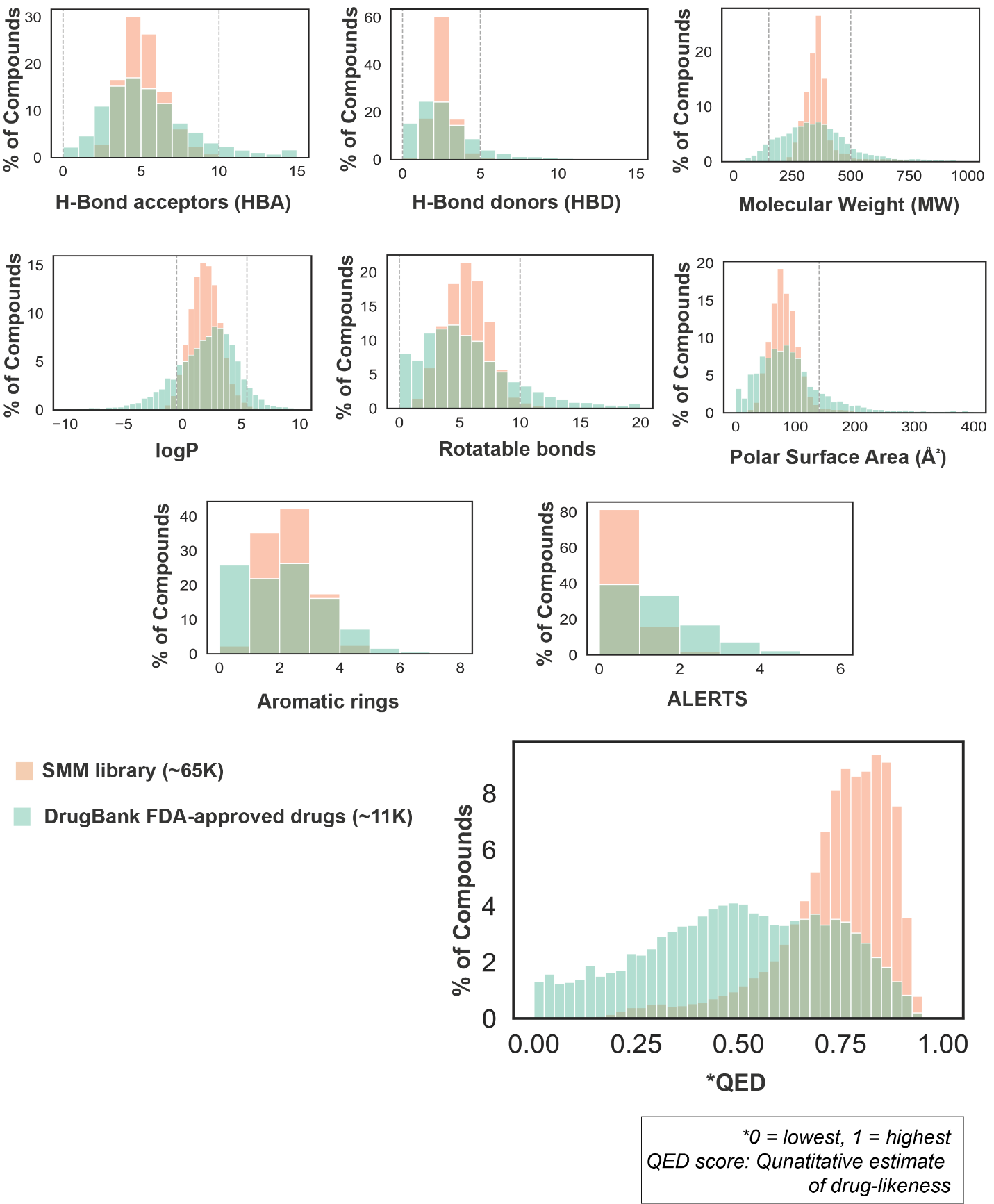

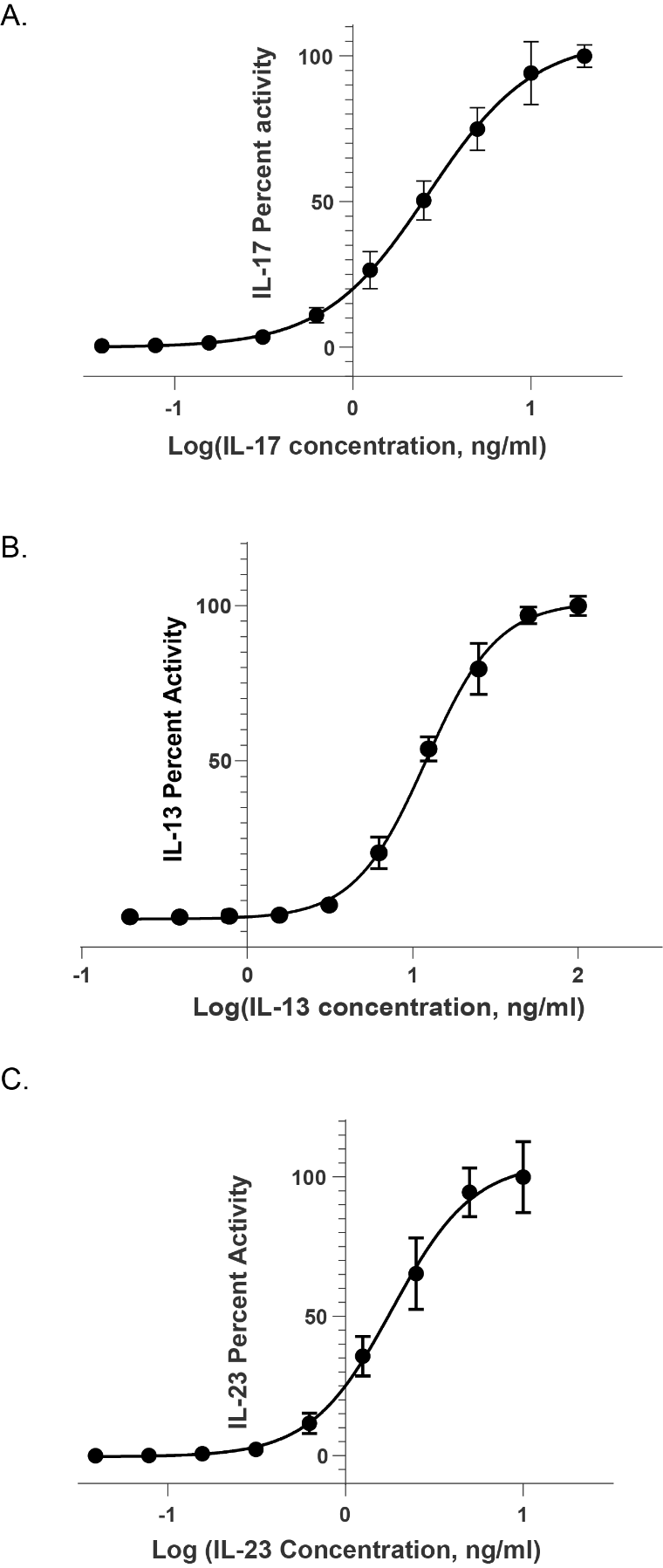
**Supplemental Figure 3. Activity of IL-17, IL-13 and IL-23 in their respective HEK-Blue cell line. (A)** IL-17 activity in HEK-Blue IL-17 cell line. **(B)** IL-13 activity in HEK-Blue IL-4/IL-13 cell line. **(C)** IL-23 activity in HEK-Blue IL-23 cell line.

**Supplemental Figure 4. Validation of Boltz-2 binding prediction by reproducing IL-17 interactions with Pfizer-designed macrocyclic IL-17 antagonist. (A)** Boltz-2 predicted the binding site of the Pfizer-designed macrocyclic IL-17 antagonist on IL-17 with a confidence score of 0.835 (ptm = 0.889; iptm = 0.891). The predicted IL-17 structure (green and grey) was aligned in PyMOL with the experimental IL-17–macrocycle complex (PDB ID: **5HI4**, shown in cyan, released in 2016). The Boltz-2-predicted macrocycle is shown in salmon, and the corresponding PDB-derived macrocycle in green. Key residues involved in ligand interaction are highlighted: **W67** (Chain A, blue) and **L97** (Chain B, orange). **(B)** Zoomed-in view of the binding pocket showing the alignment between the Boltz-2-predicted and experimentally observed macrocyclic conformations.

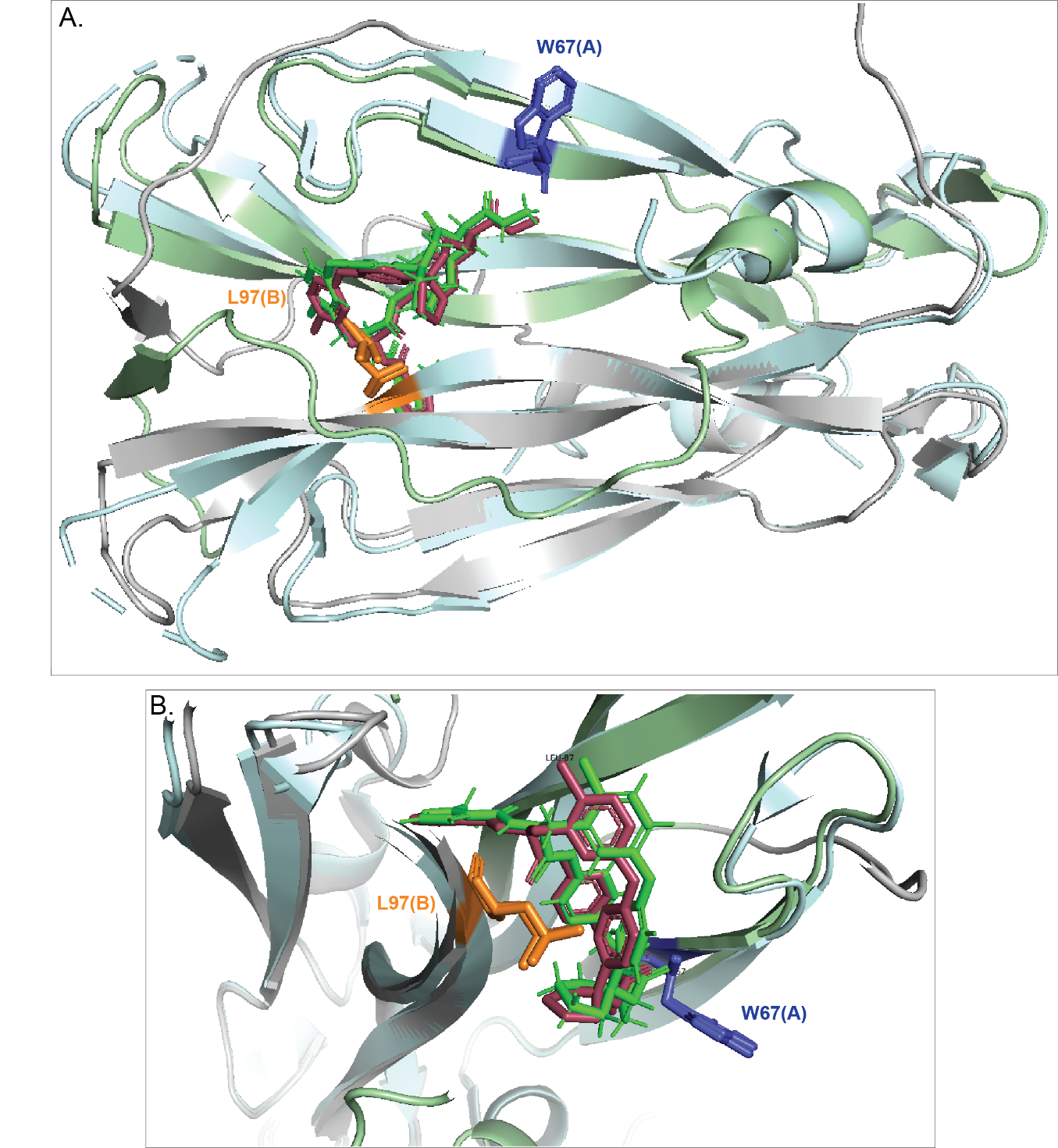

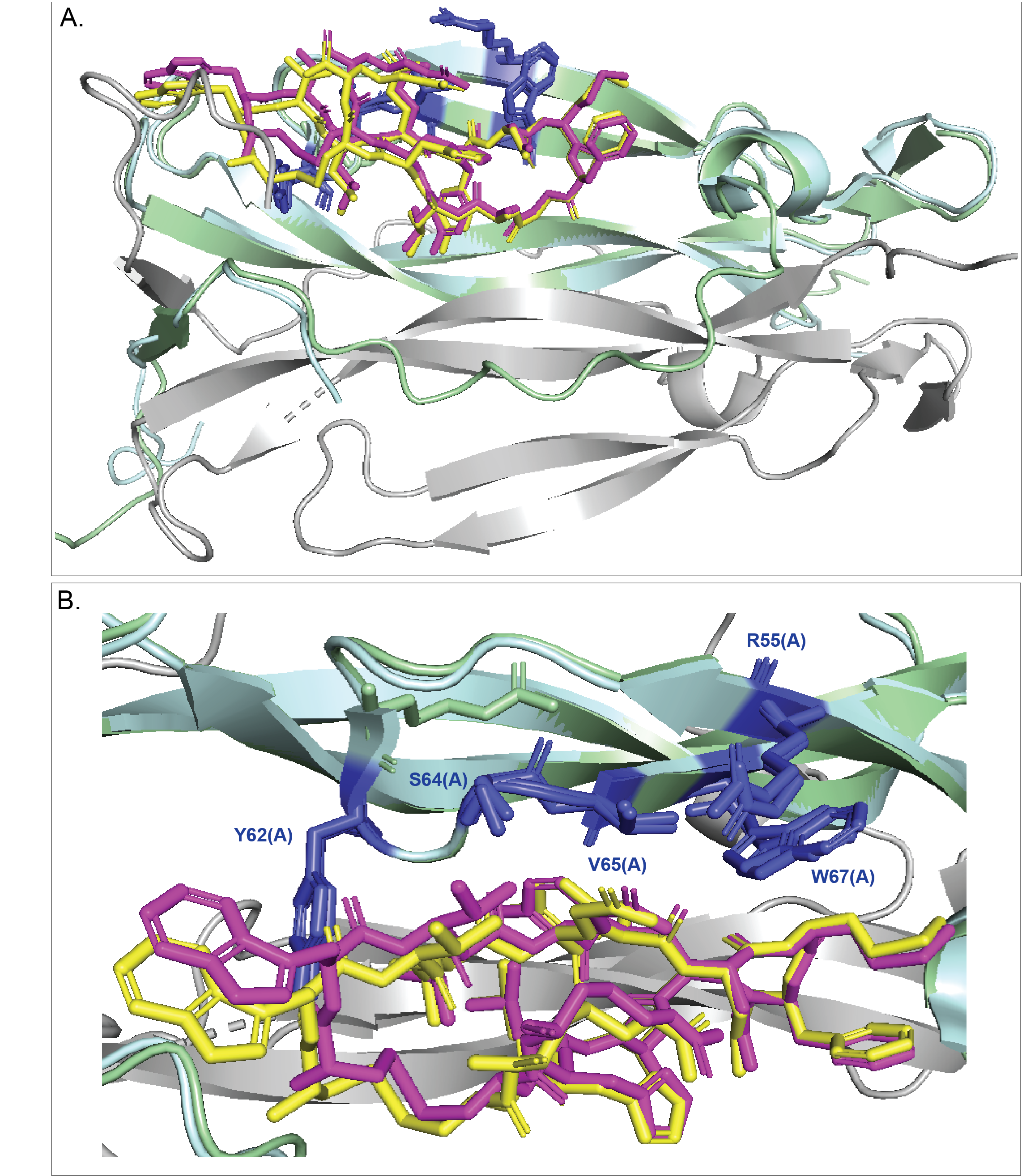
**Supplemental Figure 5. Validation of Boltz-2 binding prediction by reproducing IL-17 interactions with Eli Lilly-designed cyclic peptide inhibitor. (A)** Boltz-2 predicted the binding site of the Eli Lilly–designed macrocyclic IL-17 antagonist on IL-17 with a confidence score of 0.899 (ptm = 0.913; iptm = 0.913). The predicted IL-17 structure (green and grey) was aligned in PyMOL with the experimental IL-17–macrocycle complex (PDB ID: **5VB9**, shown in cyan, Chain A, released in 2018). The Boltz-2-predicted macrocycle is shown in dark pink, and the corresponding PDB-derived macrocycle in yellow. Key interacting residues are highlighted in blue (**Y62, S64, V65, R55, and W67**, Chain A). **(B)** Zoomed-in view of the binding pocket showing the overlap between the Boltz-2–predicted and experimentally observed macrocyclic conformations. The crystal structure of IL-17 (PDB ID: 5VB9) contains two macrocyclic ligands bound symmetrically to each IL-17 monomer; only Chain A with one bound macrocycle was used for alignment and comparison.

**Supplemental Figure 6. Boltz-2 binding prediction by reproducing IL-17 interactions with Pfizer-designed linear peptide inhibitor.** Boltz-2 predicted the binding site of the Pfizer–designed linear peptide IL-17 inhibitor on IL-17 with a confidence score of 0.855 (ptm = 0.907; iptm = 0.898). The predicted IL-17 structure (green and grey) was aligned in PyMOL with the experimental IL-17–peptide complex (PDB ID: **5HHX**, shown in cyan, Chain A, released in 2016). The Boltz-2-predicted inhibitor is shown in dark pink, and the corresponding PDB-derived inhibitor in blue. Key interacting residues are highlighted in light orange (**N108, F110, L112, and K114**, Chain A). The crystal structure of IL-17 (PDB ID: 5HHX) contains missing residues in Chain B; therefore, only Chain A bound to the peptide inhibitor was used for alignment.

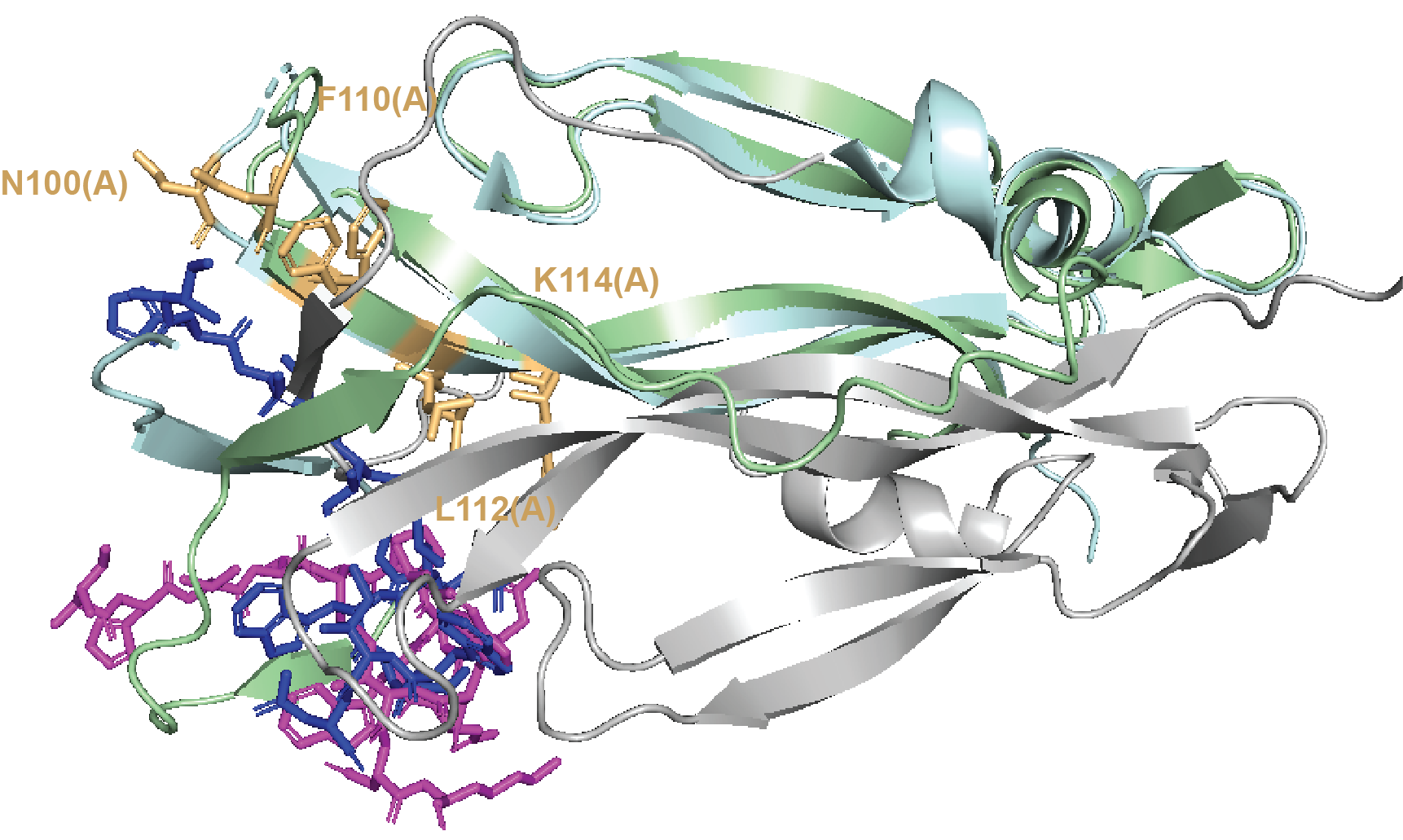

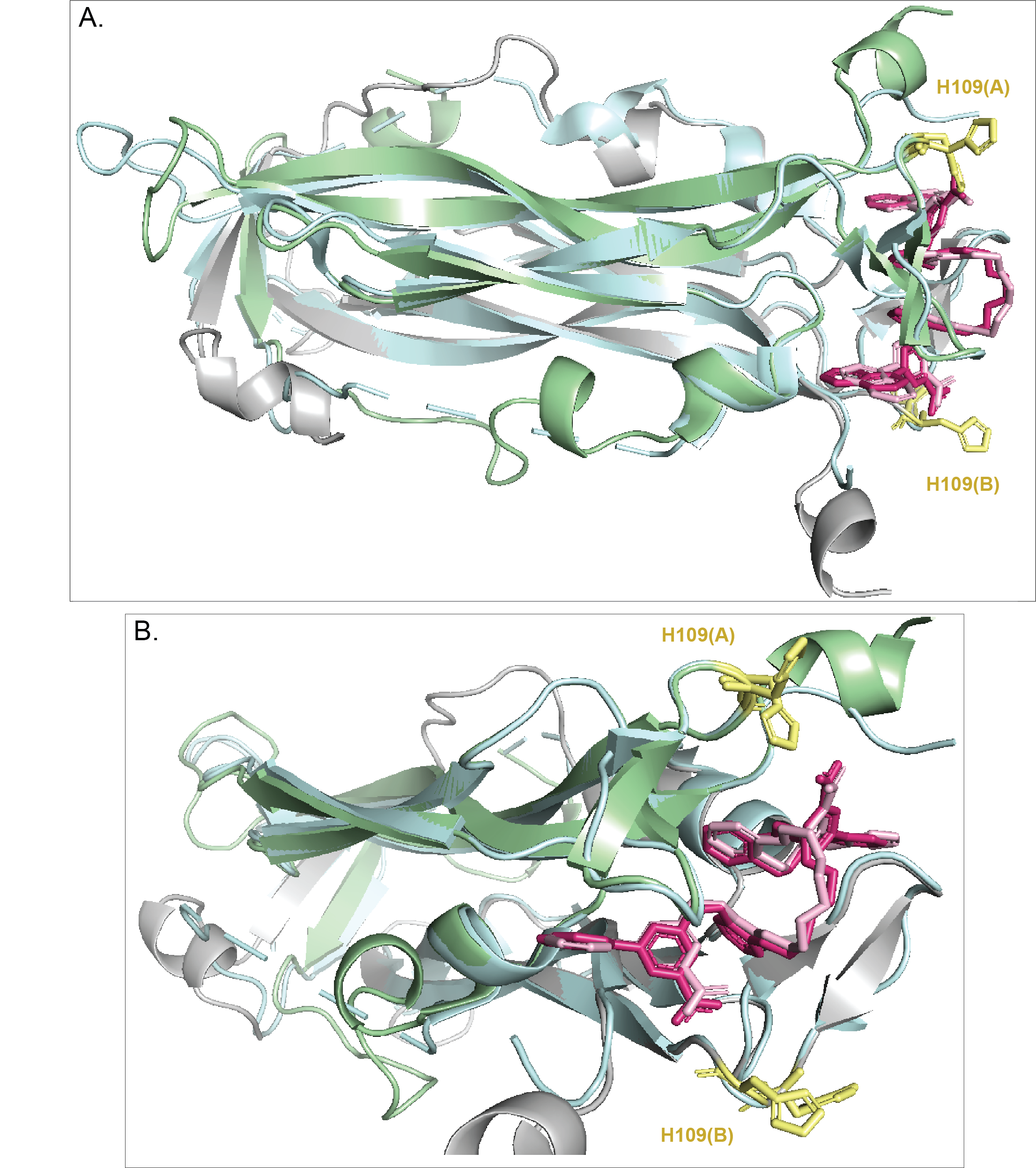
**Supplemental Figure 7. Validation of Boltz-2 binding prediction by reproducing IL-17 interactions with AbbVie-designed compound that binds to a novel C-terminus.** Boltz-2 predicted the binding site of the AbbVie–designed IL-17 inhibitor on IL-17 with a confidence score of 0.824 (ptm = 0.854; iptm = 0.843). The predicted IL-17 structure (green and grey) was aligned in PyMOL with the experimental IL-17–inhibitor complex (PDB ID: **8DYF**, shown in cyan, released in 2022). The Boltz-2-predicted inhibitor is shown in dark pink, and the corresponding PDB-derived inhibitor in light pink. Key interacting residues are highlighted in pale yellow (**H109**, Chain A and **H109**, Chain B). **(B)** Zoomed-in view of the binding pocket showing the overlap between the Boltz-2–predicted and experimentally observed AbbVie’s IL-17 antagonist conformations.

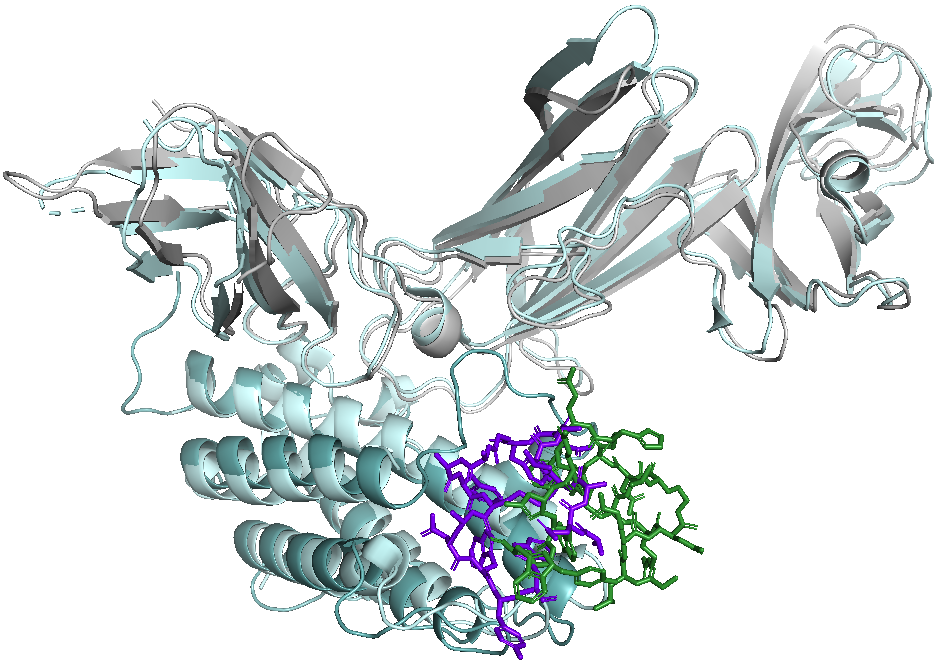
**Supplemental Figure 8. Validation of Boltz-2 binding prediction by reproducing IL-23 interactions with inhibitory peptide 23-446 at an induced orthosteric binding pocket on IL-23p19 subunit.** Boltz-2 predicted the binding site of the 23-446 peptide on IL-23p19 subunit with a confidence score of 0.882 (ptm = 0.918; iptm = 0.891). The predicted IL-23 structure (deep teal and grey) was aligned in PyMOL with the experimental IL-23–peptide complex (PDB ID: **8UUI**, shown in cyan, released in 2025). The Boltz-2-predicted inhibitor is shown in green, and the corresponding PDB-derived inhibitor in purple.

**Supplemental Figure 9. Boltz-2 prediction of IL-17 inhibitor 5 binding and *in silico* mutagenesis of the C-terminal pocket**. **(A)** Inhibitor **5** (purple) binds at the IL-17 C-terminal pocket, positioned between residues M110(A/B) (magenta) and H109(A/B) (yellow). **(B)** Close-up view showing inhibitor **5** nestled within the hydrophobic cleft formed by M110(A) and M110(B). **(C–D)** *In silico* alanine-scanning mutagenesis using Boltz-2 highlights the functional importance of M110 in stabilizing the inhibitor pose. Substitution of M110(A) or M110(B) with alanine (red) collapses the hydrophobic pocket and destabilizes the ligand orientation. The black conformation represents the displaced pose adopted by inhibitor **5** upon mutation, illustrating the loss of M110-mediated anchoring at the IL-17 C-terminus.

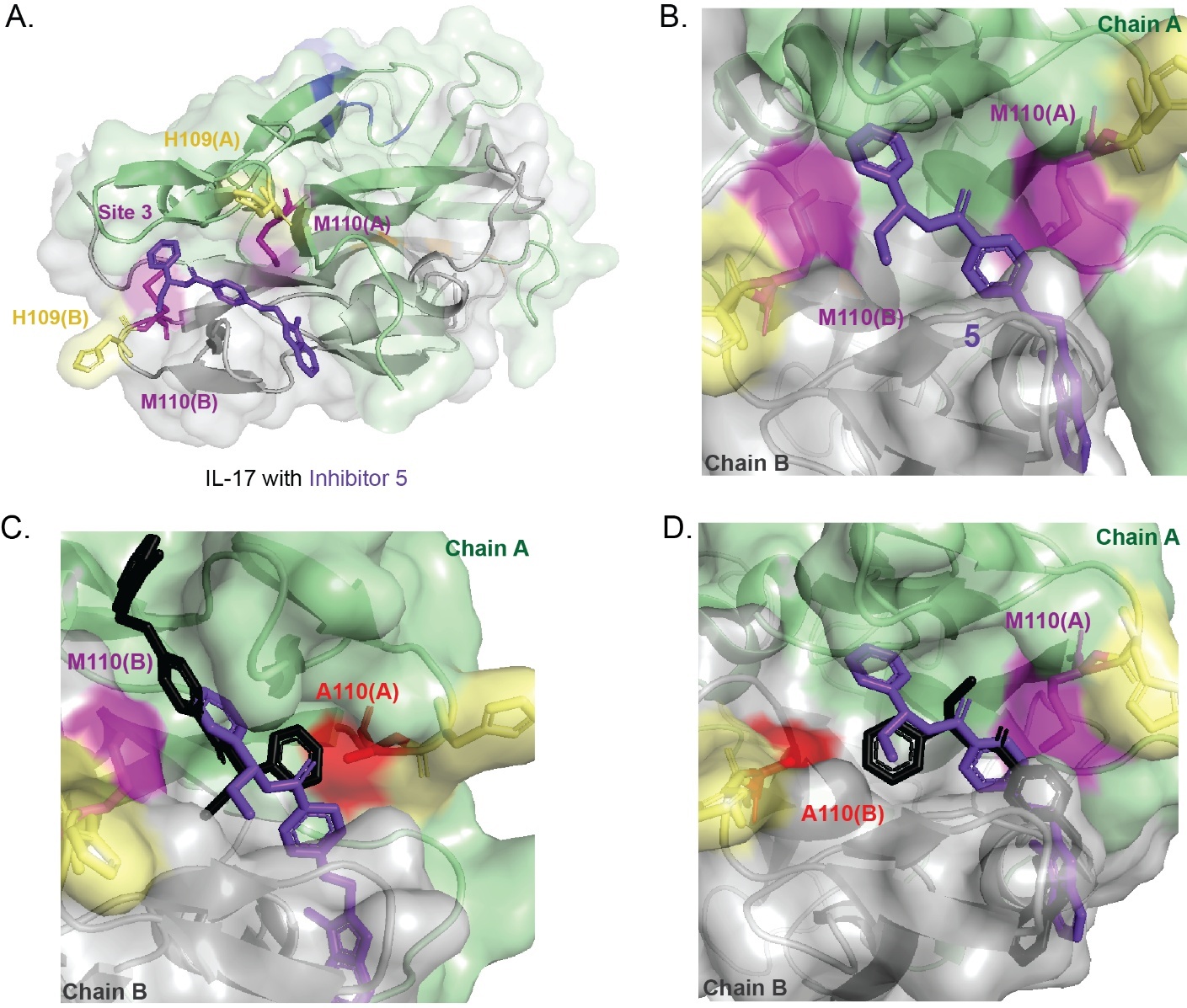

**Supplemental Figure 10. Boltz-2 prediction of IL-17 inhibitor 22 binding and in silico mutagenesis of the site-2 interface. (A)** Inhibitor **22** (purple) engages the site-2 interface of IL-17, positioned between W67(A) (blue) and L97(B) (orange). **(B)** Enlarged view showing inhibitor **22** situated between W67(A) and L97(B), spanning the cleft formed at the A–B chain interface. **(C–D)** *In silico* alanine-scanning mutagenesis using Boltz-2 shows that replacing W67(A) or L97(B) with alanine (red) alters the pocket geometry and affects the predicted ligand pose. The black conformation represents the shifted orientation adopted by inhibitor **22** upon mutation, indicating reduced compatibility of the ligand with the remodeled interface.

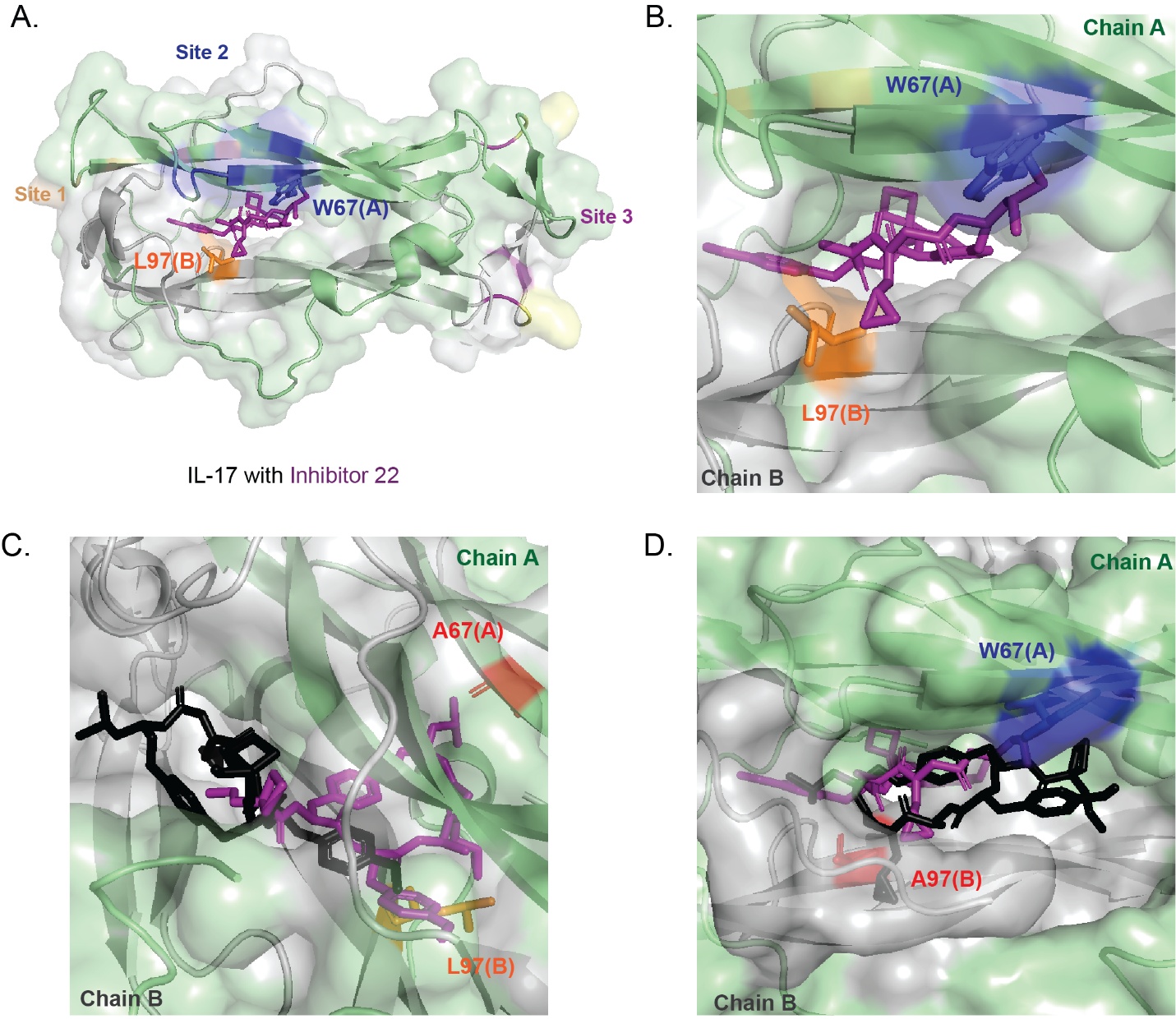

**Supplemental Figure 11. Boltz-2 prediction of IL-17 inhibitor 24 binding and in silico mutagenesis of the C-terminal pocket**. **(A)** Inhibitor **24** (cyan) occupies the C-terminal pocket of IL-17, positioned between M110(A/B) (magenta) and H109(A/B) (yellow), similar to other chemotypes engaging this region. **(B)** Expanded view showing inhibitor **24** situated within the hydrophobic cleft formed by M110(A) and M110(B). **(C–D)** *In silico* alanine-scanning mutagenesis using Boltz-2 indicates that replacing M110(A) or M110(B) with alanine (red) alters the pocket contours and affects the predicted pose of inhibitor **24**. The black conformation reflects the shifted orientation adopted by the ligand after mutation, consistent with reduced compatibility with the remodeled C-terminal site.

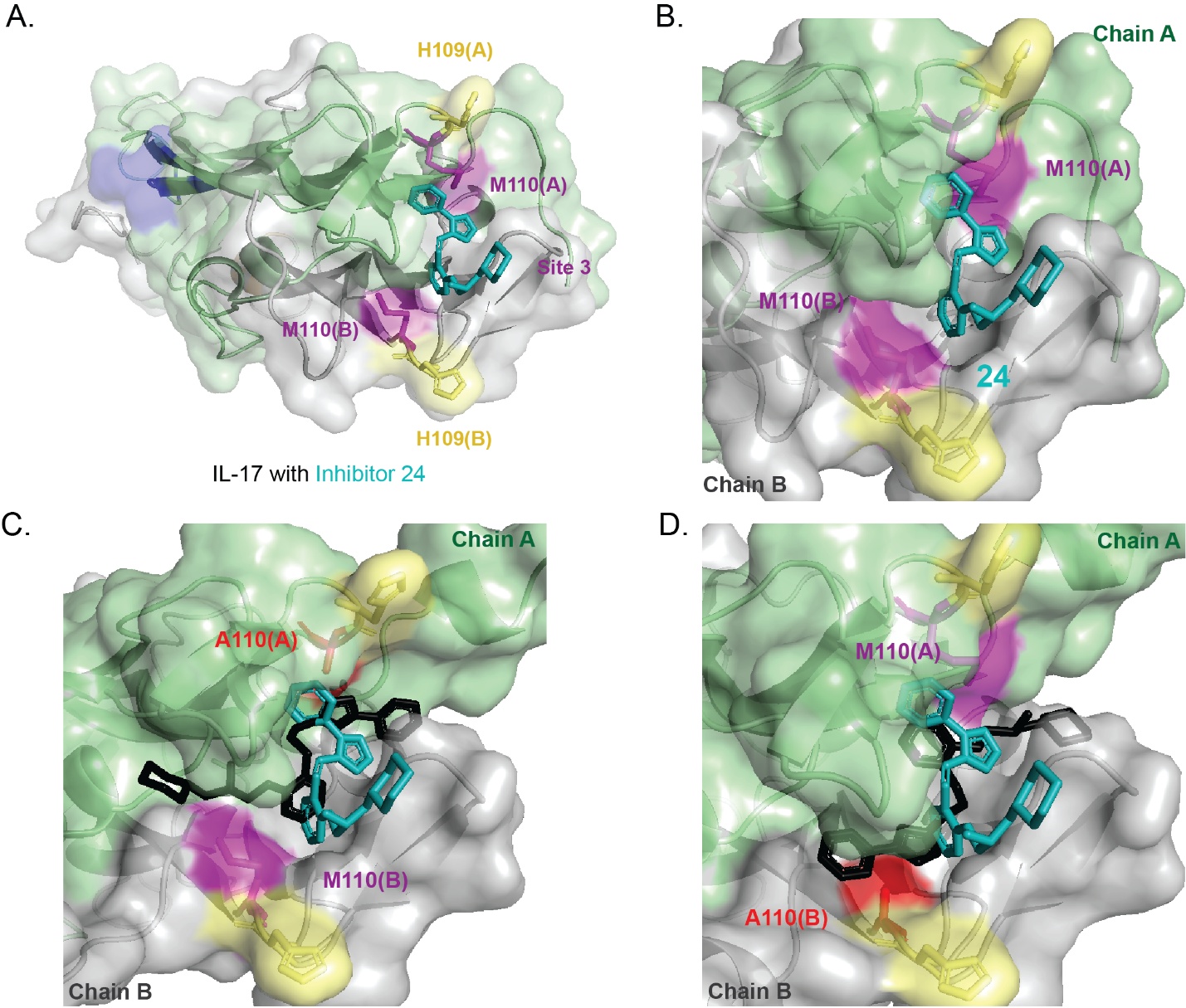

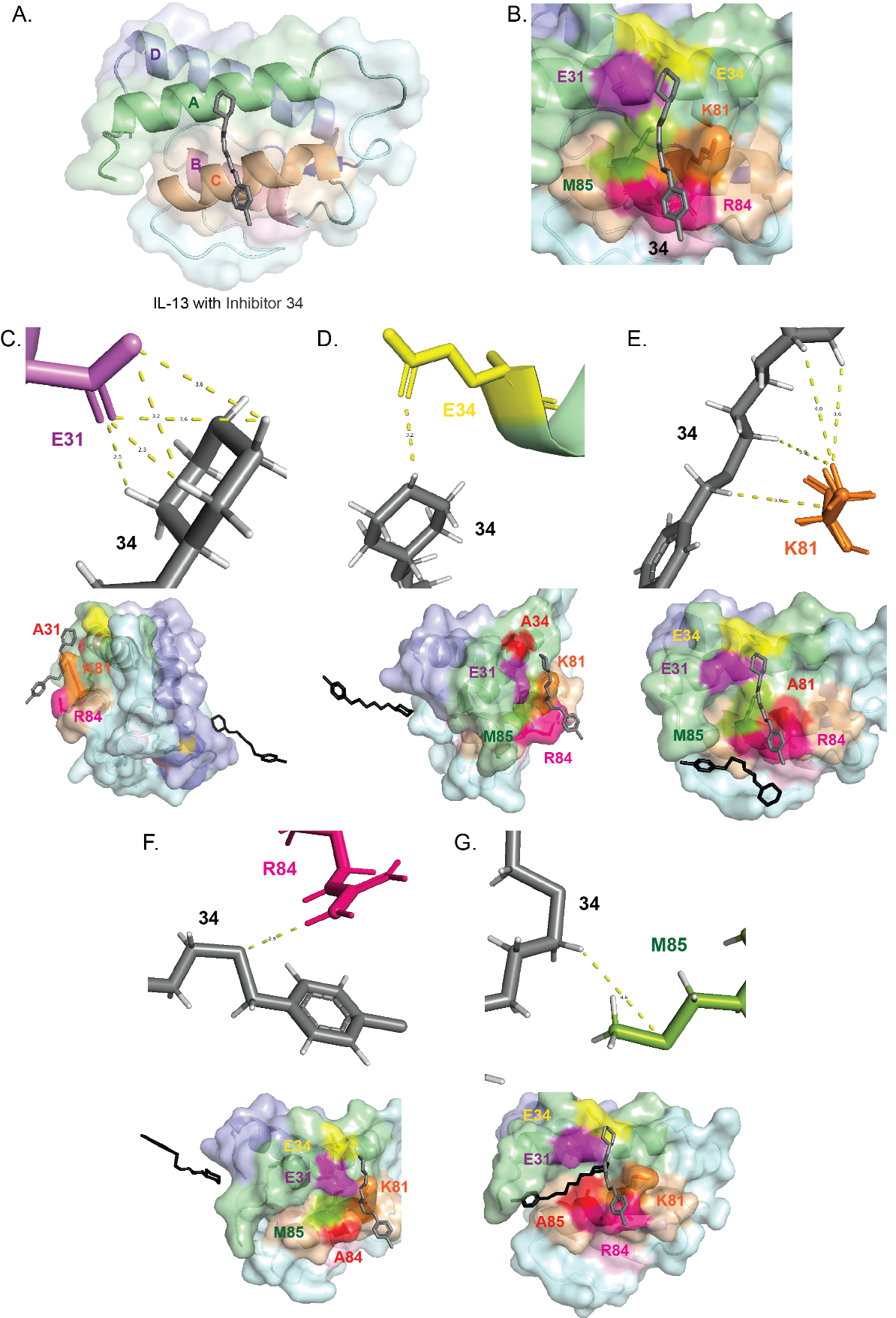
**Supplemental Figure 12**. **Boltz-2 prediction of IL-13 inhibitor 34 binding and *in silico* mutagenesis of the helix-A/C interface.** **(A)** Inhibitor **34** (dark grey) engages spans over IL-13’s helix A and C. **(B)** Enlarged view showing inhibitor **34** positioned among E31, E34, K81, R84, and M85, which create a mixed polar–hydrophobic environment around the ligand.
**(C–G)** *In silico* alanine-scanning mutagenesis using Boltz-2 was performed to assess residue contributions to this pocket. Substitutions of E31, M85, K81, R84, or E34 with alanine (red) alters local contacts and changes the predicted arrangement of inhibitor **34**. The black conformation represents the ligand pose adopted after each mutation, consistent with reduced compatibility between the modified pocket and the bound chemotype.

**Supplemental Table 1. 32 Human Cytokines screened in Small-molecule microarray.**

| **No.** | **Cytokines** | **Implication in immune-mediated disorders** |
| --- | --- | --- |
| **1.** | **IL-1α** | Rheumatoid arthritis, Type 1 Diabetes Mellitus, Cytokine release syndrome.^1–9^ |
| **2.** | **IL-1β** | Systemic juvenile idiopathic arthritis, Type 1 Diabetes Mellitus, Multiple sclerosis Cytokine release syndrome.^5,8–13^ |
| **3.** | **IL-2** | Asthma, Plaque psoriasis.^14–17^ |
| **4.** | **IL-5** | Severe asthma, Eosinophilic granulomatosis with polyangiitis.^18–22^ |
| **5.** | **IL-6** | Systemic juvenile idiopathic arthritis, Rheumatoid arthritis, Giant cell arthritis, COVID-19, Neuromyelitis optica spectrum disorders, Multicentric Castleman’s disease, Type 1 Diabetes Mellitus, Systemic lupus erythematosus, Cytokine release syndrome.^23–34^ |
| **6.** | **IL-7** | Type 1 Diabetes Mellitus, T-cell acute lymphoblastic leukemia, Multiple sclerosis, Rheumatoid arthritis.^35–40^ |
| **7.** | **IL-8** | Plaque psoriasis, Breast cancer, Melanoma, Prostate cancer, Lung cancer.^17,41–46^ |
| **8.** | **IL-10** | Systemic lupus erythematosus.^47–49^ |
| **9.** | **IL-11** | Asthma, Multiple sclerosis, Fibrosis, Inflammatory arthritis.^50–54^ |
| **10.** | **IL-12** | Moderate to severe ulcerative colitis and Crohn’s disease, Rheumatoid arthritis, Type 1 Diabetes Mellitus, Plaque psoriasis, Inflammatory bowel disease, Multiple sclerosis.^55–61^ |
| **11.** | **IL-13** | Moderate to severe atopic dermatitis, Asthma, Cancer.^62–69^ |
| **12.** | **IL-17A** | Moderate to severe plaque psoriasis, Rheumatoid arthritis, Type 1 Diabetes Mellitus, Multiple sclerosis, Cytokine release syndrome.^70–79^ |
| **13.** | **IL-18** | Rheumatoid arthritis, Angiogenesis.^80–82^ |
| **14.** | **IL-21** | Rheumatoid arthritis, Type 1 Diabetes Mellitus.^83–86^ |
| **15.** | **IL-22** | Rheumatoid arthritis, Plaque psoriasis.^87–90^ |
| **16.** | **IL-23** | Moderate to severe ulcerative colitis and Crohn’s disease, Rheumatoid arthritis, Type 1 Diabetes Mellitus, Plaque psoriasis, Multiple sclerosis.^55,56,91–94^ |
| **17.** | **IL-33** | Asthma, Rheumatoid arthritis.^95–101^ |
| **18.** | **CD40L** | Rheumatoid arthritis, Systemic lupus erythematosus, Multiple sclerosis, Inflammatory bowel disease.^102–107^ |
| **19.** | **C5** | Paroxysmal nocturnal hemoglobinuria, Systemic lupus erythematosus.^108–113^ |
| **20.** | **GM-CSF** | Rheumatoid arthritis , Multiple sclerosis.^114–122^ |
| **21.** | **G-CSF** | Rheumatoid arthritis, Cytokine release syndrome.^123–127^ |
| **22.** | **MIF** | Multiple sclerosis, Systemic lupus erythematosus, Sepsis, Acute Respiratory distress.^128–130^ |
| **23.** | **IFNα1** | Moderate to severe Systemic lupus erythematosus, Sjögren's syndrome, Type 1 Diabetes Mellitus, Rheumatoid arthritis.^131–134^ |
| **24.** | **TGF-β** | Systemic lupus erythematosus, Multiple Sclerosis, solid tumors and Asthma.^135–139^ |
| **25.** | **TNF-α** | Rheumatoid arthritis, Crohn’s disease, Ankylosing spondylitis, Psoriatic arthritis, Psoriasis, Idiopathic arthritis, Ulcerative colitis and non-infectious uveitis, Type 1 Diabetes Mellitus, Breast cancer, Cytokine release syndrome.^139–157^ |
| **26.** | **CCL1** | Cancer, Pulmonary fibrosis, Type-2 mediated allergic diseases, Inflammatory bowel disease.^158–162^ |
| **27.** | **CCL2** | Breast cancer, Rheumatoid arthritis, Multiple sclerosis, Inflammation, Fibrosis.^163,164^ |
| **28.** | **CCL3** | Breast cancer, Rheumatoid arthritis.^165–169^ |
| **29.** | **CCL5** | Breast cancer, Colorectal cancer, Rheumatoid arthritis, Allergic reactions, Inflammatory bowel disease.^166,170–174^ |
| **30.** | **CXCL1** | Lung cancer, Gastric cancer, Colorectal cancer, Breast cancer, Pancreatic cancer, Liver cancer, Ovarian cancer, Rheumatoid arthritis, Respiratory disorders, Systemic lupus erythematosus.^175–179^ |
| **31.** | **CXCL10** | Rheumatoid arthritis, Ulcerative colitis, Solid tumors.^180–184^ |
| **32.** | **CXCL12** | Autoimmune arthritis, Asthma, Psoriasis, Inflammatory bowel disease, Multiple sclerosis, Solid tumors^185–190^ |

**Supplemental Table 2. Boltz-2 calculations for screening data for prioritized cytokines.**

| **Cytokine** | **Compound type** | **Confidence score** | **ptm** | **iptm** |
| --- | --- | --- | --- | --- |
| **IL-17** | SMM-negative **1** | 0.459 | 0.767 | 0.733 |
|  | Binder **2** | 0.824 | 0.927 | 0.923 |
|  | Inhibitor **5** | 0.836 | 0.925 | 0.921 |
|  | Inhibitor **9** | 0.799 | 0.900 | 0.891 |
|  | Inhibitor **22** | 0.825 | 0.891 | 0.883 |
|  | Inhibitor **24** | 0.821 | 0.900 | 0.893 |
| **IL-13** | SMM-negative **26** | 0.578 | 0.975 | 0.964 |
|  | Binder **27** | 0.816 | 0.835 | 0.905 |
|  | Inhibitor **28** | 0.819 | 0.850 | 0.911 |
|  | Inhibitor **34** | 0.729 | 0.870 | 0.705 |
| **IL-23** | SMM-negative **35** | 0.671 | 0.886 | 0.841 |
|  | Binder **36** | 0.882 | 0.915 | 0.895 |
|  | Inhibitor **39** | 0.882 | 0.924 | 0.908 |

**Supplemental Table 3. Preliminary screening of validated IL-17 binders at 10 µM in HEK-Blue IL-17 reporter cell line.**

| **Compound ID** | **Mean percent inhibition at 10 µM** | **Standard deviation** |
| --- | --- | --- |
| 2 | 9.2 | 16.7 |
| 3 | 42.9 | 14.4 |
| 4 | 27.7 | 5.5 |
| 5 | 45.6 | 14.7 |
| 6 | 18.9 | 10.7 |
| 7 | 30 | 10.6 |
| 8 | 19.7 | 10.1 |
| 9 | 73.9 | 7.4 |
| 10 | 37.4 | 10.2 |
| 11 | 35.2 | 8.3 |
| 12 | 74.7 | 5.8 |
| 13 | 32.3 | 12.2 |
| 14 | 29.9 | 9.2 |
| 15 | 42.8 | 8.7 |
| 16 | 23.3 | 10.3 |
| 17 | 25.6 | 15.7 |
| 18 | 17.6 | 19.3 |
| 19 | 21.1 | 22.4 |
| 20 | 23.9 | 8.8 |
| 21 | 51.4 | 4.4 |
| 22 | 50.9 | 12.9 |
| 23 | 26.7 | 12.7 |
| 24 | 54.4 | 18 |
| 25 | 17.6 | 24.8 |

**Supplemental Table 4. Preliminary screening of validated IL-13 binders at 10 µM in HEK-Blue IL-4/IL-13 reporter cell line.**

| **Compound ID** | **Mean percent inhibition at 10 µM** | **Standard deviation** |
| --- | --- | --- |
| 27 | 41.7 | 7.4 |
| 28 | 51.5 | 4.7 |
| 29 | 47.9 | 8.6 |
| 30 | 60.6 | 4.8 |
| 31 | 41.3 | 2.8 |
| 32 | 47.5 | 10.4 |
| 33 | 21.1 | 5.3 |
| 34 | 63.6 | 14.2 |

**Supplemental Table 5. Preliminary screening of validated IL-23 binders at 10 µM in HEK-Blue IL-23 reporter cell line.**

| **Compound ID** | **Mean percent inhibition at 10 µM** | **Standard deviation** |
| --- | --- | --- |
| 36 | 16.4 | 3.6 |
| 37 | 37.9 | 9.4 |
| 38 | 34.6 | 6.8 |
| 39 | 89.5 | 3.3 |

**Supplemental Figure 9. Additional IL-17 binders exhibiting ≥20% inhibition at 10 µM in preliminary screening using the HEK-Blue IL-17 reporter cell line.**

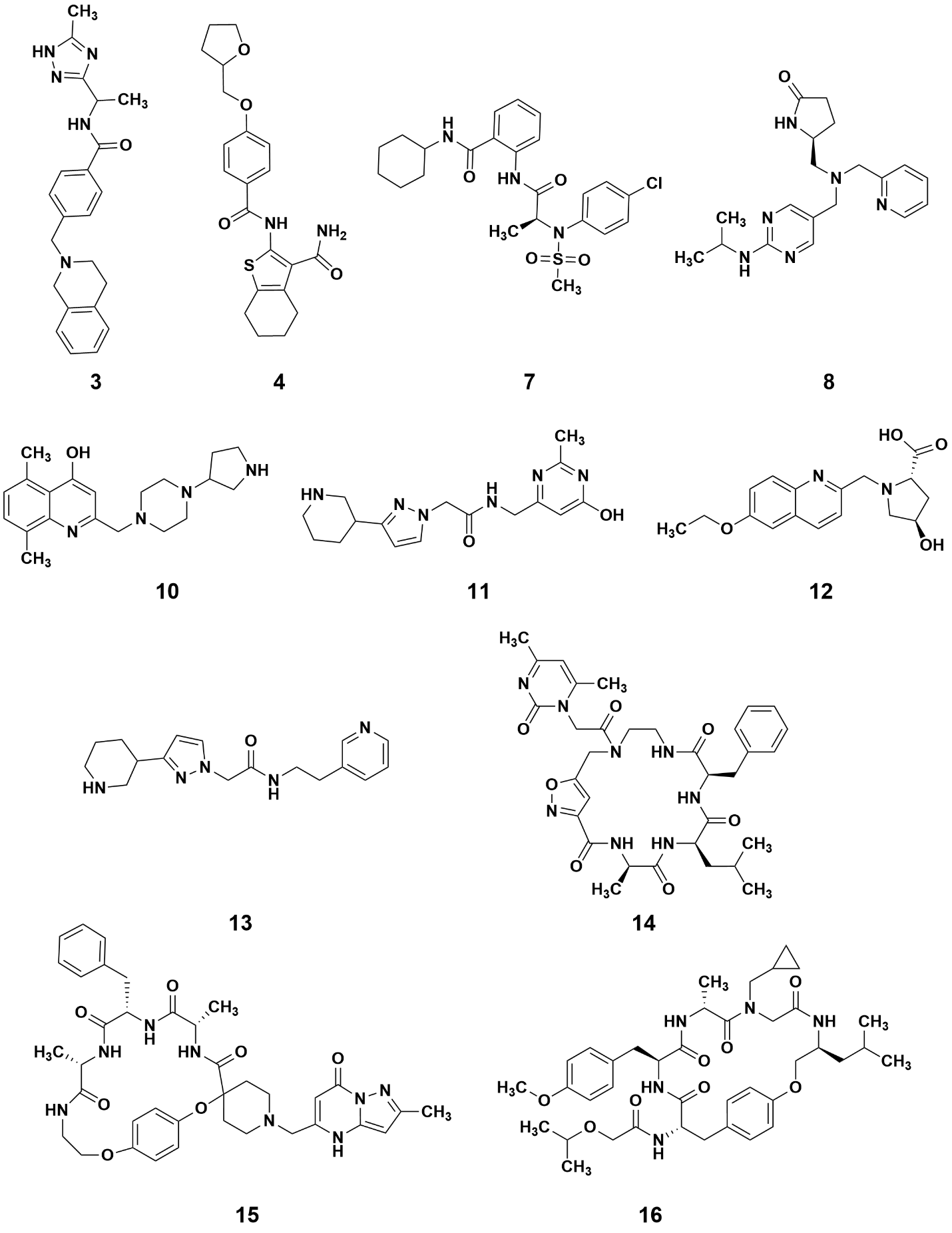

**Supplemental Figure 10. Additional IL-17 binders exhibiting ≥20% inhibition at 10 µM in preliminary screening using the HEK-Blue IL-17 reporter cell line.**

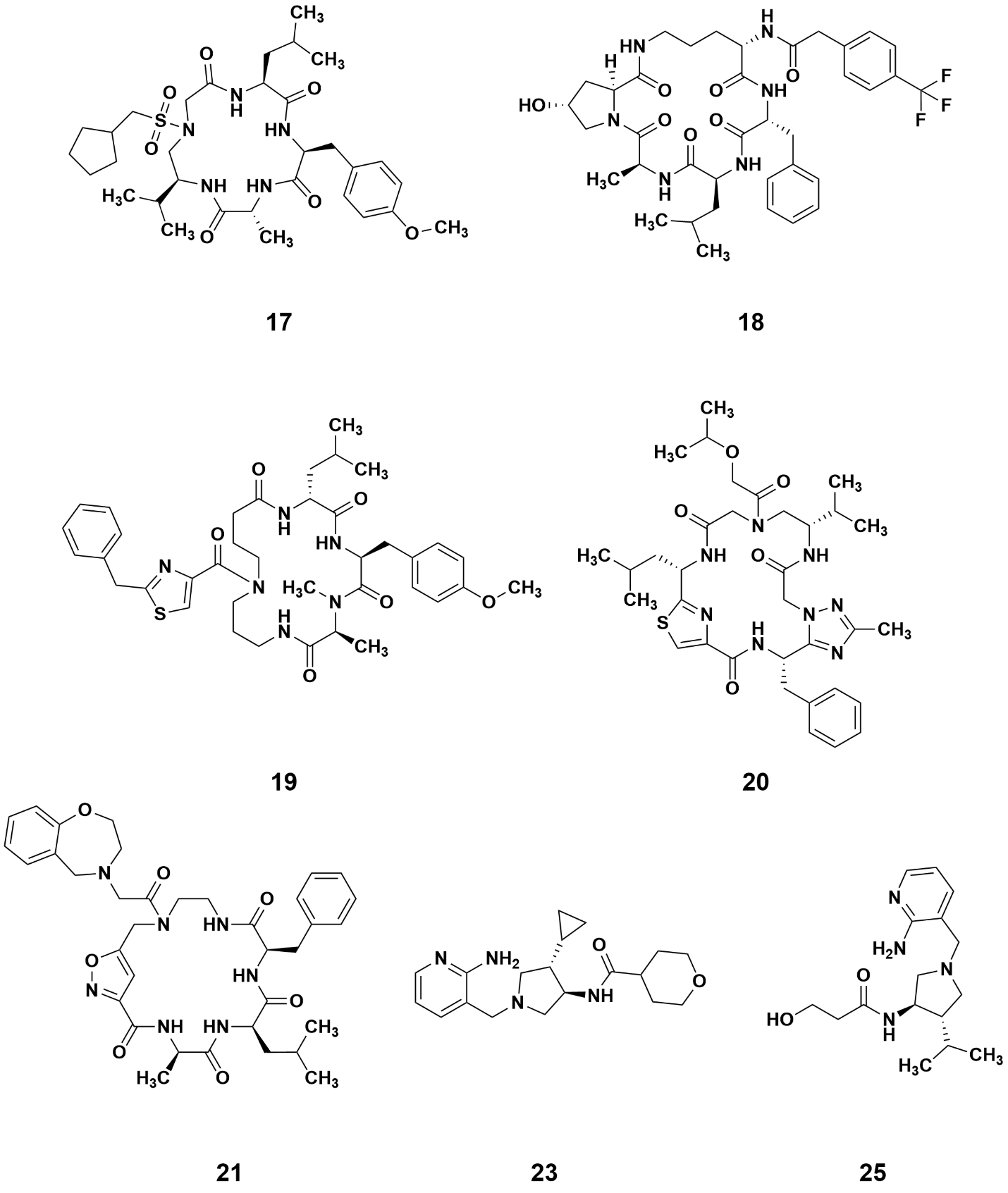

**Supplemental Figure 11. Additional IL-13 binders exhibiting ≥20% inhibition at 10 µM in preliminary screening using the HEK-Blue IL-4/IL-13 reporter cell line.**

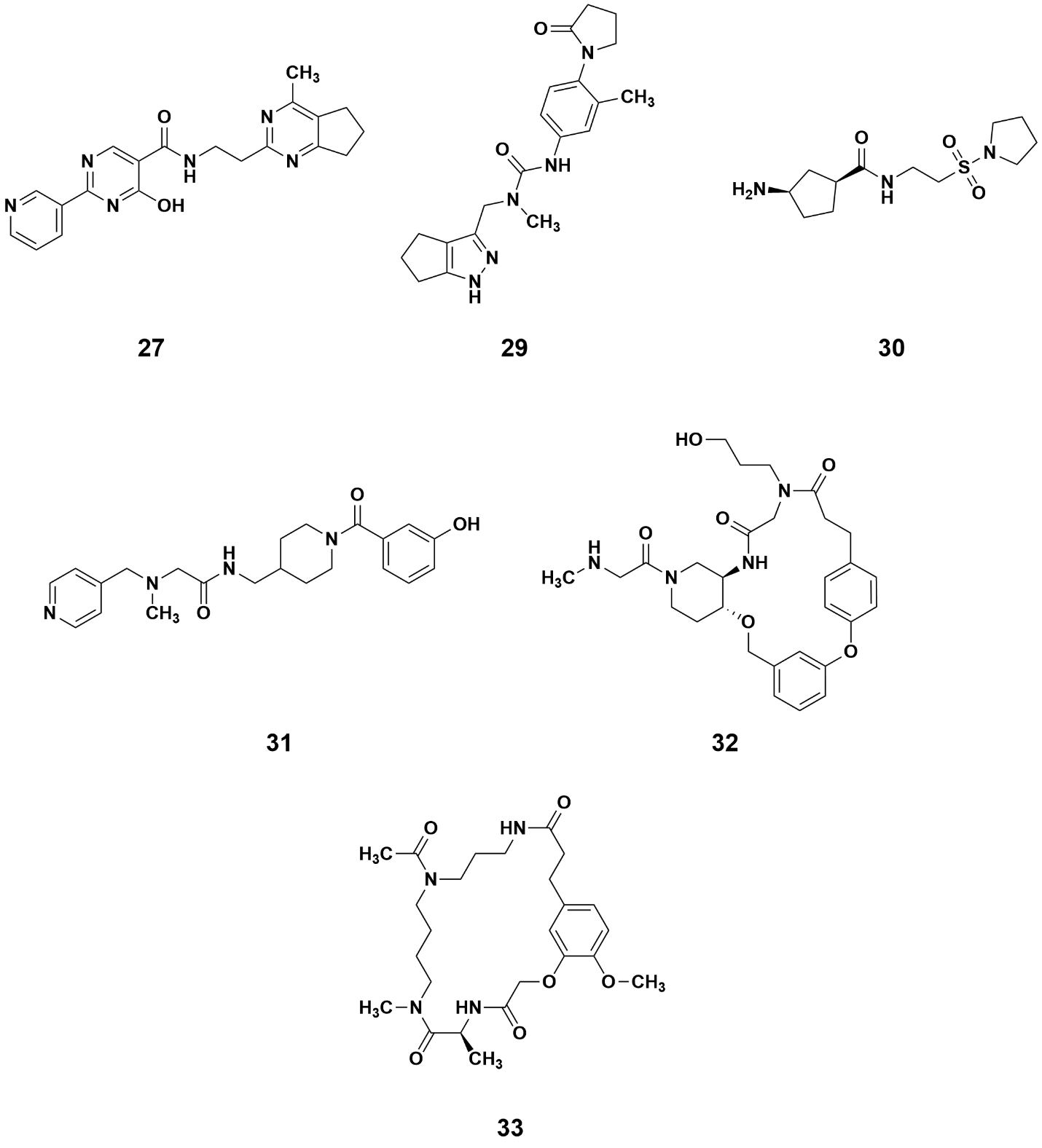

**Supplemental Figure 12. Additional IL-23 binders exhibiting ≥20% inhibition at 10 µM in preliminary screening using the HEK-Blue IL-23 reporter cell line.**

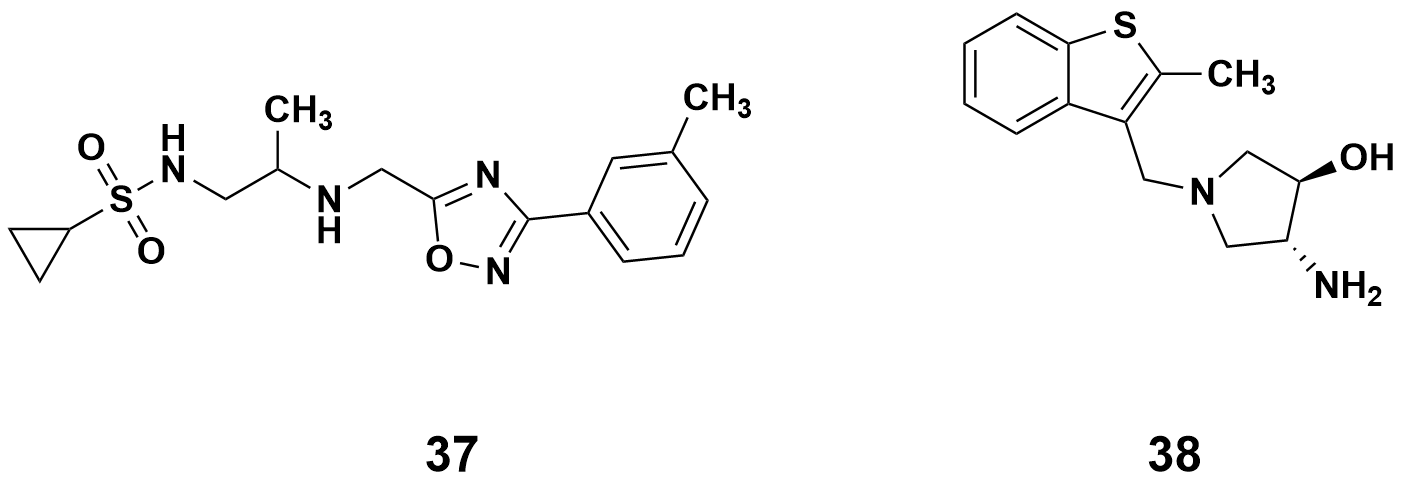

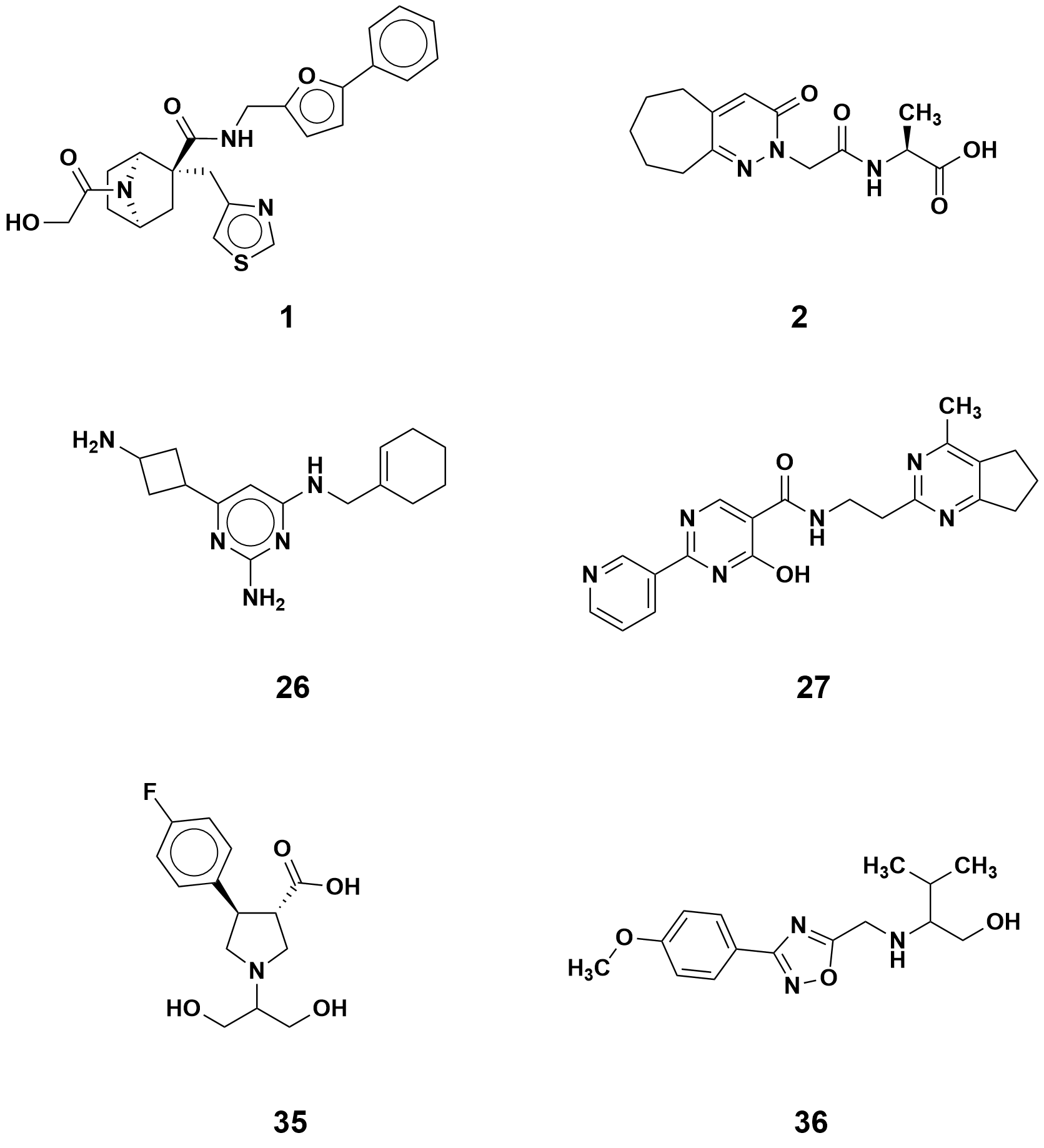
**Supplemental Figure 13. Chemical structures of non-SMM hit and binders used for IL-17, IL-13 and IL-23 Boltz-2 prediction studies.**

**References:**

(1)	Ramírez, J.; Cañete, J. D. Anakinra for the Treatment of Rheumatoid Arthritis: A Safety Evaluation. *Expert Opin Drug Saf* **2018**, *17* (7), 727–732. https://doi.org/10.1080/14740338.2018.1486819.

(2)	Di Paolo, N. C.; Shayakhmetov, D. M. Interleukin 1α and the Inflammatory Process. *Nat Immunol* **2016**, *17* (8), 906–913. https://doi.org/10.1038/ni.3503.

(3)	Cavalli, G.; Colafrancesco, S.; Emmi, G.; Imazio, M.; Lopalco, G.; Maggio, M. C.; Sota, J.; Dinarello, C. A. Interleukin 1α: A Comprehensive Review on the Role of IL-1α in the Pathogenesis and Treatment of Autoimmune and Inflammatory Diseases. *Autoimmunity Reviews* **2021**, *20* (3), 102763. https://doi.org/10.1016/j.autrev.2021.102763.

(4)	Malik, A.; Kanneganti, T.-D. Function and Regulation of IL-1α in Inflammatory Diseases and Cancer. *Immunol Rev* **2018**, *281* (1), 124–137. https://doi.org/10.1111/imr.12615.

(5)	Cron, R. Q.; Caricchio, R.; Chatham, W. W. Calming the Cytokine Storm in COVID-19. *Nat Med* **2021**, *27* (10), 1674–1675. https://doi.org/10.1038/s41591-021-01500-9.

(6)	Kyriazopoulou, E.; Poulakou, G.; Milionis, H.; Metallidis, S.; Adamis, G.; Tsiakos, K.; Fragkou, A.; Rapti, A.; Damoulari, C.; Fantoni, M.; Kalomenidis, I.; Chrysos, G.; Angheben, A.; Kainis, I.; Alexiou, Z.; Castelli, F.; Serino, F. S.; Tsilika, M.; Bakakos, P.; Nicastri, E.; Tzavara, V.; Kostis, E.; Dagna, L.; Koufargyris, P.; Dimakou, K.; Savvanis, S.; Tzatzagou, G.; Chini, M.; Cavalli, G.; Bassetti, M.; Katrini, K.; Kotsis, V.; Tsoukalas, G.; Selmi, C.; Bliziotis, I.; Samarkos, M.; Doumas, M.; Ktena, S.; Masgala, A.; Papanikolaou, I.; Kosmidou, M.; Myrodia, D.-M.; Argyraki, A.; Cardellino, C. S.; Koliakou, K.; Katsigianni, E.-I.; Rapti, V.; Giannitsioti, E.; Cingolani, A.; Micha, S.; Akinosoglou, K.; Liatsis-Douvitsas, O.; Symbardi, S.; Gatselis, N.; Mouktaroudi, M.; Ippolito, G.; Florou, E.; Kotsaki, A.; Netea, M. G.; Eugen-Olsen, J.; Kyprianou, M.; Panagopoulos, P.; Dalekos, G. N.; Giamarellos-Bourboulis, E. J. Early Treatment of COVID-19 with Anakinra Guided by Soluble Urokinase Plasminogen Receptor Plasma Levels: A Double-Blind, Randomized Controlled Phase 3 Trial. *Nat Med* **2021**, *27* (10), 1752–1760. https://doi.org/10.1038/s41591-021-01499-z.

(7)	León, X.; Bothe, C.; García, J.; Parreño, M.; Alcolea, S.; Quer, M.; Vila, L.; Camacho, M. Expression of IL-1α Correlates with Distant Metastasis in Patients with Head and Neck Squamous Cell Carcinoma. *Oncotarget* **2015**, *6* (35), 37398–37409. https://doi.org/10.18632/oncotarget.6054.

(8)	Moran, A.; Bundy, B.; Becker, D. J.; DiMeglio, L. A.; Gitelman, S. E.; Goland, R.; Greenbaum, C. J.; Herold, K. C.; Marks, J. B.; Raskin, P.; Sanda, S.; Schatz, D.; Wherrett, D. K.; Wilson, D. M.; Krischer, J. P.; Skyler, J. S.; Pickersgill, L.; de Koning, E.; Ziegler, A.-G.; Böehm, B.; Badenhoop, K.; Schloot, N.; Bak, J. F.; Pozzilli, P.; Mauricio, D.; Donath, M. Y.; Castaño, L.; Wägner, A.; Lervang, H. H.; Perrild, H.; Poulsen, T. M. Interleukin-1 Antagonism in Type 1 Diabetes of Recent Onset: Two Multicentre, Randomised, Double-Blind, Placebo-Controlled Trials. *Lancet* **2013**, *381* (9881), 10.1016/S0140-6736(13)60023-9. https://doi.org/10.1016/S0140-6736(13)60023-9.

(9)	Ablamunits, V.; Henegariu, O.; Hansen, J. B.; Opare-Addo, L.; Preston-Hurlburt, P.; Santamaria, P.; Mandrup-Poulsen, T.; Herold, K. C. Synergistic Reversal of Type 1 Diabetes in NOD Mice With Anti-CD3 and Interleukin-1 Blockade. *Diabetes* **2012**, *61* (1), 145–154. https://doi.org/10.2337/db11-1033.

(10)	*Autoinflammatory Disease Treatment | ILARIS® (canakinumab)*. https://www.ilaris.com/?site=415726-415729GK100009&utm_source=google&utm_mlr=415726-415729&utm_medium=cpc&utm_campaign=google_branded_415726-415729-ilaris-dtc-branded%3Bs%3Bph%3Bbr%3Bimm%3Bdtc%3Bbr_apr-2024&utm_content=ila_aosd_pfs_sjia_brandaware_n2_brand-general-exact&utm_term=canakinumab&gclid=CjwKCAjwg8qzBhAoEiwAWagLrDdeYFdjnxCYcIPkeTPa56jDExW7-hbn0V-XBAMAPuCOQkEPRXRbfxoCtkUQAvD_BwE&gclsrc=aw.ds (accessed 2024-06-19).

(11)	Zhao, R.; Zhou, H.; Su, S. B. A Critical Role for Interleukin-1β in the Progression of Autoimmune Diseases. *International Immunopharmacology* **2013**, *17* (3), 658–669. https://doi.org/10.1016/j.intimp.2013.08.012.

(12)	Mendiola, A. S.; Cardona, A. E. The IL-1β Phenomena in Neuroinflammatory Diseases. *J Neural Transm (Vienna)* **2018**, *125* (5), 781–795. https://doi.org/10.1007/s00702-017-1732-9.

(13)	Lukens, J. R.; Barr, M. J.; Chaplin, D. D.; Chi, H.; Kanneganti, T.-D. Inflammasome-Derived IL-1β Regulates the Production of GM-CSF by CD4+ T Cells and Γδ T Cells. *The Journal of Immunology* **2012**, *188* (7), 3107–3115. https://doi.org/10.4049/jimmunol.1103308.

(14)	Hondowicz, B. D.; An, D.; Schenkel, J. M.; Kim, K. S.; Steach, H. R.; Krishnamurty, A. T.; Keitany, G. J.; Garza, E. N.; Fraser, K. A.; Moon, J. J.; Altemeier, W. A.; Masopust, D.; Pepper, M. Interleukin-2-Dependent Allergen-Specific Tissue-Resident Memory Cells Drive Asthma. *Immunity* **2016**, *44* (1), 155–166. https://doi.org/10.1016/j.immuni.2015.11.004.

(15)	Edris, A.; De Feyter, S.; Maes, T.; Joos, G.; Lahousse, L. Monoclonal Antibodies in Type 2 Asthma: A Systematic Review and Network Meta-Analysis. *Respir Res* **2019**, *20* (1), 179. https://doi.org/10.1186/s12931-019-1138-3.

(16)	Murakami, Y.; Ishii, T.; Nunokawa, H.; Kurata, K.; Narita, T.; Yamashita, N. TLR9–IL-2 Axis Exacerbates Allergic Asthma by Preventing IL-17A Hyperproduction. *Sci Rep* **2020**, *10* (1), 18110. https://doi.org/10.1038/s41598-020-75153-y.

(17)	Tsai, Y.-C.; Tsai, T.-F. Anti-Interleukin and Interleukin Therapies for Psoriasis: Current Evidence and Clinical Usefulness. *Therapeutic Advances in Musculoskeletal* **2017**, *9* (11), 277–294. https://doi.org/10.1177/1759720X17735756.

(18)	Fala, L. Nucala (Mepolizumab): First IL-5 Antagonist Monoclonal Antibody FDA Approved for Maintenance Treatment of Patients with Severe Asthma. *American Health & Drug Benefits* **2016**, *9* (Spec Feature), 106.

(19)	Commissioner, O. of the. *FDA approves first drug for Eosinophilic Granulomatosis with Polyangiitis, a rare disease formerly known as the Churg-Strauss Syndrome*. FDA. https://www.fda.gov/news-events/press-announcements/fda-approves-first-drug-eosinophilic-granulomatosis-polyangiitis-rare-disease-formerly-known-churg (accessed 2024-10-21).

(20)	*FASENRA approved for treatment of children aged 6 to 11 with severe asthma*. https://www.astrazeneca-us.com/media/press-releases/2024/fasenra-approved-for-treatment-of-children-aged-6-to-11-with-severe-asthma.html (accessed 2024-10-21).

(21)	*Fasenra approved in the US for eosinophilic granulomatosis with polyangiitis*. https://www.astrazeneca.com/media-centre/press-releases/2024/fasenra-approved-in-the-us-for-eosinophilic-granulomatosis-with-polyangiitis.html (accessed 2024-10-21).

(22)	Pelaia, C.; Paoletti, G.; Puggioni, F.; Racca, F.; Pelaia, G.; Canonica, G. W.; Heffler, E. Interleukin-5 in the Pathophysiology of Severe Asthma. *Front Physiol* **2019**, *10*, 1514. https://doi.org/10.3389/fphys.2019.01514.

(23)	Nada, H.; Sivaraman, A.; Lu, Q.; Min, K.; Kim, S.; Goo, J.-I.; Choi, Y.; Lee, K. Perspective for Discovery of Small Molecule IL-6 Inhibitors through Study of Structure–Activity Relationships and Molecular Docking. *J. Med. Chem.* **2023**, *66* (7), 4417–4433. https://doi.org/10.1021/acs.jmedchem.2c01957.

(24)	Scott, L. J. Tocilizumab: A Review in Rheumatoid Arthritis. *Drugs* **2017**, *77* (17), 1865–1879. https://doi.org/10.1007/s40265-017-0829-7.

(25)	Loricera, J.; Blanco, R.; Hernández, J. L.; Castañeda, S.; Mera, A.; Pérez-Pampín, E.; Peiró, E.; Humbría, A.; Calvo-Alén, J.; Aurrecoechea, E.; Narváez, J.; Sánchez-Andrade, A.; Vela, P.; Díez, E.; Mata, C.; Lluch, P.; Moll, C.; Hernández, Í.; Calvo-Río, V.; Ortiz-Sanjuán, F.; González-Vela, C.; Pina, T.; González-Gay, M. Á. Tocilizumab in Giant Cell Arteritis: Multicenter Open-Label Study of 22 Patients. *Semin Arthritis Rheum* **2015**, *44* (6), 717–723. https://doi.org/10.1016/j.semarthrit.2014.12.005.

(26)	Le, R. Q.; Li, L.; Yuan, W.; Shord, S. S.; Nie, L.; Habtemariam, B. A.; Przepiorka, D.; Farrell, A. T.; Pazdur, R. FDA Approval Summary: Tocilizumab for Treatment of Chimeric Antigen Receptor T Cell-Induced Severe or Life-Threatening Cytokine Release Syndrome. *Oncologist* **2018**, *23* (8), 943–947. https://doi.org/10.1634/theoncologist.2018-0028.

(27)	Parums, D. V. Editorial: Tocilizumab, a Humanized Therapeutic IL-6 Receptor (IL-6R) Monoclonal Antibody, and Future Combination Therapies for Severe COVID-19. *Med Sci Monit* **2021**, *27*, e933973. https://doi.org/10.12659/MSM.933973.

(28)	Mospan, G.; Mospan, C.; Vance, S.; Bradshaw, A.; Meosky, K.; Bowles, K. Drug Updates and Approvals: 2017 in Review. *The Nurse Practitioner* **2017**, *42* (12), 8. https://doi.org/10.1097/01.NPR.0000526760.22854.f1.

(29)	Heo, Y.-A. Satralizumab: First Approval. *Drugs* **2020**, *80* (14), 1477–1482. https://doi.org/10.1007/s40265-020-01380-2.

(30)	Mullard, A. 2014 FDA Drug Approvals: The FDA Approved 41 New Therapeutics in 2014, but the Bumper Year Fell Short of the Commercial Power of the Drugs Approved in 2013. *Nature Reviews Drug Discovery* **2015**, *14* (2), 77–82.

(31)	Bartoli, F.; Bae, S.; Cometi, L.; Matucci Cerinic, M.; Furst, D. E. Sirukumab for the Treatment of Rheumatoid Arthritis: Update on Sirukumab, 2018. *Expert Rev Clin Immunol* **2018**, *14* (7), 539–547. https://doi.org/10.1080/1744666X.2018.1487291.

(32)	Wallace, D. J.; Strand, V.; Merrill, J. T.; Popa, S.; Spindler, A. J.; Eimon, A.; Petri, M.; Smolen, J. S.; Wajdula, J.; Christensen, J.; Li, C.; Diehl, A.; Vincent, M. S.; Beebe, J.; Healey, P.; Sridharan, S. Efficacy and Safety of an Interleukin 6 Monoclonal Antibody for the Treatment of Systemic Lupus Erythematosus: A Phase II Dose-Ranging Randomised Controlled Trial. *Annals of the Rheumatic Diseases* **2017**, *76* (3), 534–542. https://doi.org/10.1136/annrheumdis-2016-209668.

(33)	Zhong, H.; Davis, A.; Ouzounova, M.; Carrasco, R. A.; Chen, C.; Breen, S.; Chang, Y. S.; Huang, J.; Liu, Z.; Yao, Y.; Hurt, E.; Moisan, J.; Fung, M.; Tice, D. A.; Clouthier, S. G.; Xiao, Z.; Wicha, M. S.; Korkaya, H.; Hollingsworth, R. E. A Novel IL6 Antibody Sensitizes Multiple Tumor Types to Chemotherapy Including Trastuzumab-Resistant Tumors. *Cancer Res* **2016**, *76* (2), 480–490. https://doi.org/10.1158/0008-5472.CAN-15-0883.

(34)	Moore, J. B.; June, C. H. Cytokine Release Syndrome in Severe COVID-19. *Science* **2020**, *368* (6490), 473–474. https://doi.org/10.1126/science.abb8925.

(35)	Akkapeddi, P.; Fragoso, R.; Hixon, J. A.; Ramalho, A. S.; Oliveira, M. L.; Carvalho, T.; Gloger, A.; Matasci, M.; Corzana, F.; Durum, S. K.; Neri, D.; Bernardes, G. J. L.; Barata, J. T. A Fully Human Anti-IL-7Rα Antibody Promotes Antitumor Activity against T-Cell Acute Lymphoblastic Leukemia. *Leukemia* **2019**, *33* (9), 2155–2168. https://doi.org/10.1038/s41375-019-0434-8.

(36)	Herold, K. C.; Bucktrout, S. L.; Wang, X.; Bode, B. W.; Gitelman, S. E.; Gottlieb, P. A.; Hughes, J.; Joh, T.; McGill, J. B.; Pettus, J. H.; Potluri, S.; Schatz, D.; Shannon, M.; Udata, C.; Wong, G.; Levisetti, M.; Ganguly, B. J.; Garzone, P. D.; RN168 Working Group. Immunomodulatory Activity of Humanized Anti-IL-7R Monoclonal Antibody RN168 in Subjects with Type 1 Diabetes. *JCI Insight* **2019**, *4* (24), e126054, 126054. https://doi.org/10.1172/jci.insight.126054.

(37)	Bikker, A.; Hack, C. E.; Lafeber, F. P. J. G.; Roon, J. A. G. van. Interleukin-7: A Key Mediator in T Cell-Driven Autoimmunity, Inflammation, and Tissue Destruction. *Current Pharmaceutical Design* *18* (16), 2347–2356.

(38)	Meyer, A.; Parmar, P. J.; Shahrara, S. Significance of IL-7 and IL-7R in RA and Autoimmunity. *Autoimmunity Reviews* **2022**, *21* (7), 103120. https://doi.org/10.1016/j.autrev.2022.103120.

(39)	Zeng, Z.; Mao, H.; Lei, Q.; He, Y. IL-7 in Autoimmune Diseases: Mechanisms and Therapeutic Potential. *Front Immunol* **2025**, *16*, 1545760. https://doi.org/10.3389/fimmu.2025.1545760.

(40)	Chen, D.; Tang, T.-X.; Deng, H.; Yang, X.-P.; Tang, Z.-H. Interleukin-7 Biology and Its Effects on Immune Cells: Mediator of Generation, Differentiation, Survival, and Homeostasis. *Front Immunol* **2021**, *12*, 747324. https://doi.org/10.3389/fimmu.2021.747324.

(41)	Liu, Q.; Li, A.; Tian, Y.; Wu, J. D.; Liu, Y.; Li, T.; Chen, Y.; Han, X.; Wu, K. The CXCL8-CXCR1/2 Pathways in Cancer. *Cytokine & Growth Factor Reviews* **2016**, *31*, 61–71. https://doi.org/10.1016/j.cytogfr.2016.08.002.

(42)	Xiong, X.; Liao, X.; Qiu, S.; Xu, H.; Zhang, S.; Wang, S.; Ai, J.; Yang, L. CXCL8 in Tumor Biology and Its Implications for Clinical Translation. *Front Mol Biosci* **2022**, *9*, 723846. https://doi.org/10.3389/fmolb.2022.723846.

(43)	Waugh, D. J. J.; Wilson, C. The Interleukin-8 Pathway in Cancer. *Clin Cancer Res* **2008**, *14* (21), 6735–6741. https://doi.org/10.1158/1078-0432.CCR-07-4843.

(44)	Bilusic, M.; Heery, C. R.; Collins, J. M.; Donahue, R. N.; Palena, C.; Madan, R. A.; Karzai, F.; Marté, J. L.; Strauss, J.; Gatti-Mays, M. E.; Schlom, J.; Gulley, J. L. Phase I Trial of HuMax-IL8 (BMS-986253), an Anti-IL-8 Monoclonal Antibody, in Patients with Metastatic or Unresectable Solid Tumors. *J Immunother Cancer* **2019**, *7*, 240. https://doi.org/10.1186/s40425-019-0706-x.

(45)	Wang, B.; Roskos, L.; Osborn, K.; Lu, H.; Kim, M.; Pasumarthi, R.; Raie, N.; Tabrizi, M. A Population Pharmacokinetic (PK) Analysis of ABX-IL8, a Fully Human Monoclononal IGG2 Antibody, in Psoriasis Patients. *Clinical Pharmacology & Therapeutics* **2005**, *77* (2), P91–P91. https://doi.org/10.1016/j.clpt.2004.12.240.

(46)	Goldstein, L. J.; Mansutti, M.; Levy, C.; Chang, J. C.; Henry, S.; Fernandez-Perez, I.; Prausovà, J.; Staroslawska, E.; Viale, G.; Butler, B.; McCanna, S.; Ruffini, P. A.; Wicha, M. S.; Schott, A. F.; fRida Trial Investigators. A Randomized, Placebo-Controlled Phase 2 Study of Paclitaxel in Combination with Reparixin Compared to Paclitaxel Alone as Front-Line Therapy for Metastatic Triple-Negative Breast Cancer (fRida). *Breast Cancer Res Treat* **2021**, *190* (2), 265–275. https://doi.org/10.1007/s10549-021-06367-5.

(47)	Saraiva, M.; Vieira, P.; O’Garra, A. Biology and Therapeutic Potential of Interleukin-10. *J Exp Med* **2019**, *217* (1), e20190418. https://doi.org/10.1084/jem.20190418.

(48)	Tian, G.; Li, J.-L.; Wang, D.-G.; Zhou, D. Targeting IL-10 in Auto-Immune Diseases. *Cell Biochem Biophys* **2014**, *70* (1), 37–49. https://doi.org/10.1007/s12013-014-9903-x.

(49)	Biotest. *A Prospective, Double-Blind, Randomized, Placebo-Controlled, Repeated Dose, Multicentre Phase IIa Proof-of-Concept Study With BT063 in Subjects With Systemic Lupus Erythematosus*; Clinical trial registration NCT02554019; clinicaltrials.gov, 2020. https://clinicaltrials.gov/study/NCT02554019 (accessed 2023-12-31).

(50)	Seyedsadr, M.; Wang, Y.; Elzoheiry, M.; Shree Gopal, S.; Jang, S.; Duran, G.; Chervoneva, I.; Kasimoglou, E.; Wrobel, J. A.; Hwang, D.; Garifallou, J.; Zhang, X.; Khan, T. H.; Lorenz, U.; Su, M.; Ting, J. P.; Broux, B.; Rostami, A.; Miskin, D.; Markovic-Plese, S. IL-11 Induces NLRP3 Inflammasome Activation in Monocytes and Inflammatory Cell Migration to the Central Nervous System. *Proceedings of the National Academy of Sciences* **2023**, *120* (26), e2221007120. https://doi.org/10.1073/pnas.2221007120.

(51)	Ng, B.; Cook, S. A.; Schafer, S. Interleukin-11 Signaling Underlies Fibrosis, Parenchymal Dysfunction, and Chronic Inflammation of the Airway. *Exp Mol Med* **2020**, *52* (12), 1871–1878. https://doi.org/10.1038/s12276-020-00531-5.

(52)	Zhang, X.; Kiapour, N.; Kapoor, S.; Merrill, J. R.; Xia, Y.; Ban, W.; Cohen, S. M.; Midkiff, B. R.; Jewells, V.; Shih, Y.-Y. I.; Markovic-Plese, S. IL-11 Antagonist Suppresses Th17 Cell-Mediated Neuroinflammation and Demyelination in a Mouse Model of Relapsing-Remitting Multiple Sclerosis. *Clin Immunol* **2018**, *197*, 45–53. https://doi.org/10.1016/j.clim.2018.08.006.

(53)	Fung, K. Y.; Louis, C.; Metcalfe, R. D.; Kosasih, C. C.; Wicks, I. P.; Griffin, M. D. W.; Putoczki, T. L. Emerging Roles for IL-11 in Inflammatory Diseases. *Cytokine* **2022**, *149*, 155750. https://doi.org/10.1016/j.cyto.2021.155750.

(54)	Gurfein, B. T.; Zhang, Y.; López, C. B.; Argaw, A. T.; Zameer, A.; Moran, T. M.; John, G. R. IL-11 Regulates Autoimmune Demyelination. *J Immunol* **2009**, *183* (7), 4229–4240. https://doi.org/10.4049/jimmunol.0900622.

(55)	Marwaha, A. K.; Chow, S.; Pesenacker, A. M.; Cook, L.; Sun, A.; Long, S. A.; Yang, J. H. M.; Ward-Hartstonge, K. A.; Williams, E.; Domingo-Vila, C.; Halani, K.; Harris, K. M.; Tree, T. I. M.; Levings, M. K.; Elliott, T.; Tan, R.; Dutz, J. P. A Phase 1b Open-Label Dose-Finding Study of Ustekinumab in Young Adults with Type 1 Diabetes. *Immunotherapy Advances* **2022**, *2* (1), ltab022. https://doi.org/10.1093/immadv/ltab022.

(56)	Segal, B. M.; Constantinescu, C. S.; Raychaudhuri, A.; Kim, L.; Fidelus-Gort, R.; Kasper, L. H.; Ustekinumab MS Investigators. Repeated Subcutaneous Injections of IL12/23 P40 Neutralising Antibody, Ustekinumab, in Patients with Relapsing-Remitting Multiple Sclerosis: A Phase II, Double-Blind, Placebo-Controlled, Randomised, Dose-Ranging Study. *Lancet Neurol* **2008**, *7* (9), 796–804. https://doi.org/10.1016/S1474-4422(08)70173-X.

(57)	Becher, B.; Durell, B. G.; Noelle, R. J. Experimental Autoimmune Encephalitis and Inflammation in the Absence of Interleukin-12. *J Clin Invest* **2002**, *110* (4), 493–497. https://doi.org/10.1172/JCI15751.

(58)	Haskó, G.; Szabó, C. IL-12 as a Therapeutic Target for Pharmacological Modulation in Immune-Mediated and Inflammatory Diseases: Regulation of T Helper 1/T Helper 2 Responses. *Br J Pharmacol* **1999**, *127* (6), 1295–1304. https://doi.org/10.1038/sj.bjp.0702689.

(59)	Hamza, T.; Barnett, J. B.; Li, B. Interleukin 12 a Key Immunoregulatory Cytokine in Infection Applications. *Int J Mol Sci* **2010**, *11* (3), 789–806. https://doi.org/10.3390/ijms11030789.

(60)	Tian, Z.; Zhao, Q.; Teng, X. Anti-IL23/12 Agents and JAK Inhibitors for Inflammatory Bowel Disease. *Front. Immunol.* **2024**, *15*. https://doi.org/10.3389/fimmu.2024.1393463.

(61)	Greving, C. N. A.; Towne, J. E. A Role for IL-12 in IBD after All? *Immunity* **2019**, *51* (2), 209–211. https://doi.org/10.1016/j.immuni.2019.07.008.

(62)	*FDA Approves Lilly’s EBGLYSS^TM^ (lebrikizumab-lbkz) for Adults and Children 12 Years and Older with Moderate-to-Severe Atopic Dermatitis | Eli Lilly and Company*. https://investor.lilly.com/news-releases/news-release-details/fda-approves-lillys-ebglysstm-lebrikizumab-lbkz-adults-and (accessed 2024-10-21).

(63)	Ultsch, M.; Bevers, J.; Nakamura, G.; Vandlen, R.; Kelley, R. F.; Wu, L. C.; Eigenbrot, C. Structural Basis of Signaling Blockade by Anti-IL-13 Antibody Lebrikizumab. *Journal of Molecular Biology* **2013**, *425* (8), 1330–1339. https://doi.org/10.1016/j.jmb.2013.01.024.

(64)	Association, N. E. *LEO Pharma Inc. Announces U.S. FDA approval of Adbry® (tralokinumab-ldrm) for the Treatment of Moderate-to-severe Atopic Dermatitis in Pediatric Patients Aged 12-17 Years*. National Eczema Association. https://nationaleczema.org/blog/leo-121523/ (accessed 2024-10-21).

(65)	*FDA Approves Lebrikizumab Treatment of Eczema Among Patients Aged 12 and Older*. Dermatology | GW School of Medicine and Health Sciences. https://dermatology.smhs.gwu.edu/news/fda-approves-lebrikizumab-treatment-eczema-among-patients-aged-12-and-older (accessed 2024-10-21).

(66)	Bagnasco, D.; Ferrando, M.; Varricchi, G.; Passalacqua, G.; Canonica, G. W. A Critical Evaluation of Anti-IL-13 and Anti-IL-4 Strategies in Severe Asthma. *Int Arch Allergy Immunol* **2016**, *170* (2), 122–131. https://doi.org/10.1159/000447692.

(67)	Rabe, K. F.; Nair, P.; Brusselle, G.; Maspero, J. F.; Castro, M.; Sher, L.; Zhu, H.; Hamilton, J. D.; Swanson, B. N.; Khan, A.; Chao, J.; Staudinger, H.; Pirozzi, G.; Antoni, C.; Amin, N.; Ruddy, M.; Akinlade, B.; Graham, N. M. H.; Stahl, N.; Yancopoulos, G. D.; Teper, A. Efficacy and Safety of Dupilumab in Glucocorticoid-Dependent Severe Asthma. *N Engl J Med* **2018**, *378* (26), 2475–2485. https://doi.org/10.1056/NEJMoa1804093.

(68)	Castro, M.; Corren, J.; Pavord, I. D.; Maspero, J.; Wenzel, S.; Rabe, K. F.; Busse, W. W.; Ford, L.; Sher, L.; FitzGerald, J. M.; Katelaris, C.; Tohda, Y.; Zhang, B.; Staudinger, H.; Pirozzi, G.; Amin, N.; Ruddy, M.; Akinlade, B.; Khan, A.; Chao, J.; Martincova, R.; Graham, N. M. H.; Hamilton, J. D.; Swanson, B. N.; Stahl, N.; Yancopoulos, G. D.; Teper, A. Dupilumab Efficacy and Safety in Moderate-to-Severe Uncontrolled Asthma. *N Engl J Med* **2018**, *378* (26), 2486–2496. https://doi.org/10.1056/NEJMoa1804092.

(69)	Knudson, K. M.; Hwang, S.; McCann, M. S.; Joshi, B. H.; Husain, S. R.; Puri, R. K. Recent Advances in IL-13Rα2-Directed Cancer Immunotherapy. *Front Immunol* **2022**, *13*, 878365. https://doi.org/10.3389/fimmu.2022.878365.

(70)	Huangfu, L.; Li, R.; Huang, Y.; Wang, S. The IL-17 Family in Diseases: From Bench to Bedside. *Sig Transduct Target Ther* **2023**, *8* (1), 1–22. https://doi.org/10.1038/s41392-023-01620-3.

(71)	Blegvad, C.; Skov, L.; Zachariae, C. Ixekizumab for the Treatment of Psoriasis: An Update on New Data since First Approval. *Expert Rev Clin Immunol* **2019**, *15* (2), 111–121. https://doi.org/10.1080/1744666X.2019.1559730.

(72)	Commissioner, O. of the. *FDA approves new psoriasis drug Taltz*. FDA. https://www.fda.gov/news-events/press-announcements/fda-approves-new-psoriasis-drug-taltz (accessed 2024-10-21).

(73)	Di Lernia, V.; Bombonato, C.; Motolese, A. COVID-19 in an Elderly Patient Treated with Secukinumab. *Dermatol Ther* **2020**, *33* (4), e13580. https://doi.org/10.1111/dth.13580.

(74)	Komiyama, Y.; Nakae, S.; Matsuki, T.; Nambu, A.; Ishigame, H.; Kakuta, S.; Sudo, K.; Iwakura, Y. IL-17 Plays an Important Role in the Development of Experimental Autoimmune Encephalomyelitis. *J Immunol* **2006**, *177* (1), 566–573. https://doi.org/10.4049/jimmunol.177.1.566.

(75)	Havrdová, E.; Belova, A.; Goloborodko, A.; Tisserant, A.; Wright, A.; Wallstroem, E.; Garren, H.; Maguire, R. P.; Johns, D. R. Activity of Secukinumab, an Anti-IL-17A Antibody, on Brain Lesions in RRMS: Results from a Randomized, Proof-of-Concept Study. *J Neurol* **2016**, *263* (7), 1287–1295. https://doi.org/10.1007/s00415-016-8128-x.

(76)	Emamaullee, J. A.; Davis, J.; Merani, S.; Toso, C.; Elliott, J. F.; Thiesen, A.; Shapiro, A. M. J. Inhibition of Th17 Cells Regulates Autoimmune Diabetes in NOD Mice. *Diabetes* **2009**, *58* (6), 1302–1311. https://doi.org/10.2337/db08-1113.

(77)	Abdel-Moneim, A.; Bakery, H. H.; Allam, G. The Potential Pathogenic Role of IL-17/Th17 Cells in Both Type 1 and Type 2 Diabetes Mellitus. *Biomedicine & Pharmacotherapy* **2018**, *101*, 287–292. https://doi.org/10.1016/j.biopha.2018.02.103.

(78)	Li, C.-R.; Mueller, E. E.; Bradley, L. M. Islet Antigen-Specific Th17 Cells Can Induce TNFα-Dependent Autoimmune Diabetes. *J Immunol* **2014**, *192* (4), 1425–1432. https://doi.org/10.4049/jimmunol.1301742.

(79)	Chao, C.-C.; Chen, S.-J.; Adamopoulos, I. E.; Davis, N.; Hong, K.; Vu, A.; Kwan, S.; Fayadat-Dilman, L.; Asio, A.; Bowman, E. P. Anti-IL-17A Therapy Protects against Bone Erosion in Experimental Models of Rheumatoid Arthritis. *Autoimmunity* **2011**, *44* (3), 243–252. https://doi.org/10.3109/08916934.2010.517815.

(80)	Dayer, J.-M. Interleukin-18, Rheumatoid Arthritis, and Tissue Destruction. *J Clin Invest* **1999**, *104* (10), 1337–1339. https://doi.org/10.1172/JCI8731.

(81)	Gracie, J. A.; Forsey, R. J.; Chan, W. L.; Gilmour, A.; Leung, B. P.; Greer, M. R.; Kennedy, K.; Carter, R.; Wei, X.-Q.; Xu, D.; Field, M.; Foulis, A.; Liew, F. Y.; McInnes, I. B. A Proinflammatory Role for IL-18 in Rheumatoid Arthritis. *J Clin Invest* **1999**, *104* (10), 1393–1401. https://doi.org/10.1172/JCI7317.

(82)	Morel, J. C.; Park, C. C.; Woods, J. M.; Koch, A. E. A Novel Role for Interleukin-18 in Adhesion Molecule Induction through NF Kappa B and Phosphatidylinositol (PI) 3-Kinase-Dependent Signal Transduction Pathways. *J Biol Chem* **2001**, *276* (40), 37069–37075. https://doi.org/10.1074/jbc.M103574200.

(83)	Liu, S. M.; Lee, D. H.; Sullivan, J. M.; Chung, D.; Jäger, A.; Shum, B. O. V.; Sarvetnick, N. E.; Anderson, A. C.; Kuchroo, V. K. Differential IL-21 Signaling in APCs Leads to Disparate Th17 Differentiation in Diabetes-Susceptible NOD and Diabetes-Resistant NOD.Idd3 Mice. *J Clin Invest* **2011**, *121* (11), 4303–4310. https://doi.org/10.1172/JCI46187.

(84)	Van Belle, T. L.; Nierkens, S.; Arens, R.; von Herrath, M. G. Interleukin-21 Receptor-Mediated Signals Control Autoreactive T Cell Infiltration in Pancreatic Islets. *Immunity* **2012**, *36* (6), 10.1016/j.immuni.2012.04.005. https://doi.org/10.1016/j.immuni.2012.04.005.

(85)	Sutherland, A. P. R.; Van Belle, T.; Wurster, A. L.; Suto, A.; Michaud, M.; Zhang, D.; Grusby, M. J.; von Herrath, M. Interleukin-21 Is Required for the Development of Type 1 Diabetes in NOD Mice. *Diabetes* **2009**, *58* (5), 1144–1155. https://doi.org/10.2337/db08-0882.

(86)	Long, D.; Chen, Y.; Wu, H.; Zhao, M.; Lu, Q. Clinical Significance and Immunobiology of IL-21 in Autoimmunity. *Journal of Autoimmunity* **2019**, *99*, 1–14. https://doi.org/10.1016/j.jaut.2019.01.013.

(87)	Yi, C.; Yi, Y.; Wei, J.; Jin, Q.; Li, J.; Sacitharan, P. K. Targeting IL-22 and IL-22R Protects against Experimental Osteoarthritis. *Cell Mol Immunol* **2021**, *18* (5), 1329–1331. https://doi.org/10.1038/s41423-020-0491-y.

(88)	Pan, H.-F.; Li, X.-P.; Zheng, S. G.; Ye, D.-Q. Emerging Role of Interleukin-22 in Autoimmune Diseases. *Cytokine Growth Factor Rev* **2013**, *24* (1), 51–57. https://doi.org/10.1016/j.cytogfr.2012.07.002.

(89)	Marijnissen, R. J.; Koenders, M. I.; Smeets, R. L.; Stappers, M. H. T.; Nickerson-Nutter, C.; Joosten, L. A. B.; Boots, A. M. H.; van den Berg, W. B. Increased Expression of Interleukin-22 by Synovial Th17 Cells during Late Stages of Murine Experimental Arthritis Is Controlled by Interleukin-1 and Enhances Bone Degradation. *Arthritis & Rheumatism* **2011**, *63* (10), 2939–2948. https://doi.org/10.1002/art.30469.

(90)	Lindahl, H.; Olsson, T. Interleukin-22 Influences the Th1/Th17 Axis. *Front Immunol* **2021**, *12*, 618110. https://doi.org/10.3389/fimmu.2021.618110.

(91)	Colquhoun, M.; Kemp, A. K. Ustekinumab. In *StatPearls*; StatPearls Publishing: Treasure Island (FL), 2024.

(92)	Lee, K. M.-C.; Sherlock, J. P.; Hamilton, J. A. The Role of Interleukin (IL)-23 in Regulating Pain in Arthritis. *Arthritis Res Ther* **2022**, *24* (1), 89. https://doi.org/10.1186/s13075-022-02777-y.

(93)	Tang, C.; Chen, S.; Qian, H.; Huang, W. Interleukin-23: As a Drug Target for Autoimmune Inflammatory Diseases. *Immunology* **2012**, *135* (2), 112–124. https://doi.org/10.1111/j.1365-2567.2011.03522.x.

(94)	Cua, D. J.; Sherlock, J.; Chen, Y.; Murphy, C. A.; Joyce, B.; Seymour, B.; Lucian, L.; To, W.; Kwan, S.; Churakova, T.; Zurawski, S.; Wiekowski, M.; Lira, S. A.; Gorman, D.; Kastelein, R. A.; Sedgwick, J. D. Interleukin-23 Rather than Interleukin-12 Is the Critical Cytokine for Autoimmune Inflammation of the Brain. *Nature* **2003**, *421* (6924), 744–748. https://doi.org/10.1038/nature01355.

(95)	Oboki, K.; Ohno, T.; Kajiwara, N.; Arae, K.; Morita, H.; Ishii, A.; Nambu, A.; Abe, T.; Kiyonari, H.; Matsumoto, K.; Sudo, K.; Okumura, K.; Saito, H.; Nakae, S. IL-33 Is a Crucial Amplifier of Innate Rather than Acquired Immunity. *Proceedings of the National Academy of Sciences* **2010**, *107* (43), 18581–18586. https://doi.org/10.1073/pnas.1003059107.

(96)	Li, Y.; Fu, Y.; Chen, H.; Liu, X.; Li, M. Blocking Interleukin-33 Alleviates the Joint Inflammation and Inhibits the Development of Collagen-Induced Arthritis in Mice. *J Immunol Res* **2020**, *2020*, 4297354. https://doi.org/10.1155/2020/4297354.

(97)	Cayrol, C.; Girard, J.-P. Interleukin-33 (IL-33): A Nuclear Cytokine from the IL-1 Family. *Immunological Reviews* **2018**, *281* (1), 154–168. https://doi.org/10.1111/imr.12619.

(98)	Wechsler, M. E.; Ruddy, M. K.; Pavord, I. D.; Israel, E.; Rabe, K. F.; Ford, L. B.; Maspero, J. F.; Abdulai, R. M.; Hu, C.-C.; Martincova, R.; Jessel, A.; Nivens, M. C.; Amin, N.; Weinreich, D. M.; Yancopoulos, G. D.; Goulaouic, H. Efficacy and Safety of Itepekimab in Patients with Moderate-to-Severe Asthma. *New England Journal of Medicine* **2021**, *385* (18), 1656–1668. https://doi.org/10.1056/NEJMoa2024257.

(99)	Ding, W.; Zou, G.-L.; Zhang, W.; Lai, X.-N.; Chen, H.-W.; Xiong, L.-X. Interleukin-33: Its Emerging Role in Allergic Diseases. *Molecules* **2018**, *23* (7), 1665. https://doi.org/10.3390/molecules23071665.

(100)	Miller, A. M. Role of IL-33 in Inflammation and Disease. *J Inflamm* **2011**, *8* (1), 22. https://doi.org/10.1186/1476-9255-8-22.

(101)	Cayrol, C.; Girard, J.-P. Interleukin-33 (IL-33): A Critical Review of Its Biology and the Mechanisms Involved in Its Release as a Potent Extracellular Cytokine. *Cytokine* **2022**, *156*, 155891. https://doi.org/10.1016/j.cyto.2022.155891.

(102)	Karnell, J. L.; Rieder, S. A.; Ettinger, R.; Kolbeck, R. Targeting the CD40-CD40L Pathway in Autoimmune Diseases: Humoral Immunity and Beyond. *Advanced Drug Delivery Reviews* **2019**, *141*, 92–103. https://doi.org/10.1016/j.addr.2018.12.005.

(103)	Karnell, J. L.; Albulescu, M.; Drabic, S.; Wang, L.; Moate, R.; Baca, M.; Oganesyan, V.; Gunsior, M.; Thisted, T.; Yan, L.; Li, J.; Xiong, X.; Eck, S. C.; de los Reyes, M.; Yusuf, I.; Streicher, K.; Müller-Ladner, U.; Howe, D.; Ettinger, R.; Herbst, R.; Drappa, J. A CD40L-Targeting Protein Reduces Autoantibodies and Improves Disease Activity in Patients with Autoimmunity. *Science Translational Medicine* **2019**, *11* (489), eaar6584. https://doi.org/10.1126/scitranslmed.aar6584.

(104)	Danese, S.; Sans, M.; Fiocchi, C. The CD40/CD40L Costimulatory Pathway in Inflammatory Bowel Disease. *Gut* **2004**, *53* (7), 1035–1043. https://doi.org/10.1136/gut.2003.026278.

(105)	Senhaji, N.; Kojok, K.; Darif, Y.; Fadainia, C.; Zaid, Y. The Contribution of CD40/CD40L Axis in Inflammatory Bowel Disease: An Update. *Front. Immunol.* **2015**, *6*. https://doi.org/10.3389/fimmu.2015.00529.

(106)	Mabrouk, M.; Wahnou, H.; Merhi, Y.; Abou-Saleh, H.; Guessous, F.; Zaid, Y. The Role of Soluble CD40L in Autoimmune Diseases. *J Transl Autoimmun* **2025**, *10*, 100288. https://doi.org/10.1016/j.jtauto.2025.100288.

(107)	Pucino, V.; Gardner, D. H.; Fisher, B. A. Rationale for CD40 Pathway Blockade in Autoimmune Rheumatic Disorders. *The Lancet Rheumatology* **2020**, *2* (5), e292–e301. https://doi.org/10.1016/S2665-9913(20)30038-2.

(108)	Alexion Pharmaceuticals, Inc. *A Randomized, Double-Blind, Placebo-Controlled, Multi-Center Trial to Evaluate the Safety and Efficacy of Eculizumab in Patients With Relapsing Neuromyelitis Optica (NMO)*; Clinical trial registration NCT01892345; clinicaltrials.gov, 2019. https://clinicaltrials.gov/study/NCT01892345 (accessed 2023-12-31).

(109)	Walport, M. J. Complement and Systemic Lupus Erythematosus. *Arthritis Res* **2002**, *4* (Suppl 3), S279–S293. https://doi.org/10.1186/ar586.

(110)	Thurman, J. M.; Yapa, R. Complement Therapeutics in Autoimmune Disease. *Front. Immunol.* **2019**, *10*. https://doi.org/10.3389/fimmu.2019.00672.

(111)	Cantarelli, C.; Leventhal, J.; Cravedi, P. Complement in Lupus: Biomarker, Therapeutic Target, or a Little Bit of Both? *Kidney Int Rep* **2021**, *6* (8), 2031–2032. https://doi.org/10.1016/j.ekir.2021.06.016.

(112)	Li, N. L.; Birmingham, D. J.; Rovin, B. H. Expanding the Role of Complement Therapies: The Case for Lupus Nephritis. *J Clin Med* **2021**, *10* (4), 626. https://doi.org/10.3390/jcm10040626.

(113)	Dmytrijuk, A.; Robie-Suh, K.; Cohen, M. H.; Rieves, D.; Weiss, K.; Pazdur, R. FDA Report: Eculizumab (Soliris) for the Treatment of Patients with Paroxysmal Nocturnal Hemoglobinuria. *Oncologist* **2008**, *13* (9), 993–1000. https://doi.org/10.1634/theoncologist.2008-0086.

(114)	McQualter, J. L.; Darwiche, R.; Ewing, C.; Onuki, M.; Kay, T. W.; Hamilton, J. A.; Reid, H. H.; Bernard, C. C. A. Granulocyte Macrophage Colony-Stimulating Factor. *J Exp Med* **2001**, *194* (7), 873–882. https://doi.org/10.1084/jem.194.7.873.

(115)	Constantinescu, C. S.; Asher, A.; Fryze, W.; Kozubski, W.; Wagner, F.; Aram, J.; Tanasescu, R.; Korolkiewicz, R. P.; Dirnberger-Hertweck, M.; Steidl, S.; Libretto, S. E.; Sprenger, T.; Radue, E. W. Randomized Phase 1b Trial of MOR103, a Human Antibody to GM-CSF, in Multiple Sclerosis. *Neurology Neuroimmunology & Neuroinflammation* **2015**, *2* (4), e117. https://doi.org/10.1212/NXI.0000000000000117.

(116)	Palle, P.; Monaghan, K. L.; Milne, S. M.; Wan, E. C. K. Cytokine Signaling in Multiple Sclerosis and Its Therapeutic Applications. *Med Sci (Basel)* **2017**, *5* (4), 23. https://doi.org/10.3390/medsci5040023.

(117)	*GSK announces phase III start for its anti GM-CSF antibody, otilimab, in patients with rheumatoid arthritis (RA) | GSK*. https://www.gsk.com/en-gb/media/press-releases/gsk-announces-phase-iii-start-for-its-anti-gm-csf-antibody-otilimab-in-patients-with-rheumatoid-arthritis-ra/ (accessed 2025-11-30).

(118)	Huizinga, T. W. J.; Batalov, A.; Stoilov, R.; Lloyd, E.; Wagner, T.; Saurigny, D.; Souberbielle, B.; Esfandiari, E. Phase 1b Randomized, Double-Blind Study of Namilumab, an Anti-Granulocyte Macrophage Colony-Stimulating Factor Monoclonal Antibody, in Mild-to-Moderate Rheumatoid Arthritis. *Arthritis Research & Therapy* **2017**, *19* (1), 53. https://doi.org/10.1186/s13075-017-1267-3.

(119)	*GSK presents new efficacy and safety data of an anti GM-CSF antibody in patients with rheumatoid arthritis | GSK*. https://www.gsk.com/en-gb/media/press-releases/gsk-presents-new-efficacy-and-safety-data-of-an-anti-gm-csf-antibody-in-patients-with-rheumatoid-arthritis/ (accessed 2025-11-30).

(120)	Bhattacharya, P.; Budnick, I.; Singh, M.; Thiruppathi, M.; Alharshawi, K.; Elshabrawy, H.; Holterman, M. J.; Prabhakar, B. S. Dual Role of GM-CSF as a Pro-Inflammatory and a Regulatory Cytokine: Implications for Immune Therapy. *J Interferon Cytokine Res* **2015**, *35* (8), 585–599. https://doi.org/10.1089/jir.2014.0149.

(121)	Shiomi, A.; Usui, T.; Mimori, T. GM-CSF as a Therapeutic Target in Autoimmune Diseases. *Inflamm Regen* **2016**, *36*, 8. https://doi.org/10.1186/s41232-016-0014-5.

(122)	Lotfi, N.; Thome, R.; Rezaei, N.; Zhang, G.-X.; Rezaei, A.; Rostami, A.; Esmaeil, N. Roles of GM-CSF in the Pathogenesis of Autoimmune Diseases: An Update. *Department of Neurology Faculty Papers* **2019**, *10*, 1265.

(123)	Lawlor, K. E.; Campbell, I. K.; Metcalf, D.; O’Donnell, K.; van Nieuwenhuijze, A.; Roberts, A. W.; Wicks, I. P. Critical Role for Granulocyte Colony-Stimulating Factor in Inflammatory Arthritis. *Proceedings of the National Academy of Sciences* **2004**, *101* (31), 11398–11403. https://doi.org/10.1073/pnas.0404328101.

(124)	Hamilton, J. A.; Cook, A. D.; Tak, P. P. Anti-Colony-Stimulating Factor Therapies for Inflammatory and Autoimmune Diseases. *Nat Rev Drug Discov* **2017**, *16* (1), 53–70. https://doi.org/10.1038/nrd.2016.231.

(125)	Scalzo-Inguanti, K.; Monaghan, K.; Edwards, K.; Herzog, E.; Mirosa, D.; Hardy, M.; Sorto, V.; Huynh, H.; Rakar, S.; Kurtov, D.; Braley, H.; Wilson, N.; Busfield, S.; Nash, A.; Andrews, A. A Neutralizing Anti-G-CSFR Antibody Blocks G-CSF-Induced Neutrophilia without Inducing Neutropenia in Nonhuman Primates. *J Leukoc Biol* **2017**, *102* (2), 537–549. https://doi.org/10.1189/jlb.5A1116-489R.

(126)	Scalzo-Inguanti, K.; Monaghan, K.; Edwards, K.; Taylor, S.; Nash, A.; Andrews, A.; Busfield, S. 164: CSL324, a Humanised Anti G-CSFR Antibody, Can Inhibit Neutrophil Migration While Not Impaction on Neutrophil Number or Effector Functions. *Cytokine* **2014**, *70* (1), 68. https://doi.org/10.1016/j.cyto.2014.07.171.

(127)	Murthy, H.; Iqbal, M.; Chavez, J. C.; Kharfan-Dabaja, M. A. Cytokine Release Syndrome: Current Perspectives. *Immunotargets Ther* **2019**, *8*, 43–52. https://doi.org/10.2147/ITT.S202015.

(128)	Fox, R. J.; Coffey, C. S.; Conwit, R.; Cudkowicz, M. E.; Gleason, T.; Goodman, A.; Klawiter, E. C.; Matsuda, K.; McGovern, M.; Naismith, R. T.; Ashokkumar, A.; Barnes, J.; Ecklund, D.; Klingner, E.; Koepp, M.; Long, J. D.; Natarajan, S.; Thornell, B.; Yankey, J.; Bermel, R. A.; Debbins, J. P.; Huang, X.; Jagodnik, P.; Lowe, M. J.; Nakamura, K.; Narayanan, S.; Sakaie, K. E.; Thoomukuntla, B.; Zhou, X.; Krieger, S.; Alvarez, E.; Apperson, M.; Bashir, K.; Cohen, B. A.; Coyle, P. K.; Delgado, S.; Dewitt, L. D.; Flores, A.; Giesser, B. S.; Goldman, M. D.; Jubelt, B.; Lava, N.; Lynch, S. G.; Moses, H.; Ontaneda, D.; Perumal, J. S.; Racke, M.; Repovic, P.; Riley, C. S.; Severson, C.; Shinnar, S.; Suski, V.; Weinstock-Guttman, B.; Yadav, V.; Zabeti, A. Phase 2 Trial of Ibudilast in Progressive Multiple Sclerosis. *New England Journal of Medicine* **2018**, *379* (9), 846–855. https://doi.org/10.1056/NEJMoa1803583.

(129)	Bilsborrow, J. B.; Doherty, E.; Tilstam, P. V.; Bucala, R. Macrophage Migration Inhibitory Factor (MIF) as a Therapeutic Target for Rheumatoid Arthritis and Systemic Lupus Erythematosus. *Expert Opin Ther Targets* **2019**, *23* (9), 733–744. https://doi.org/10.1080/14728222.2019.1656718.

(130)	Calandra, T.; Roger, T. Macrophage Migration Inhibitory Factor: A Regulator of Innate Immunity. *Nat Rev Immunol* **2003**, *3* (10), 791–800. https://doi.org/10.1038/nri1200.

(131)	AstraZeneca. *A Phase II, Multicenter, Open-Label, Dose-Escalation Study to Evaluate Safety and Tolerability of IV or SC Dose of MEDI-545, a Fully Human Monoclonal Antibody Directed Against Interferon Alpha Subtypes, in Japanese Patients Who Have Systemic Lupus Erythematosus (SLE)*; Clinical trial registration NCT01031836; clinicaltrials.gov, 2018. https://clinicaltrials.gov/study/NCT01031836 (accessed 2023-12-31).

(132)	Lee, A. J.; Ashkar, A. A. The Dual Nature of Type I and Type II Interferons. *Front. Immunol.* **2018**, *9*. https://doi.org/10.3389/fimmu.2018.02061.

(133)	Ji, L.; Li, T.; Chen, H.; Yang, Y.; Lu, E.; Liu, J.; Qiao, W.; Chen, H. The Crucial Regulatory Role of Type I Interferon in Inflammatory Diseases. *Cell Biosci* **2023**, *13* (1), 230. https://doi.org/10.1186/s13578-023-01188-z.

(134)	Londe, A. C.; Fernandez-Ruiz, R.; Julio, P. R.; Appenzeller, S.; Niewold, T. B. Type I Interferons in Autoimmunity: Implications in Clinical Phenotypes and Treatment Response. *The Journal of Rheumatology* **2023**, *50* (9), 1103–1113. https://doi.org/10.3899/jrheum.2022-0827.

(135)	Tirado-Rodriguez, B.; Ortega, E.; Segura-Medina, P.; Huerta-Yepez, S. TGF-β: An Important Mediator of Allergic Disease and a Molecule with Dual Activity in Cancer Development. *J Immunol Res* **2014**, *2014*, 318481. https://doi.org/10.1155/2014/318481.

(136)	Kim, B.-G.; Malek, E.; Choi, S. H.; Ignatz-Hoover, J. J.; Driscoll, J. J. Novel Therapies Emerging in Oncology to Target the TGF-β Pathway. *J Hematol Oncol* **2021**, *14* (1), 55. https://doi.org/10.1186/s13045-021-01053-x.

(137)	Kubiczkova, L.; Sedlarikova, L.; Hajek, R.; Sevcikova, S. TGF-β – an Excellent Servant but a Bad Master. *J Transl Med* **2012**, *10* (1), 183. https://doi.org/10.1186/1479-5876-10-183.

(138)	Zarzynska, J. M. Two Faces of TGF-Beta1 in Breast Cancer. *Mediators of Inflammation* **2014**, *2014*, e141747. https://doi.org/10.1155/2014/141747.

(139)	Esquivel-Velázquez, M.; Ostoa-Saloma, P.; Palacios-Arreola, M. I.; Nava-Castro, K. E.; Castro, J. I.; Morales-Montor, J. The Role of Cytokines in Breast Cancer Development and Progression. *J Interferon Cytokine Res* **2015**, *35* (1), 1–16. https://doi.org/10.1089/jir.2014.0026.

(140)	Darif, D.; Hammi, I.; Kihel, A.; El Idrissi Saik, I.; Guessous, F.; Akarid, K. The Pro-Inflammatory Cytokines in COVID-19 Pathogenesis: What Goes Wrong? *Microb Pathog* **2021**, *153*, 104799. https://doi.org/10.1016/j.micpath.2021.104799.

(141)	Mastrandrea, L.; Yu, J.; Behrens, T.; Buchlis, J.; Albini, C.; Fourtner, S.; Quattrin, T. Etanercept Treatment in Children with New-Onset Type 1 Diabetes: Pilot Randomized, Placebo-Controlled, Double-Blind Study. *Diabetes Care* **2009**, *32* (7), 1244–1249. https://doi.org/10.2337/dc09-0054.

(142)	Lu, J.; Liu, J.; Li, L.; Lan, Y.; Liang, Y. Cytokines in Type 1 Diabetes: Mechanisms of Action and Immunotherapeutic Targets. *Clin Transl Immunology* **2020**, *9* (3), e1122. https://doi.org/10.1002/cti2.1122.

(143)	McInnes, I. B.; Schett, G. Cytokines in the Pathogenesis of Rheumatoid Arthritis. *Nat Rev Immunol* **2007**, *7* (6), 429–442. https://doi.org/10.1038/nri2094.

(144)	Mazloom, S. E.; Yan, D.; Hu, J. Z.; Ya, J.; Husni, M. E.; Warren, C. B.; Fernandez, A. P. TNF-α Inhibitor-Induced Psoriasis: A Decade of Experience at the Cleveland Clinic. *J Am Acad Dermatol* **2020**, *83* (6), 1590–1598. https://doi.org/10.1016/j.jaad.2018.12.018.

(145)	Johansson, A.; Hamzah, J.; Payne, C. J.; Ganss, R. Tumor-Targeted TNFα Stabilizes Tumor Vessels and Enhances Active Immunotherapy. *Proceedings of the National Academy of Sciences* **2012**, *109* (20), 7841–7846. https://doi.org/10.1073/pnas.1118296109.

(146)	Holbrook, J.; Lara-Reyna, S.; Jarosz-Griffiths, H.; McDermott, M. F. Tumour Necrosis Factor Signalling in Health and Disease. *F1000Research* **2019**, *8*.

(147)	Szlosarek, P. W.; Balkwill, F. R. Tumour Necrosis Factor α: A Potential Target for the Therapy of Solid Tumours. *The Lancet Oncology* **2003**, *4* (9), 565–573. https://doi.org/10.1016/S1470-2045(03)01196-3.

(148)	Brown, E. R.; Charles, K. A.; Hoare, S. A.; Rye, R. L.; Jodrell, D. I.; Aird, R. E.; Vora, R.; Prabhakar, U.; Nakada, M.; Corringham, R. E.; DeWitte, M.; Sturgeon, C.; Propper, D.; Balkwill, F. R.; Smyth, J. F. A Clinical Study Assessing the Tolerability and Biological Effects of Infliximab, a TNF-α Inhibitor, in Patients with Advanced Cancer. *Annals of Oncology* **2008**, *19* (7), 1340–1346. https://doi.org/10.1093/annonc/mdn054.

(149)	Gerriets, V.; Goyal, A.; Khaddour, K. Tumor Necrosis Factor Inhibitors. In *StatPearls*; StatPearls Publishing: Treasure Island (FL), 2024.

(150)	Ellis, C. R.; Azmat, C. E. Adalimumab. In *StatPearls*; StatPearls Publishing: Treasure Island (FL), 2024.

(151)	Kornbluth, A. Infliximab Approved for Use in Crohn’s Disease: A Report on the FDA GI Advisory Committee Conference. *Inflammatory Bowel Diseases* **1998**, *4* (4), 328–329. https://doi.org/10.1097/00054725-199811000-00014.

(152)	St. Clair, E. W.; van der Heijde, D. M. F. M.; Smolen, J. S.; Maini, R. N.; Bathon, J. M.; Emery, P.; Keystone, E.; Schiff, M.; Kalden, J. R.; Wang, B.; DeWoody, K.; Weiss, R.; Baker, D.; Group, A.-C. S. of P. R. I. for the T. of R. A. of E. O. S. Combination of infliximab and methotrexate therapy for early rheumatoid arthritis: A randomized, controlled trial. *Arthritis & Rheumatism* **2004**, *50* (11), 3432–3443. https://doi.org/10.1002/art.20568.

(153)	Targan, S. R.; Hanauer, S. B.; Deventer, S. J. H. van; Mayer, L.; Present, D. H.; Braakman, T.; DeWoody, K. L.; Schaible, T. F.; Rutgeerts, P. J. A Short-Term Study of Chimeric Monoclonal Antibody cA2 to Tumor Necrosis Factor α for Crohn’s Disease. *New England Journal of Medicine* **1997**, *337* (15), 1029–1036. https://doi.org/10.1056/NEJM199710093371502.

(154)	Leone, G. M.; Mangano, K.; Petralia, M. C.; Nicoletti, F.; Fagone, P. Past, Present and (Foreseeable) Future of Biological Anti-TNF Alpha Therapy. *Journal of Clinical Medicine* **2023**, *12* (4), 1630. https://doi.org/10.3390/jcm12041630.

(155)	Nelson, A. L.; Dhimolea, E.; Reichert, J. M. Development Trends for Human Monoclonal Antibody Therapeutics. *Nat Rev Drug Discov* **2010**, *9* (10), 767–774. https://doi.org/10.1038/nrd3229.

(156)	Bain, B.; Brazil, M. Adalimumab. *Nature Reviews Drug Discovery* **2003**, *2* (9), 693–694. https://doi.org/10.1038/nrd1182.

(157)	Lang, L. FDA Approves Cimzia to Treat Crohn’s Disease. *Gastroenterology* **2008**, *134* (7), 1819. https://doi.org/10.1053/j.gastro.2008.04.034.

(158)	Kuehnemuth, B.; Piseddu, I.; Wiedemann, G. M.; Lauseker, M.; Kuhn, C.; Hofmann, S.; Schmoeckel, E.; Endres, S.; Mayr, D.; Jeschke, U.; Anz, D. CCL1 Is a Major Regulatory T Cell Attracting Factor in Human Breast Cancer. *BMC Cancer* **2018**, *18* (1), 1278. https://doi.org/10.1186/s12885-018-5117-8.

(159)	Das, S.; Sarrou, E.; Podgrabinska, S.; Cassella, M.; Mungamuri, S. K.; Feirt, N.; Gordon, R.; Nagi, C. S.; Wang, Y.; Entenberg, D.; Condeelis, J.; Skobe, M. Tumor Cell Entry into the Lymph Node Is Controlled by CCL1 Chemokine Expressed by Lymph Node Lymphatic Sinuses. *J Exp Med* **2013**, *210* (8), 1509–1528. https://doi.org/10.1084/jem.20111627.

(160)	Campbell, J. R.; McDonald, B. R.; Mesko, P. B.; Siemers, N. O.; Singh, P. B.; Selby, M.; Sproul, T. W.; Korman, A. J.; Vlach, L. M.; Houser, J.; Sambanthamoorthy, S.; Lu, K.; Hatcher, S. V.; Lohre, J.; Jain, R.; Lan, R. Y. Fc-Optimized Anti-CCR8 Antibody Depletes Regulatory T Cells in Human Tumor Models. *Cancer Res* **2021**, *81* (11), 2983–2994. https://doi.org/10.1158/0008-5472.CAN-20-3585.

(161)	Diao, S.; Li, L.; Zhang, J.; Ji, M.; Sun, L.; Shen, W.; Wu, S.; Chen, Z.; Huang, C.; Li, J. Macrophage-Derived CCL1 Targets CCR8 Receptor in Hepatic Stellate Cells to Promote Liver Fibrosis through JAk/STAT Pathway. *Biochemical Pharmacology* **2025**, *237*, 116884. https://doi.org/10.1016/j.bcp.2025.116884.

(162)	Barsheshet, Y.; Wildbaum, G.; Levy, E.; Vitenshtein, A.; Akinseye, C.; Griggs, J.; Lira, S. A.; Karin, N. CCR8+FOXp3+ Treg Cells as Master Drivers of Immune Regulation. *Proceedings of the National Academy of Sciences* **2017**, *114* (23), 6086–6091. https://doi.org/10.1073/pnas.1621280114.

(163)	Xu, M.; Wang, Y.; Xia, R.; Wei, Y.; Wei, X. Role of the CCL2‐CCR2 Signalling Axis in Cancer: Mechanisms and Therapeutic Targeting. *Cell Prolif* **2021**, *54* (10), e13115. https://doi.org/10.1111/cpr.13115.

(164)	Gschwandtner, M.; Derler, R.; Midwood, K. S. More Than Just Attractive: How CCL2 Influences Myeloid Cell Behavior Beyond Chemotaxis. *Front. Immunol.* **2019**, *10*. https://doi.org/10.3389/fimmu.2019.02759.

(165)	Shin, S. Y.; Lee, D. H.; Lee, J.; Choi, C.; Kim, J.-Y.; Nam, J.-S.; Lim, Y.; Lee, Y. H. C-C Motif Chemokine Receptor 1 (CCR1) Is a Target of the EGF-AKT-mTOR-STAT3 Signaling Axis in Breast Cancer Cells. *Oncotarget* **2017**, *8* (55), 94591–94605. https://doi.org/10.18632/oncotarget.21813.

(166)	O’Hayre, M.; Salanga, C. L.; Handel, T. M.; Allen, S. J. Chemokines and Cancer: Migration, Intracellular Signalling and Intercellular Communication in the Microenvironment. *Biochem J* **2008**, *409* (3), 635–649. https://doi.org/10.1042/BJ20071493.

(167)	Mollica Poeta, V.; Massara, M.; Capucetti, A.; Bonecchi, R. Chemokines and Chemokine Receptors: New Targets for Cancer Immunotherapy. *Front. Immunol.* **2019**, *10*. https://doi.org/10.3389/fimmu.2019.00379.

(168)	Yang, Y.-L.; Li, X.-F.; Song, B.; Wu, S.; Wu, Y.-Y.; Huang, C.; Li, J. The Role of CCL3 in the Pathogenesis of Rheumatoid Arthritis. *Rheumatol Ther* **2023**, *10* (4), 793–808. https://doi.org/10.1007/s40744-023-00554-0.

(169)	Wei, C.; Liu, J.; Wu, B.; Shen, T.; Fan, J.; Lin, Y.; Li, K.; Guo, Y.; Shang, Y.; Zhou, B.; Xie, H. Blockage of CCL3 with Neutralizing Antibody Reduces Neuroinflammation and Reverses Alzheimer Disease Phenotypes. *Brain, Behavior, and Immunity* **2025**, *128*, 400–415. https://doi.org/10.1016/j.bbi.2025.04.034.

(170)	Ishida, T.; Ueda, R. CCR4 as a Novel Molecular Target for Immunotherapy of Cancer. *Cancer Sci* **2006**, *97* (11), 1139–1146. https://doi.org/10.1111/j.1349-7006.2006.00307.x.

(171)	Jiao, X.; Nawab, O.; Patel, T.; Kossenkov, A. V.; Halama, N.; Jaeger, D.; Pestell, R. G. Recent Advances Targeting CCR5 for Cancer and Its Role in Immuno-Oncology. *Cancer Res* **2019**, *79* (19), 4801–4807. https://doi.org/10.1158/0008-5472.CAN-19-1167.

(172)	Marques, R. E.; Guabiraba, R.; Russo, R. C.; Teixeira, M. M. Targeting CCL5 in Inflammation. *Expert Opin Ther Targets* **2013**, *17* (12), 1439–1460. https://doi.org/10.1517/14728222.2013.837886.

(173)	Zhang, X.-F.; Zhang, X.-L.; Wang, Y.-J.; Fang, Y.; Li, M.-L.; Liu, X.-Y.; Luo, H.-Y.; Tian, Y. The Regulatory Network of the Chemokine CCL5 in Colorectal Cancer. *Ann Med* *55* (1), 2205168. https://doi.org/10.1080/07853890.2023.2205168.

(174)	Shan, J.; Xu, Y.; Lun, Y. Comprehensive Analysis of the Potential Biological Significance of CCL5 in Pan-Cancer Prognosis and Immunotherapy. *Sci Rep* **2024**, *14* (1), 22138. https://doi.org/10.1038/s41598-024-73251-9.

(175)	Cheng, Y.; Ma, X.; Wei, Y.; Wei, X.-W. Potential Roles and Targeted Therapy of the CXCLs/CXCR2 Axis in Cancer and Inflammatory Diseases. *Biochimica et Biophysica Acta (BBA) - Reviews on Cancer* **2019**, *1871* (2), 289–312. https://doi.org/10.1016/j.bbcan.2019.01.005.

(176)	Hou, C.-H.; Chen, P.-C.; Liu, J.-F. CXCL1 Enhances COX-II Expression in Rheumatoid Arthritis Synovial Fibroblasts by CXCR2, PLC, PKC, and NF-κB Signal Pathway. *International Immunopharmacology* **2023**, *124*, 110909. https://doi.org/10.1016/j.intimp.2023.110909.

(177)	Zhou, C.; Gao, Y.; Ding, P.; Wu, T.; Ji, G. The Role of CXCL Family Members in Different Diseases. *Cell Death Discov.* **2023**, *9* (1), 212. https://doi.org/10.1038/s41420-023-01524-9.

(178)	Zhang, L.; Li, Q.; Zhou, C.; Zhang, Z.; Zhang, J.; Qin, X. Immune-Dysregulated Neutrophils Characterized by Upregulation of CXCL1 May Be a Potential Factor in the Pathogenesis of Abdominal Aortic Aneurysm and Systemic Lupus Erythematosus. *Heliyon* **2023**, *9* (7), e18037. https://doi.org/10.1016/j.heliyon.2023.e18037.

(179)	Korbecki, J.; Maruszewska, A.; Bosiacki, M.; Chlubek, D.; Baranowska-Bosiacka, I. The Potential Importance of CXCL1 in the Physiological State and in Noncancer Diseases of the Cardiovascular System, Respiratory System and Skin. *Int J Mol Sci* **2022**, *24* (1), 205. https://doi.org/10.3390/ijms24010205.

(180)	Kuhne, M.; Preston, B.; Wallace, S.; Chen, S.; Vasudevan, G.; Witte, A.; Cardarelli, P. MDX-1100, a Fully Human Anti-CXCL10 (IP-10) Antibody, Is a High Affinity, Neutralizing Antibody That Has Entered Phase I Clinical Trials for the Treatment of Ulcerative Colitis (UC). (131.20). *J Immunol* **2007**, *178* (1_Supplement), S241. https://doi.org/10.4049/jimmunol.178.Supp.131.20.

(181)	Yellin, M.; Paliienko, I.; Balanescu, A.; Ter-Vartanian, S.; Tseluyko, V.; Xu, L.-A.; Tao, X.; Cardarelli, P. M.; Leblanc, H.; Nichol, G.; Ancuta, C.; Chirieac, R.; Luo, A. A Phase II, Randomized, Double-Blind, Placebo-Controlled Study Evaluating the Efficacy and Safety of MDX-1100, a Fully Human Anti-CXCL10 Monoclonal Antibody, in Combination with Methotrexate in Patients with Rheumatoid Arthritis. *Arthritis Rheum* **2012**, *64* (6), 1730–1739. https://doi.org/10.1002/art.34330.

(182)	Elemam, N. M.; Hannawi, S.; Maghazachi, A. A. Role of Chemokines and Chemokine Receptors in Rheumatoid Arthritis. *Immunotargets Ther* **2020**, *9*, 43–56. https://doi.org/10.2147/ITT.S243636.

(183)	Ma, X.; Norsworthy, K.; Kundu, N.; Rodgers, W. H.; Gimotty, P. A.; Goloubeva, O.; Lipsky, M.; Li, Y.; Holt, D.; Fulton, A. CXCR3 Expression Is Associated with Poor Survival in Breast Cancer and Promotes Metastasis in a Murine Model. *Mol Cancer Ther* **2009**, *8* (3), 490–498. https://doi.org/10.1158/1535-7163.MCT-08-0485.

(184)	Andrews, S. P.; Cox, R. J. Small Molecule CXCR3 Antagonists. *J. Med. Chem.* **2016**, *59* (7), 2894–2917. https://doi.org/10.1021/acs.jmedchem.5b01337.

(185)	Williams, S. A.; Harata-Lee, Y.; Comerford, I.; Anderson, R. L.; Smyth, M. J.; McColl, S. R. Multiple Functions of CXCL12 in a Syngeneic Model of Breast Cancer. *Mol Cancer* **2010**, *9* (1), 250. https://doi.org/10.1186/1476-4598-9-250.

(186)	Marcuzzi, E.; Angioni, R.; Molon, B.; Calì, B. Chemokines and Chemokine Receptors: Orchestrating Tumor Metastasization. *Int J Mol Sci* **2018**, *20* (1), 96. https://doi.org/10.3390/ijms20010096.

(187)	Zlotnik, A.; Burkhardt, A. M.; Homey, B. Homeostatic Chemokine Receptors and Organ-Specific Metastasis. *Nat Rev Immunol* **2011**, *11* (9), 597–606. https://doi.org/10.1038/nri3049.

(188)	Mousavi, A. CXCL12/CXCR4 Signal Transduction in Diseases and Its Molecular Approaches in Targeted-Therapy. *Immunology Letters* **2020**, *217*, 91–115. https://doi.org/10.1016/j.imlet.2019.11.007.

(189)	Cambier, S.; Gouwy, M.; Proost, P. The Chemokines CXCL8 and CXCL12: Molecular and Functional Properties, Role in Disease and Efforts towards Pharmacological Intervention. *Cell Mol Immunol* **2023**, *20* (3), 217–251. https://doi.org/10.1038/s41423-023-00974-6.

(190)	García-Cuesta, E. M.; Santiago, C. A.; Vallejo-Díaz, J.; Juarranz, Y.; Rodríguez-Frade, J. M.; Mellado, M. The Role of the CXCL12/CXCR4/ACKR3 Axis in Autoimmune Diseases. *Front. Endocrinol.* **2019**, *10*. https://doi.org/10.3389/fendo.2019.00585.
